# Biodiversity and niche partitioning in an anaerobic benzene degrading culture

**DOI:** 10.1101/2020.07.17.208124

**Authors:** Chrats Melkonian, Lucas Fillinger, Siavash Atashgahi, Ulisses Nunes da Rocha, Esther Kuiper, Brett Olivier, Martin Braster, Willi Gottstein, Rick Helmus, John Parsons, Hauke Smidt, Marcelle van der Waals, Jan Gerritse, Bernd W Brandt, Douwe Molenaar, Rob van Spanning

## Abstract

A key question in microbial ecology is what the driving forces behind the persistence of large biodiversity in natural environments are. We studied a microbial community with more than 100 different types of species which evolved in a 15-years old bioreactor with benzene as the main carbon and free energy source and nitrate as the electron acceptor. We demonstrate that only a few community members are able to degrade benzene, and that most of the others feed on the metabolic left-overs or on the contents of dead cells making up a food web with different trophic levels. As a result of niche partitioning, a high species richness is maintained and the complexity of a natural community is stabilized in a relatively simple environment. This view highlights the importance of species interactions and interdependencies, which drive microbial community structure and function. These mechanisms may well be conserved across ecosystems.

## Introduction

Microbes typically live in complex and diverse communities [1, 2, 3] the richness of which may in part be determined by environmental conditions like the availability of suitable carbon and free energy sources and the spatial and temporal variability of this environment [4, 5]. Besides environmental conditions, many ecological factors determine the structure and activity of these communities. Such as competition for resources, collaboration by exchange of products, inhibition in a chemical warfare and spatial organization [6]. The extent to which environmental effects and interactions with other organisms can determine species richness in a community is still an open question [4, 7]. Here we approached this question by studying a microbial community living in an anaerobic fixed film bioreactor with benzene as the main sources for carbon and free energy and nitrate as the terminal electron acceptor, thereby creating a relatively simple environment. This microbial community was inoculated from soil samples of a benzene-contaminated industrial site and has been maintained already for 15 years [8], which is enough time to purge a putative initial diversity by mechanisms like competitive exclusion and random fluctuations [9, 10, 11]. Nevertheless, the culture is currently remarkably rich in species [12, 13]. Spatial and temporal heterogeneity as well as the variety of interactions between organisms are then created and maintained by the community itself, aided by mechanisms like wall adherence and patch formation.

Over the years, a biofilm developed on the interior glass wall of the bioreactor, hosting more than 100 different types of species based on 16S rRNA sequencing [13]. It has been hypothesized that these types of microbial community may hold many types of interaction between key players in benzene degradation and other members of the community [14]. A recent study on the same bioreactor revealed high levels of transcripts for an anaerobic benzene carboxylase and a benzoate-coenzyme A ligase produced by *Peptococcaceae* [12]. This finding was in line with other research on benzene degradation where this species was a key player in anaerobic benzene degradation [15, 16]. Benzene is an aromatic hydrocarbon, which occurs in crude oil and petroleum products like fuels. Due to its high toxicity and water solubility, benzene is of major concern as environmental contaminant [17]. Microbes can efficiently open the ring-structure of benzene in the presence of oxygen [18]. However, in hydrocarbon-contaminated subsurface environments, oxygen is rapidly depleted [19, 20] leading to anaerobic benzene degradation, which takes place at a lower rate [19, 21]. Although the biochemistry of anaerobic benzene degradation is relatively well-known [22, 23, 21], the related metabolic pathways are less clear. A comprehensive overview was recently given by Meckenstock et al [24] and it is also a field of active research [25, 26, 27, 28, 8, 15, 29, 14, 30, 31].

The aim of this work was to get a more fundamental understanding of the diversity, structure, metabolic potential and dynamics of the anaerobic benzene-degrading microbial community. We used metagenomics to obtain metagenome-assembled genomes (MAGs), which are assumed to be accurate representations of genomes of individual species [32]. The inferred functions of their genes indicated the potential physiological properties of these species [33]. Metatranscriptomes originating from the biofilm and the liquid phase of the bioreactor were mapped to these MAGs to obtain a measure of their global activity, as well as of the activity of individual genes and pathways. This integrated approach yields a view of the abundances, phenotypes, and activities of these species in these phases [34, 35]. An additional experiment was designed to independently identify the main organisms that drive anaerobic benzene degradation. To this end, we inoculated a series of batch cultures of the 15 years-old microbial community at low cell densities. The samples were sacrificed over time and analyzed for metabolomics on the one hand, and for their community composition based on 16S rRNA gene amplicon sequencing on the other hand. As such we i) identified the drivers for benzene degradation, ii) got insight in the behavior of the culture and its microbial interactions, and iii) propose the relevance of niche partitioning [36] in order to explain the unexpected diversity of species in a bioreactor fed with benzene as main source of carbon and free energy.

## Results

### Diversity and activity of the anaerobic benzene degrading community

We reconstructed 111 MAGs from the metagenomes derived from two samples taken from the biofilm of the culture. From these, 47 high quality MAGs were selected (see Methods, Table S1). The transcriptomes obtained from samples taken from the biofilm and the liquid phase of the culture, were mapped to the predicted genes of all 111 MAGs. Both in the biofilm and the liquid samples the abundance of RNA mapped to MAGs correlated positively with the abundance of the DNA assigned to the MAGs (Fig. S1 A & B, Table S1). The specific RNA abundance per MAG was calculated in biofilm and liquid phase samples as a measure of its transcriptional activity (see Materials and Methods for its definition). In Fig. 1 **a** the specific RNA abundance of biofilm versus liquid revealed a gradient of transcriptional activity (high/intermediate/low), with MAGs 3 and 9 displaying the highest activity, a dozen displaying intermediate activity and the majority displaying low activity. The same conclusion can be drawn when transcriptional activity is calculated as the percentage of transcribed genes per MAG. The majority of the MAGs was found to have a significantly higher specific RNA abundance in samples from the biofilm compared to those from the liquid phase with the exception of four MAGs, including MAG 9, which had the highest overall transcriptional activity in both phases (Fig. S2 & Fig. S3).

**Figure 1.**
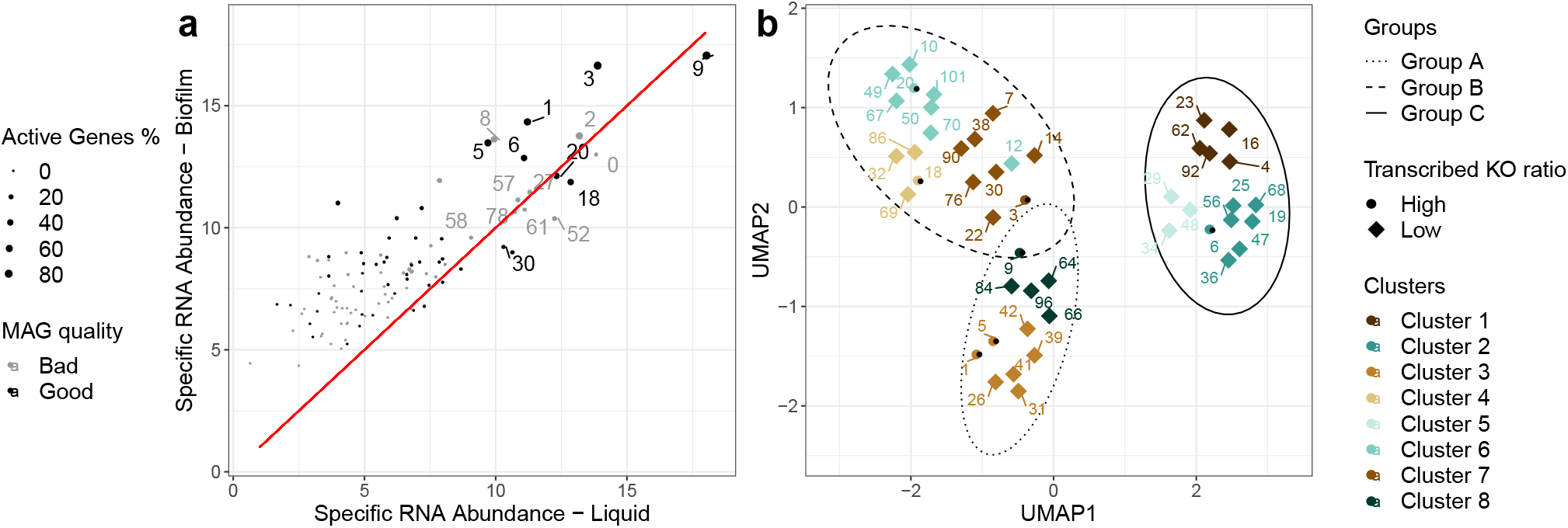
(a) Relationship between specific RNA abundance in biofilm and liquid for all MAGs. The black colored points indicate good/high quality MAGs which were used for further analysis. The relative placement of the points with the red diagonal line reveals on which phase the specific RNA abundance is higher. The size reflects the percentage of identified genes with mapped mRNA. Note that MAG 9 occupies the highest specific RNA abundance in both phases, with MAG 3 coming as second. MAGs 1, 2, 5, 6, 8, 18 and 20 have a medium level of specific RNA abundance. (b) Functional landscape of the selected MAGs. The colors of the points indicate 8 functional clusters of MAGs, while the ellipses further gather them into three groups. The shape of the points indicates the ratio of the transcribed KOs (KEGG Orthology) where the circular points are from the dominant members (as defined in the text) of the community and the rhombus points are from the non-dominant members of the community. For the dimensional reduction and clustering we used Uniform Manifold Approximation and Projection (UMAP) and affinity propagation, respectively, on the binary (presence/absence) KO matrix. Note that the dominant MAGs are distributed across the functional groups.

### Global functional groups and their relation to taxonomic groups

The 47 high quality MAGs were further analyzed by annotating predicted genes using orthology relationships. The presence and absence of genes in the Kyoto Encyclopedia of Genes and Genomes (KEGG) orthology (KO) groups was used to determine potential functional groups of MAGs. Based on these results, the MAGs were divided into 8 clusters (Fig. S3), which were classified into three main groups based on a UMAP dimension reduction. In Fig. 1 **b** the potential functional landscape is visualized. MAGs of group A (Clusters 3 and 8) and group B (Clusters 4, 6 and 7) were more similar in terms of identified function compared to group C (Clusters 1, 2 and 5). Furthermore, members of group C were found to have a significantly higher absolute number of annotated KOs, but not a significantly different ratio of KO annotations (Fig. S5). Only seven out of the 47 selected MAGs were found to have a high specific RNA abundance and high transcribed KO ratio (Fig. S5). Those seven MAGs are distributed over the three groups (Fig. 1 **b**; group A: 9, 1, 5; group B: 18, 20 and 3; group C: 6). We considered these MAGs as putative dominant species and prime candidates to explore their transcription profile in more detail. The overall taxonomic grouping of the 47 selected MAGs corresponded well with the functional grouping. Functional group A has seven members of the Chloroflexi, four of the Actinobacteria and one of the Firmicutes; group B is composed of 11 members of the Bacteroidetes, three of the Gemmatimonadetes, two of the Verrucomicrobia and one member each of the Acidobacteria, of the Armatimonadetes, of the Myxococci and of the Planctomycetes; and group C is composed of 15 members of the Proteobacteria (Fig. S4 & Table S2). An additional 22 MAGs showed significant matches with the Silva 132 database of ribosomal RNA (Table S2). Further details about the taxonomy of the MAGs is presented in the Section S1.10 and Section S2.2.

### Genomic potential of the community members

A substantial part of KOs is shared between the three functional groups (2807 out of 6449 unique KOs), while many other KOs are unique for each group. Moreover, the different clusters within each group showed differences in their KO profiles (Fig. S6). Feature selection revealed 193 KOs, which further mapped into 16 KEGG pathways as discriminators for the three functional groups (Fig. S7 A & B, Fig. S8). We performed a number of targeted searches on potential functionality (genes) and activity (mRNAs) of each of the MAGs. We found that MAGs 3, 6 and 9, as well as those from the Proteobacteria have more and relatively highly expressed genes required for motility and/or adhesion, such as the biosynthesis of flagella and/or pilus systems (Fig. S9 & Fig. S10). Notably, all MAGs from the Chloroflexi expressed relatively high levels of mRNAs encoding peptidases, extracellular solute-binding proteins and specific ABC-type transporters. This property is shared with one of the Bacteroidetes (MAG 18). Members of cluster 2 from the Proteobacteria were found to contain the majority of genes encoding secretion systems, with MAG 36 (*Hydrogenophagus*) having genes for even different types of these systems (Fig. S11). Additionally, we noted relatively high concentrations of mRNAs encoding a type II secretion system from MAG 66 (Actinobacteria), types II and III from MAG 22 (Armatimonadetes) and type VI from MAG 7 (Acidobacteria) and MAG 10 (Ignavibacteria). Genes encoding nitric oxide dismutase (NOD) were found in MAGs 33 (re-assigned on MAG 0), 34 and 71. From those, only MAG 34 passed the quality control and was classified as an unknown member of the *γ*-proteobacteria (Fig. S37). MAG 3 belongs to a member of the Planctomycetes with the highest similarity to *Candidatus* Kuenenia stuttgartiensis. The MAG has the key genes for anaerobic ammonium oxidation (anammox), hydrazine synthase and dehydrogenase. A further description of the metabolic and structural characteristics of the MAGs is available in Section S2.3.

### Metabolism of benzene

In order to characterize the metabolic potential of MAGs, we selected all known pathways involved in anaerobic benzene degradation, central carbon metabolism and nitrate reduction (Section S1.14 for details, Fig. S14 - Fig. S19). A network of these pathways using lumped reactions was created for visualization (Fig. S28 - Fig. S32). Using these custom pathways and corresponding networks, we found only two MAGs from the dominant species with the potential to activate and further degrade benzene anaerobically (primary consumers, Fig. 2). These are MAGs 6 and 9 that were identified as members of *Rhodocyclaceae* and *Peptococcaceae* and belong to groups C and A, respectively (Fig. 2 **c**). In both MAGs we observed a high expression of genes involved in anaerobic benzene degradation, such as UbiD/UbiX-related carboxylases, benzoate-CoA ligase and benzoyl-CoA reductase. Some non-dominant Proteobacteria (MAGs 19, 25, 47 and 68) have the potential to be primary consumers as well (Fig. 2 **d**). Surprisingly, MAGs 19, 36, 47, 56 and 68 showed high expression of genes involved in the aerobic degradation of aromatic compounds, including benzoyl-CoA oxygenase. Moreover, MAGs 19 and 68 had high concentrations of mRNA encoding protocatechuate 4,5-dioxygenase, another oxygen demanding enzyme central in 3,4-dihydroxybenzoate metabolism. For further details on metabolism of benzene we refer to Section S2.5.

**Figure 2.**
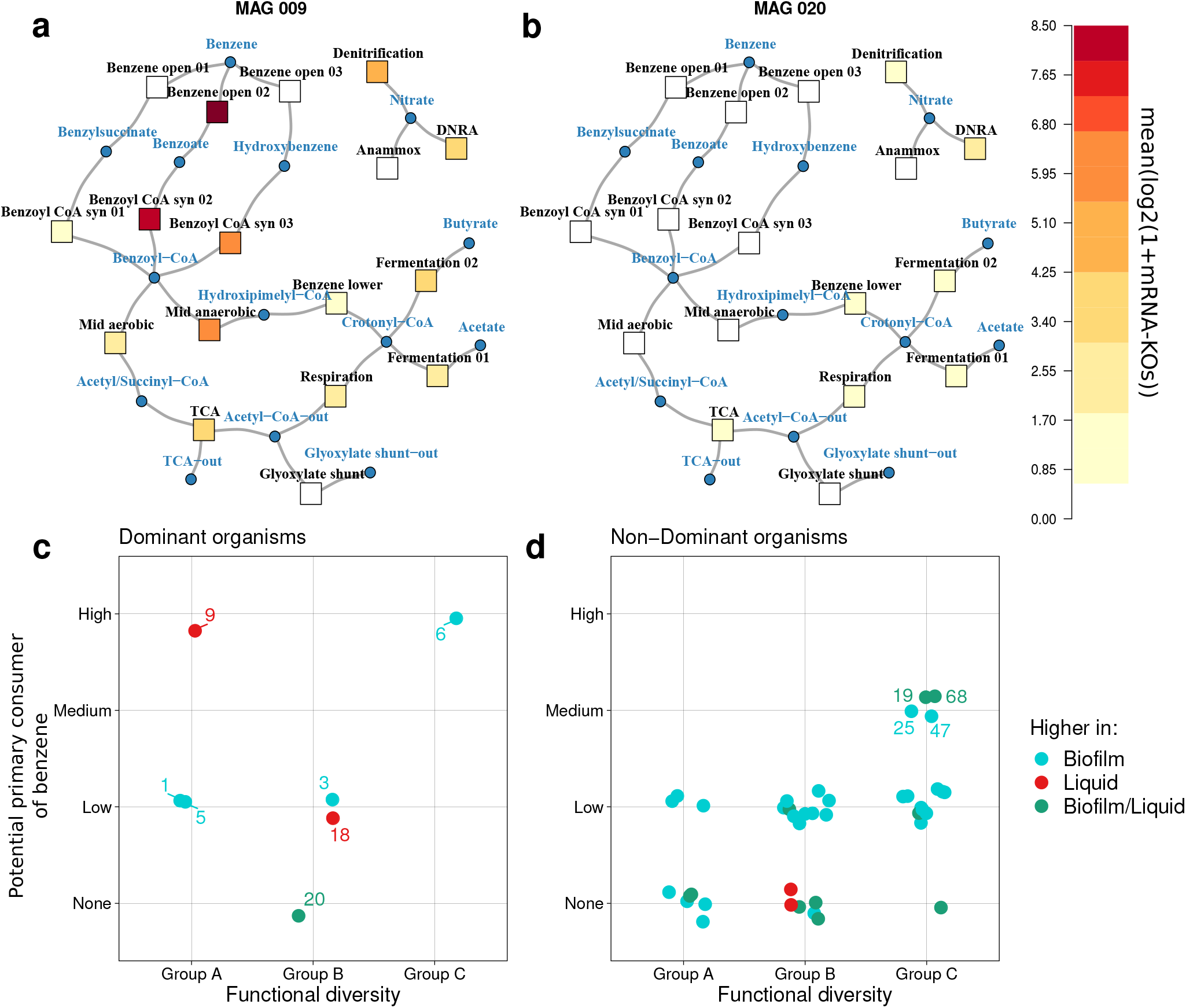
Metabolic heat maps of anaerobic benzene degradation in MAGs 9 (a) and 20 (b), which are dominant members of the community with the highest and lowest potential for benzene metabolism, respectively. The square nodes represent the average transcription of a lumped group of genes that encode enzymes of each branch, and the circular nodes correspond to intermediate compounds. The two lower figures are overviews illustrating the potential of MAGs to be a primary consumer of benzene (the ability to activate and further degrade benzene) based on a custom classification for (c) dominant and (d) non-dominant MAGs. The colors indicate the culture phase in which MAGs show significantly higher specific RNA abundance. The functional diversity is derived from the three groups of Fig. 1. Note that only a few MAGs are assigned as potential primary consumers of benzene, while the majority of the MAGs are not, including dominant MAGs such as 1, 3, 5, 18 and 20. Data are shown jittered for both axis.

### Nitrogen cycling

Nitrate is the main electron acceptor supplied to the benzene-degrading microbial community. The major processes of respiratory electron flow to nitrate and nitrite are i) dissimilatory reduction of nitrate or nitrite to ammonium (DNRA) and ii) sequential reduction of nitrate to dinitrogen gas (denitrification). Most of the 47 MAGs have the potential to perform DNRA (8 MAGs), denitrification (31 MAGs), or both (5 MAGs). Only 3 MAGs, 3, 7 and 36, are unable to do so although they have a gene encoding a nitrate reductase but lack one encoding a nitrite reductase (Section S2.1 for details).

### Succession of communities in batch grown cultures

A succession experiment was performed to investigate the behavior of the community starting from highly diluted cultures with benzene as carbon and free energy source and nitrate as electron acceptor. The cultures were sacrificed at different time intervals up to 34 days after inoculation. Over this time period, individual batch cultures displayed high variability in rates of benzene consumption (Fig. 3 **a**). Some cultures depleted benzene within 22 to 26 days, whereas others had not even started consuming benzene after 34 days. We identified three stages based on the pattern of nitrate consumption (Fig. 3 **a**). Stage 1 is characterized by a lack of benzene consumption (Fig. S33). In stage 2, benzene is consumed till a residual concentration of approximately 0.04 mM, and in stage 3 benzene is depleted down to a residual concentration of around 0.005 mM. Notably, the differences in benzene degradation rates could not be explained by differences in cell densities. As shown in Fig. 3 **b**, cell densities increased quickly after inoculation and then slowly rose up to 20 days. After that time, there were cultures in which benzene had been depleted, but with a range of densities between 10^5^ and 2 × 10^6^ cells per mL. We also observed cultures with high cell density in which benzene degradation did not occur. It may well be that these cultures consumed the acids and vitamins as alternative carbon and free energy source (Fig. 3 **d**). Fig. 3 **c** shows how nitrate consumption, nitrite production and benzene consumption are linked. The data points of the individual cultures form a continuum, suggesting a single deterministic process of benzene degradation. The three stages corresponded to segments of this continuum. Cultures in the early phase of stage 1 consume nitrate without accumulation of nitrite. In a later phase of stage 1, the cultures consumed nitrate and produced nitrite at a stoichiometric 1:1 ratio until a nitrate concentration of about 0.5 mM, indicated by the sloped grey line in Fig. 3 **c**. The later phase corresponds to stages 2 and 3, where nitrate was further consumed whereas the nitrite concentration increased at a lower rate up to 0.7 mM and 0.8 mM, respectively (Fig. S34). Benzene is consumed only in this phase. We measured concentrations for some of the vitamins in the medium samples (*i.e.* those listed in Table S5). However, we could not detect any of the intermediates of benzene metabolism (Table S6 and Table S7) in the medium samples.

**Figure 3.**
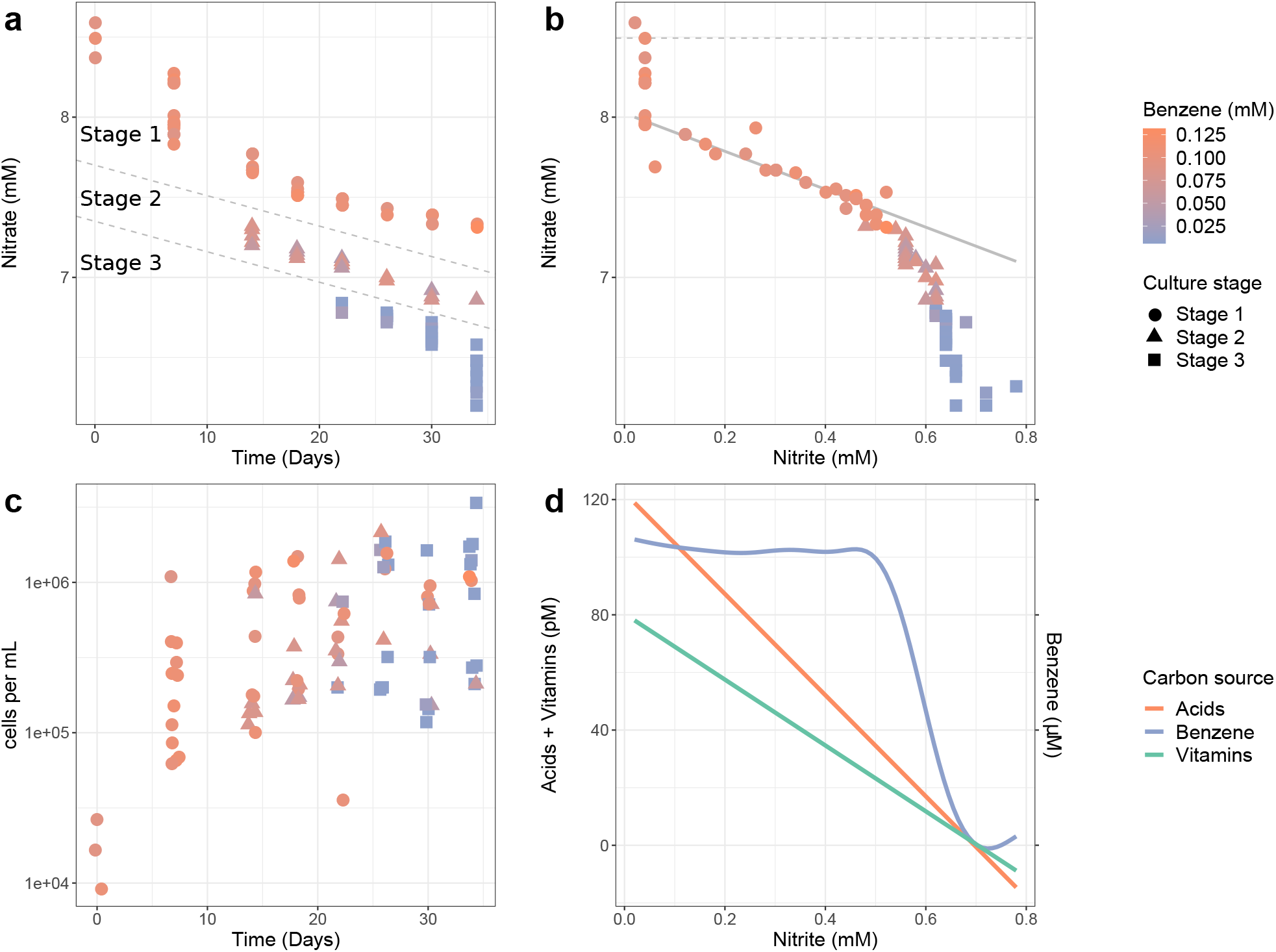
Data analyses of cultures sacrificed at different time points after inoculation. (a) Decrease of the nitrate and benzene concentrations. Stages are separated by the grey lines and explained in the text. (b) Relation between the onset of benzene degradation, nitrate and nitrite levels and culture stage. The dashed line indicates the average of the measured starting concentration of nitrate. The sloped grey line indicates a stoichiometrical (1:1) ratio of consumption and production of nitrate and nitrite. The intercept of this line was chosen by eye. Note that the culture stage and benzene concentration change only above a nitrite concentration of 0.5 mM or a nitrate concentration below 7.3 mM. (c) Correlations of cell density, stage and benzene concentration. Data are shown jittered relative to the time axis. (d) Relation between acids (nicotinic, pantothenic and para-aminobenzoic acids), vitamins (biotin and vitamin B12) and benzene consumption with accompanying nitrite levels (Fig. S35 for detailed view on each carbon source). The functions were fitted with a generalized additive model with integrated smoothness estimation. Acids and vitamins were additives in the original media. Numbers and increments of left and right y-axis are the same. Note the initial consumption of acids followed by vitamins, only after which benzene consumption starts. The sudden drop in the fitted benzene consumption plot coincides with the decrease in nitrite production in stage 2 (Fig. S34 for details).

### Community composition of the batch grown cultures

We also determined the community composition in each of the cultures by amplicon sequencing of the 16S rDNA gene. Reads corresponding to 192 operational taxonomic units (OTUs) were identified and, after correction for 16S rDNA gene copy numbers, expressed as fractions of total corrected reads. Using the measured total cell densities, OTU-specific cell densities were calculated for each sample. We subsequently identified OTUs of which the cell densities correlated with the three stages of culture development (Fig. 4 & Fig. S36). The strongest correlations were observed for OTU624837510 and OTU91680185 whose sequences belong to the genus *Thermincola* and the family of the *Peptococcaceae*, respectively (See Table 1 & Table S9). Not surprisingly, OTU624837510 is found to be identical to the 16S rDNA genes from MAG 9 of the metagenome. Additionally, OTU717462002 correlated well with the progression of stage 2 to 3. It was identified as a member of the *Rhodocyclaceae* and taxonomically identical to MAG 6. We further found significant matches between the 16S sequences of another seven of the selected MAGs and of the apparently corresponding OTUs from the succession experiment (all 100 % identity, Table S2).

**Figure 4.**
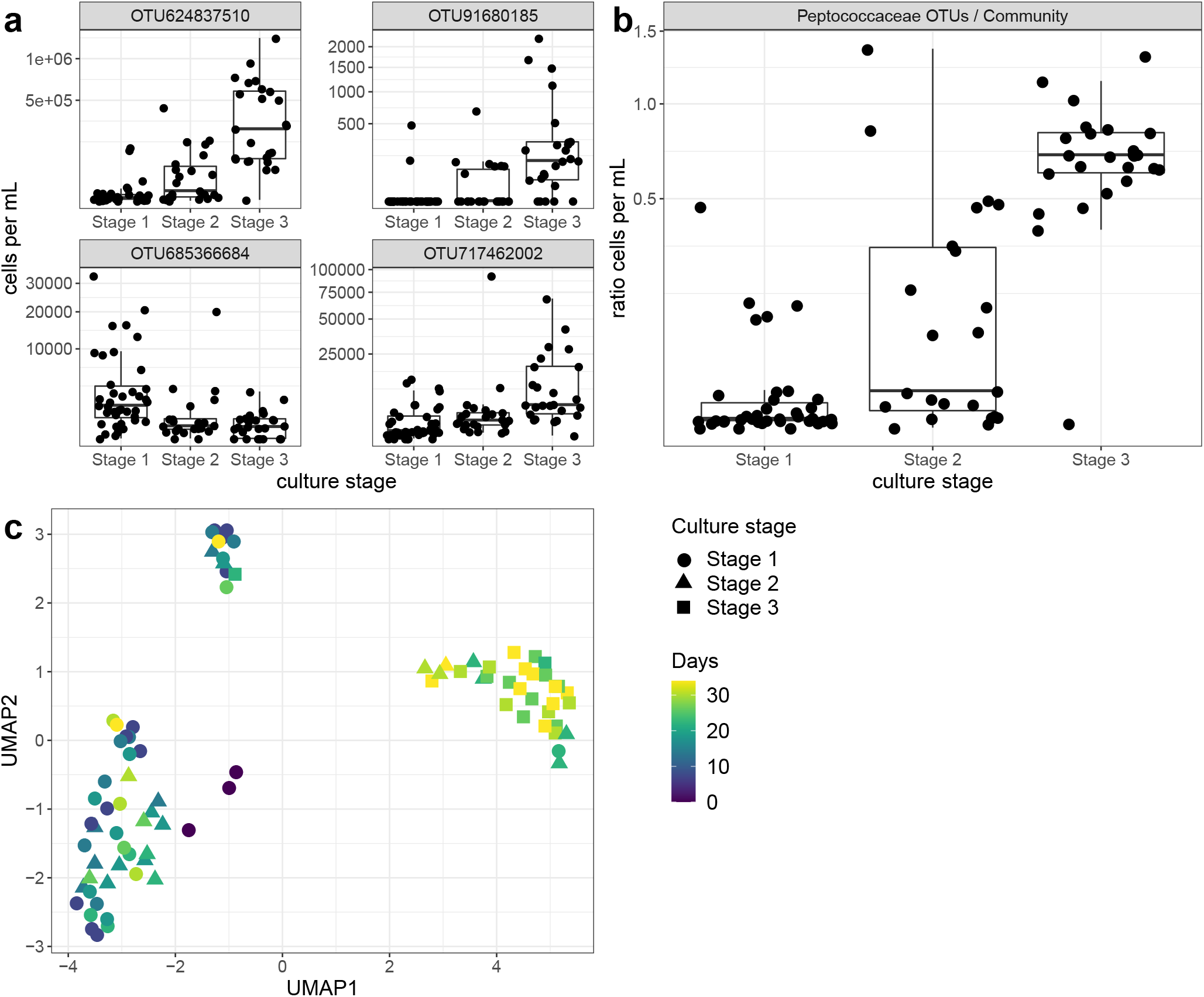
(a) Relationship between culture stage and cell abundance of 4 selected OTUs (by random forest variable importance and PERMANOVA analysis). (b) Relationship between culture stage and ratio of cell abundance between the *Peptococcaceae* OTUs with the rest of the community. The abundance axis has a square-root scale, and points are jittered relative to the culture stage axis. Note that OTU685366684 was correlated with stage 1 and identified as *Pseudomonas aeruginosa* species, a taxonomy not strongly represented in the bioreactor. The succession in time of the community structure in relationship with the culture stage is shown in (c), with usage of UMAP and Bray–Curtis dissimilarity on relative cell abundance matrix. Note that the most diverse community structures are observed in the bottom left cluster of the panel, which include points that belong to stages 1 and 2 throughout all time points of succession (day 0 to 34). A smaller, denser cluster in the top exhibit a similar trend. The cluster in right is composed of cultures from stage 2 and 3 (mid and late phase of the succession). Based on PERMANOVA, the community structures were found significantly different based on the three culture stages (pseudo-*R*2: 0.268 with p-value: 0.001).

**Table 1.**
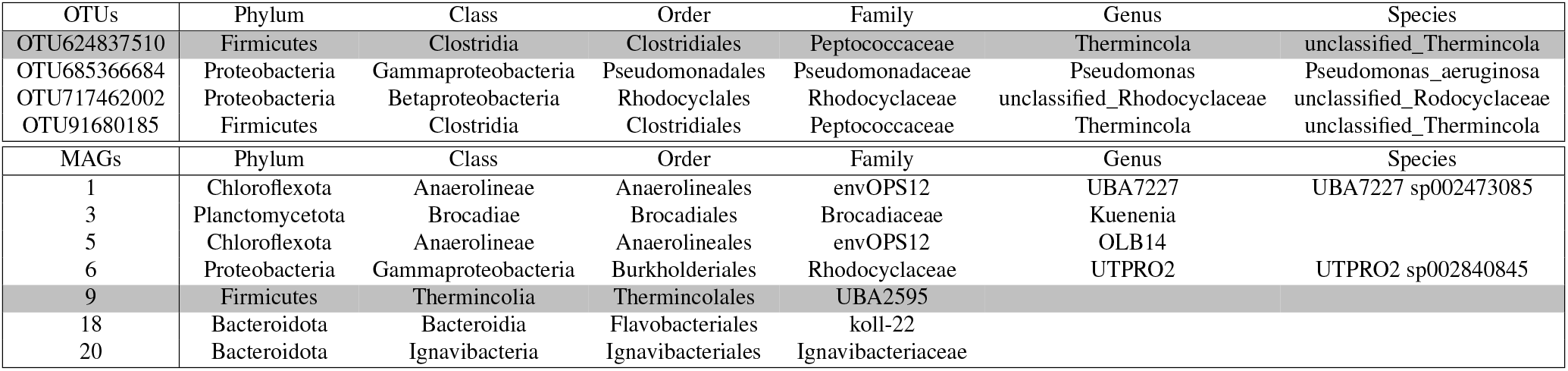
(top) Taxonomy assignment of the strongly correlated OTUs of the succession experiment with cultures stage, selection is based on random forest variable importance and PERMANOVA analysis. (bottom) The dominant MAGs from the microbial community in the bioreactor. Note that the grey backgrounds indicate an identical match between the OTU624837510 sequence from the succession experiment and the identified 16S rDNA sequence in MAG 9 from the bioreactor.

## Discussion

Here we applied a genome-centric transcriptomic approach to obtained insights into the community diversity, structure, function and dynamics of an anaerobic microbial community that developed in a bioreactor during 15 years with benzene as main source of carbon and free energy and nitrate as electron acceptor. In addition, the feed contains some vitamins as minor carbon source and ammonium as minor free energy source. The metabolic potential and gene expression activity of the MAGs that make part of the microbial community suggest niche partitioning [36] with seven representative dominant members, all of which we will discuss below. We hypothesize that they interact with each other via syntrophy, scavenging, predation or even cheating (Fig. 5). Maintenance of and interactions within this relatively complex community may be envisaged in the bioreactor where a continuous flow system is in contact with a biofilm. As such, it may host species that grow at much lower rates than the dilution rate [8, 37].

**Figure 5.**
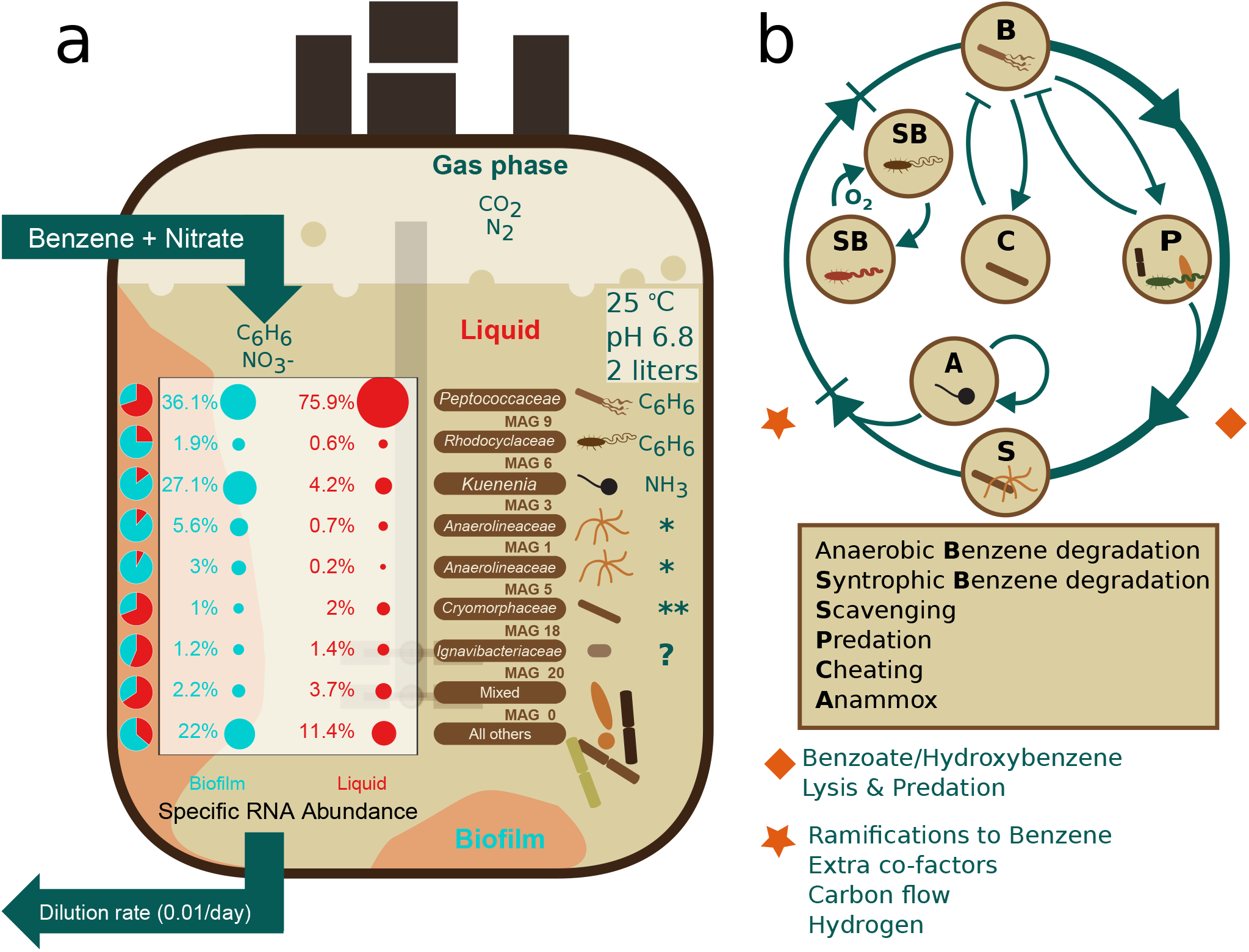
(a) Schematic representation of the anaerobic benzene degrading microbial community in a 15-years old bioreactor. Characteristics of the 7 dominant MAGs with regard to expression levels in biofilm and liquid along with their preferred free energy sources are shown. Pie charts show the ratio of relative specific RNA abundance between biofilm (teal) and liquid (red) with the corresponding values. Values and symbols in teal are derived from samples of the biofilm phase, those in red from samples of the liquid phase. The terms biofilm and liquid phases are explained in the Methods section. Note that most expression in both phases is derived from anaerobic benzene degradation by a member of the *Peptococcaceae*, while that of anaerobic ammonia oxidation by *Candidatus* Kuenenia stuttgartiensis is highest in the biofilm. (* - Lysate) *Anaerolineaceae* potentially using extracellular peptides and proteinaceous polymers derived from lysed cells as carbon and free energy source. (**) *Cryomorphaceae* potentially using secondary metabolites derived from anaerobic benzene degradation. (?) prevalent carbon and free energy source unknown. (b) Schematic representation of dominant niches and their interactions. The outer ring represents the main carbon flow between the primary consumer **B** and the other members of the community. The diamond symbol indicates mechanisms or metabolic left-overs with which **B** feeds the other members of the community and the rhombus symbol indicates what it gets in return. We hypothesize that the community members interact with each other via syntrophy, scavenging, predation or even cheating (See Table S8 for definitions). Those interactions are represented with the arrows that indicate transition and bar-headed arrows to specify inhibition - in or within the outer ring. The arrow that returns back to **A** represents the autotrophic niche, which is occupied by Anammox for free energy transduction and carbon fixation.

Indeed, we found the highest transcription activity of most of the community members in the biofilm, while only four members of the community were shown to be more active in the liquid phase. Our findings show that only two members of the community are directly involved in the degradation of benzene. These are members of the *Rhodocyclaceae* (MAG 6) and of the *Peptococcaceae* (MAG 9), the latter of which is the highest active in expressing genes for benzene metabolism. The member of the *Rhodocyclaceae* was found to be more active in the biofilm as compared to the liquid phase. The member of the *Peptococcaceae* on the other hand is one of the four species that is more active in the liquid phase. In line with this observation is the high expression levels of genes encoding the flagellar system, suggesting that *Peptococcaceae* (MAG 9) is actively moving. It was shown previously that a member of the *Peptococcaceae* was an important member of the anaerobic benzene degrading community [8, 13, 12, 27, 38]. Our data confirm that hypothesis and we postulate that a member of the *Peptococcaceae* is one of the primary benzene consumers with benzoate/hydroxybenzene, benzoyl-CoA and hydroxypimelyl-CoA as intermediates. Some members of the Burkholderiales (MAGs 19, 25, 47 and 68) were also classified as benzene consumers. We noticed that they have the potential to make enzymes for degradation of benzoate and hydroxybenzene, two intermediates of benzene degradation, which may diffuse out of the primary consumers and become available to other members [13]. Burkholderiales were described as important species in enrichment cultures of anaerobic benzene-degrading microcosms and were suggested to use the methylation pathway for anaerobic benzene activation [8, 39, 40].

In our study, we found expression of genes encoding enzymes for benzene degradation that typically combine anaerobic steps with aerobic ones, one of which includes oxygenation to convert benzoyl-CoA to acetyl-CoA and succinyl-CoA. This may seem surprising for an anoxic culture, but the explanation might well be that oxygen is produced locally. Indeed, such an incident production of oxygen has been observed in the anaerobic benzene-degrading microbial community in a previous study [12]. We hypothesize that such production is achieved by certain species that express a nitric oxide dismutase (NOD), and which may come available to other members of the community. The presence of at least two species that can make NODs in our culture is convincing as their relevant protein sequences show the typical characteristics of a NOD (Fig. S37). We envisage the oxygen producers in close proximity of the oxygen consuming members resulting in very low steady state levels of oxygen. We further noticed that a gene encoding protocatechuate 4,5-dioxygenase is highly expressed in two different *β*-proteobacterium (MAGs 19 and 68, successively). The corresponding enzyme catalyzes the oxygen-dependent ring opening of 3,4-dihydroxybenzoate to yield 4-carboxy-2-hydroxymuconate semialdehyde. This observation adds weight to the suggestion that there is local oxygen production in the bioreactor that allows these two species to degrade dihydroxybenzoate aerobically. Moreover, they have also high levels of mRNA encoding benzoyl-CoA oxygenase, which is an oxygen-dependent key enzyme in one of the central branches of benzene degradation. They share this property with yet another three MAGs, 36, 47 and 56.

Other dominant members are exemplified by MAGs 1 and 5 (Chloroflexi), MAGs 18 (Bacteroides) and 20 (Ignavibac-terium). Chloroflexi are detected in a wide range of anaerobic habitats where they are highly abundant and seem to play an important role in formation of flocs and biofilms [41, 42, 43]. In our culture, all Chloroflexi showed significantly higher expression levels in the biofilm as opposed to the liquid phase. Without exception, they all showed relatively high levels of mRNAs encoding extracellular peptidases, solute binding proteins and specific ABC-type transporters. This observation suggests that they grow by cutting extracellular peptides and proteinaceous polymers into smaller molecules that can be transported into the cell to use them as carbon and free energy sources. Other species (MAGs 12 and 49) appear to focus on the synthesis of enzymes for fatty acids uptake and metabolism (see Supplementary information). We therefore speculate that cell lysis of other members of the community may result in the release of these macromolecules in the bioreactor. Such lysis may occur by bacterial members of the community that express at least one of the types II, III or VI secretion systems, the genes and mRNAs of which we identified in some of the MAGs (Fig. S11). Secretion systems induce cell death by the introduction of toxins and other effector molecules in the host cell [44, 45, 46]. Perhaps this type of predation is another strategy for some members of the community to occupy their own niche and to survive as secondary consumers. In other studies, members from the Bacteroidetes were regarded as putative biomass scavengers during syntrophic breakdown of benzene and to express genes for a type VI secretion system [28, 47]. The latter does not seem to be the case for the dominant Bacteroides in our bioreactor (MAG 18) as the annotation pipeline did not yield a secretion system for this MAG. Instead, it rather behaves as the Chloroflexi in the sense that it expresses extracellular peptidases and peptide uptake systems. Moreover, it expresses enzymes for the anaerobic degradation of benzoyl-CoA. The reason for the dominance of MAG 20 is unknown.

Another important niche within the community is occupied by a member of the Planctomycetes (MAG 3), which is an autotrophic organism that uses anammox for free energy transduction and carbon fixation [48]. Genes encoding subunits of the key enzyme of anammox, hydrazine synthase, show a high similarity with those from *Candidatus* Kuenenia stuttgartiensis [48]. As such, it is unique in its choice for the free energy source. This autotrophic species is known to have a low specific growth rate but it is maintained in the bioreactor by nestling in the biofilm as judged by the allocated expression activity. Then we noticed a few other MAGs with the potential to occupy yet other niches using unique types of metabolism. We found upregulated expression of a cluster of genes encoding enzymes for methylamine metabolism in MAG 16, which belongs to the *α*-proteobacteria. Methylamine is a C1-compound that is formed during decomposition of proteins and may be taken up specifically by this methylotrophic organisms to use it as nitrogen, carbon and free energy source [49]. Another *α*-proteobacteria (MAG 4) has high levels of mRNA expressed from a cluster of genes encoding enzymes for formate metabolism.

The succession experiment and downstream correlation analyses not only identified the drivers of the community, but also the assignment of 3 different culture stages with regard to benzene and nitrate consumption rates. Only the cultures that make part of stage 2 or stage 3 reduce nitrate and degrade benzene, in parallel with a lower production rate of nitrite. We postulate that phase 1 is dominated by consumption of the acids and vitamins along with an unbalanced reduction of nitrate into nitrite and further. We then investigated which OTUs are positively correlated with each of the three culture stages. The OTUs belonging to the family of *Peptococcaceae* showed the strongest correlation, followed by an OTU that belongs to the family of *Rhodocyclaceae*. These two are also the most likely primary consumers of benzene in the bioreactor. A partial confirmation of this identity is that the 16S RNA sequence of the most abundant OTU was found to be identical with the one of the *Peptococcaceae* in MAG 9.

Overall, our integrative systems ecology approach revealed that many different niches are occupied in the anoxic benzene degrading bioreactor. This niche partitioning in turn results in a community with a relatively high biodiversity (Fig. 5 **a**). Only a few species appear to metabolize benzene or breakdown products thereof. Yet, a wide range of microorganisms do not seem to feed on benzene but most likely on intermediates of benzene degradation, on biomolecules of lysed cells, or autotrophically with ammonium as free energy source. As a result, most of the community members make up a specialized food web with different trophic levels despite the limited resources. We envisage similar organizations of and interactions within communities across many different ecosystems. Whether or not such communities are driven by key species as we have shown for the community in this study remains to be elucidated.

## Methods

### Microbial community in a fixed film bioreactor

#### Metagenomics sequencing

We selected three samples to use for total DNA high-throughput sequencing. Two of these samples were from brown biofilm (denoted Biofilm 3 and Biofilm 4) collected directly from the bioreactor (Section S1.9 for details) and a third sample (denoted BATCH) was produced by pooling equimolar concentration of DNA from batch cultures prepared in triplicate and inoculated separately. The metagenome libraries using Biofilm 3, Biofilm 4 and BATCH samples were prepared using total DNA extraction for subsequent cluster generation and DNA sequencing using the low-throughput Illumina TruSeq DNA Sample preparation Kit (Illumina, San Diego, California, USA) following instructions from the manufacturer. Later, the libraries were analyzed in a Bioanalyzer 2100 (Agilent Technologies, Santa Clara, CA, USA) and diluted to approximately 8 pM with the addition of 5% PhiX and sequenced using two runs in a Illumina HiSeq 2500 platform (Illumina, San Diego, California, USA) according to the instructions of the manufacturer. The BATCH metagenome was sequenced at the user sequencer facility at the VUmc (Amsterdam, The Netherlands). Biofilm 3 and Biofilm 4 metagenomes were sequenced at GATC-Biotech, Konstanz, Germany.

#### Metagenomics analysis

Adapter cutting and quality assessment were performed by Trim Galore v0.4.0 [50], a wrapper for Cutadapt [51] and FastQC [52]. Bases with a Phred score below 20 were cut off. The paired option was applied with the standard length cut-off of 20 bases, removing a read pair when one or both of the reads is shorter than 20 bases after quality trimming. We used the most diverse and deeply sequenced sample Biofilm 4 for initial assembly. Reads passing the quality assessment were assembled with IDBA_UD v1.1.1 under standard parameters for short reads with the exception that the maximum kmer size was set at 160 [53]. Contigs resulting from the assembly were placed into 111 MAGs with MaxBin v2.1.1 under standard settings and using quality assessed reads from three sequenced samples [54]. MaxBin is dependant on several programs of which the following versions were installed: FragGeneScan v1.20 [55], Bowtie2 v2.2.6 [56] and HMMER3 v3.1b1 [57]. For each MAG, genes were predicted with prodigal v2.6.2 for single genome parameterization [58], followed by functional annotation of the amino acid sequences of eggNOG-mapper v1.0.3 [59]. After a first round of downstream analysis, we refined the MAGs derived from MaxBin using the Anvi’o v6.1 metagenomics workflow [32]. From the refined collection we kept 51 MAGs with completion above 90% and redundancy less than 10%, using the Anvi’o reported scores. Exceptions were made on MAGs 13, 35, 37, 58, and 59, which initially passed the completion/redundancy criteria but later were found highly mixed based on taxonomical classification of their contigs during the manual refinement step. Therefore, they were excluded for further analysis. Differently, MAG 3 (completion: 98.5 %, redundancy: 12.6 %) was included for further analysis, leaving finally 47 high-quality MAGs. The final taxonomy was assigned with GTDB-Tk v1.0.2 [60], as well with alignment search of the rRNAs genes identified in the MAGs (Section S1.10 for details). From the eggNOG annotations, we extracted the KEGG orthology [61] for each MAG and a feature matrix *K* was constructed of dimensions *k* × *m* where *m* is the number of MAGs and *k* is the number of KOs. The entries *K*_*q j*_ are 1 if the KO *q* is present in MAG *j* and 0 otherwise. Overall 6449 unique KOs was found between the selected MAGs. We used the matrix *K* as an measure of indication of potential functionality for the microbial community.

#### Multi-omics analysis

We used in total five out of six metatranscriptomes previously obtained from bioreactor samples and described by our collaborators [12, 13] (Section S1.11 for details). All five samples were gathered by scraping off a confined area of thick parts of the biofilm or thin ones, originally referred to as brown and white biofilm, respectively. The sixth sample was a partial mixture of both liquid and biofilm phases and not included for further analyses. The two brown biofilm samples (B3 and B4) were estimated to contain around 80% biomass, while the white biofilm samples (E1, E2 and E3) contained around 10%. Brown and white biofilm samples were therefore regarded as biofilm and liquid phase, respectively. The mRNA reads were mapped against the predicted genes of each MAG using bowtie v2.3.4.1, samtools v1.2.1 [62], and assigned together with eggNOG annotations. The DNA Abundance of MAGs was calculated by Anvi’o workflow [32]. For RNA Abundance (Eqs. (1) and (2)) we denoted each MAG by *j* ∈ {1 … *m*}, each gene in MAG *j* by *i* ∈ {1 … *n*_*j*_} and each sample by *s* ∈ {1 … 5} where 1 … 2 are samples from the biofilm and 3 … 5 are samples from the liquid phase. Then, we defined *r*_*jis*_ as mRNA read counts per MAG, gene and sample, and *Tr*_*s*_ as the total mRNA read count of a sample. The matrix *RNA Abundance*_*jp*_ resulted from the concatenation of RNA Abundance of the column vectors of biofilm and liquid: *B*_*j*_ and *L*_*j*_ respectively.

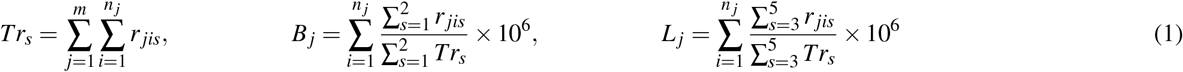

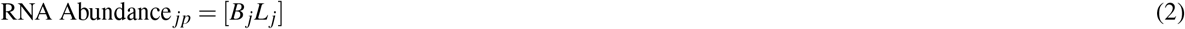

where *p* indicates the phase, biofilm or liquid (*p* ∈ {*b, l*}). To obtain a final measure of potential activity, we normalized the RNA Abundance by the total genome size per MAG into specific RNA Abundance (Eq. (3)). Therefore, if *G*_*j*_ correspond to genome size of a MAG *j* and the overline symbol (like 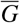) indicates the average taken over all MAGs (similarly 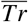 and 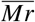 indicate averages below), then:

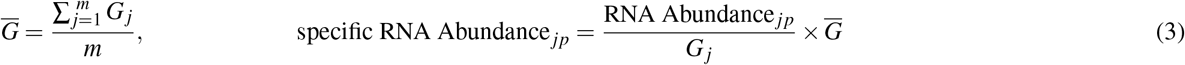

To calculate the percentage of active ORFs ratio, we consider an ORF to be potential active if the average value of the mapped mRNA reads across all samples was equal or bigger than 1. For the differential transcribed analysis of the MAGs we calculated matrix ΔRNA Abundance_*js*_ (Eq. (4)) for all samples. We donated *r* and *Tr* same as above and *Mr* as the total mRNAs per MAG, then:

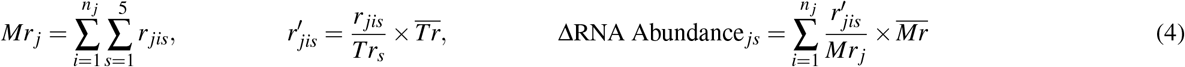

Linear modeling, empirical Bayes moderation and multiple testing with Benjamini-Hochberg method were used to validate the differences at mRNA expression [63, 64]. For the analysis of DNA, RNA, ΔRNA and specific RNA abundance the log2 scale was used. We used R-packages apcluster, uwot and boruta for downstream clustering, dimensionality reduction and importance feature ranking analysis respectively [65, 66, 67, 68, 69, 70] (Section S1.12 for details). Also, R packages KEGGREST were used to get KEGG pathway information, ggplot2, ggrepel for visualization and ShortRead for sequencing processing [71, 72, 73, 74]. Schematic representations and figures was created and polished respectively in Inkscape.

#### Targeted analysis on anaerobic benzene metabolism

To investigate anaerobic benzene degradation we devised the theoretical peripheral and central metabolism based on literature information (Section S1.13, Fig. S12 & Fig. S13 for details). For our analysis we simplified the theoretical scheme by selecting the KEGG reactions and the corresponding KOs. This led to 12 custom-made pathways (Section S1.14 for details). We calculated a specific RNA abundance per gene *i* and MAG *j* as in Eq. (3) (but without averaging over the genes) and added this value to each entry of the *K* matrix (see “Metagenomics analysis” above), when gene *i* was assigned to KO *q*. The resulting matrix we denote by *KT* (Eq. (5)):

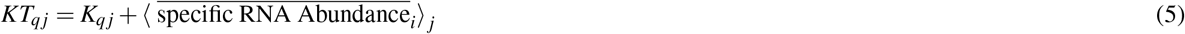

*KT* was used for further analysis on global metabolism, structural components, construct the custom-made pathway selections and visualization of graph/networks [75] (Section S1.14 for details). We used classification scheme on the custom-made pathways (Section S2.4 for details) to assign MAGs as potential primary consumer (High/Medium/Low/None classes) of benzene. Finally, we performed custom searches on functions of interest with blast+ [76]. To do so, we used “makeblastdb” and “blastp” with default parameterization, custom filtering was applied (identity range % > (50 - 80) & alignment length above half of targets sequence length) to obtain the best hits.

### Succession of microbial communities

#### Experimental design

A succession study in batch cultures was carried out to determine important species as drivers for anaerobic benzene degradation. Before performing the succession experiment the benzene-degrading microbial community from the bioreactor was adapted to batch growth as follows. Under anoxic conditions, 0.5 mL of the original bioreactors culture was inoculated in 100 mL serum bottles containing 50 mL of the same phosphate and bicarbonate-buffered medium as used in the bioreactor, and containing 100 *μ*M benzene and excess of nitrate (4.7 mM) as electron acceptor (Section S1.1 for medium composition). After benzene was consumed (Section S1.2 for benzene determination), the medium was spiked again with 100 *μ*M benzene. Once three times benzene completely depleted, the culture was transferred to fresh medium under anoxic conditions (Section S1.1). The succession experiments were performed after three such transfers (over a course of 6 months in total) in 50 mL batch culture. The medium used in the succession experiment was the same medium as in [8]. It consists of a mixture of basal salt medium, phosphate solution, carbonate solution, trace elements solution, vitamin solution, nitrate as electron acceptor and 100 *μ*M benzene. The serum bottles had a 90:10 N_2_:CO_2_ (v/v) atmosphere, and were sealed with Viton stoppers and capped with aluminum crimps. The cultures were incubated at 25°C in the dark (Section S1.1 for details). For the succession experiment, a total of 96 anoxic serum bottles with 100 *μ*M benzene were inoculated with 2 × 10^4^ cells per mL at the same time. We determined cell numbers by separating cell aggregates by gentle sonication (3 × 30 sec at 15 microns of amplitude with 30 sec intervals each), followed by cell staining using SYBR-Green II, and by counting cells in a Accuri C6 Flow Cytometer System (Accuri Cytometers, Ltd., Cambridge, UK) (Section S1.3 for details). The following cultivation and counting controls were performed: (i) no cells added, (ii) no benzene added and (iii) no vitamins added. We prepared five individual bottles for each control (15 in total). The cultures (including the 15 controls) were set up in the 30 mL serum bottles with 20 mL medium. All other conditions kept the same as the adaptation step (mentioned above). A pilot experiment was performed to determine general trends of benzene consumption over time by taking quadruplicate samples under the same conditions of the final succession experiment. Based on the pilot experiment, we inoculated 96 bottles from which we sacrificed 12 bottles each at time points 2h, 7 days, 14 days, 18 days, 22 days, 26 days, 30 days and 34 days after inoculation. The controls were sampled 34 days after the beginning of the experiment. Total DNA was extracted from each of the 96 samples and the 5 controls using a modified CTAB/phenol-chloroform described previously [77] (Section S1.4 for details). DNA from 9 bottles at time point 2h were too low, leaving the DNA from 87 experimental samples for further analyses. The DNA was used for 16S rDNA gene amplicon sequencing including negative controls (i.e. no addition of template DNA to the PCR). We analyzed all samples when sacrificed for benzene by gas chromatography and cell number by flow cytometry (Section S1.2, Section S1.3 & Table S4 for details). We also filtered two times 2 mL of each cell suspension through 0.22 *μ*m nitrocellulose filter membranes (Merck, Darmstadt, Germany). One of the filtrates was used to determine nitrate and nitrite concentrations by capilary electrophoresis (Section S1.5 for details) and targeted metabolomics with the corresponding controls. Targeted metabolomics was used to measure the concentration of the different vitamins used in the medium, including biotin and vitamin B12, and potential intermediates of anaerobic degradation benzene degradation by LC-MS/MS (Section S1.6 & Section S1.7 for details).

### 16S rRNA gene high-throughput sequencing

The 16S rRNA V3-V4 region of the samples from the succession experiment was sequenced using the primers S-D-Bact-0341-b-S-17 and S-D-Bact-0785-a-A-21 [78]. To minimize PCR bias, we performed PCR reactions in triplicate for each sample. Due to low cell biomass each 25 *μ*L reaction contained 0.05 *μ*g of DNA (Section S1.8 for details). Amplicons from all samples were pooled in equimolar concentrations into one composite sample and were paired-end sequenced at the Vrije Universiteit Amsterdam Medical Center (Amsterdam, The Netherlands) on a MiSeq Desktop Sequencer with a 600-cycle MiSeq Reagent Kit v3 (Illumina, San Diego, California, USA) following instructions of the manufacturer. High-throughput sequencing raw data were demultiplexed and processed using a modified version of the Brazilian Microbiome Project 16S rRNA profiling analysis pipeline [79] (Section S1.8 for details). To estimate the total number of species present at the start of the succession experiment we pooled all the sacrificed samples together and identified 192 OTUs that shared more than 97.0% sequence similarity. The OTU abundance was normalized to the total equivalent of cell numbers using the estimated 16S copy number per cell for each OTU. To do so, the ribosomal RNA operon copy number database, hereafter rrnDB [80] was used. These estimates were run using RDP Classifier version 2.10.1 and RDP training set No. 10 incorporating 16S copy number data from the downloadable pan-taxa tables of rrnDB version 4.2.3 [81]. The taxonomic assignment for each OTU (Section S1.8 for details) was used to link the correspondent rrnDB entry. Further, we normalize the data to the equivalent cell number per mL (OTU-specific cell densities) using the flow cytometry cell counts [82]. Selection of important OTUs, which correlated with the three stages of the culture development, was performed with random forest implementation of R-package party [83]. We used default parameters for conditional variable importance, which uses permutation on mean decrease in accuracy. For that, 1000 trees and 10 numbers of random sampling were set. Also, we used R-package vegan implementation of Permutational Multivariate Analysis of Variance (PERMANOVA) on 999 permutations [84, 85, 86].

## Acknowledgements

We thank Frank Bruggeman, Laura Luzia and Bastian Hornung for their significant contribution with discussions and proofreading. Bas Teusink, Evelina Tutucci, Paul Iturbe Espinoza, Philipp Savakis and Huub J. M. Op den Camp for helpful discussions. This study was supported by a grant of BE-Basic-FES funds from the Dutch Ministry of Economic Affairs. The research of CM is supported by a Grand Solution grant from Innovation Fund Denmark (grant no. 6150-00033B), The FoodTranscriptomics project.

## Author contributions statement

CM & DM conceived the methodology, wrote the code, and performed the analysis. Succession experiment and flow cytometry was performed by LF & UNdR. Metabolomics was performed by LF, RH & JP. Nucleic acid analysis was performed by UNdR, EK & BS. BB contributed to the 16 rDNA sequence data processing and analysis. MvdW and JG conceived and conducted the continuous culture experiment. SA and HS performed metatranscriptomic analysis. LF, RvS. & UNdR theoretical reconstruction of metabolic pathways. Initial computational metabolic pathway reconstruction was done by WG & BO. We are grateful to the late Wilfred Röling (Department of Molecular Cell Biology, VU) for his scientific input on experimental design. Unfortunately, he passed away before publication.

CM & RvS wrote the paper. All authors reviewed the manuscript.

## Code & Data availability

Code and processed data/tables can be found in: https://github.com/SystemsBioinformatics/anaerobic-benzene-degrading-culture. Metagenomics shotgun & 16S rRNA gene amplicon sequencing are available in the European Nucleotide Archive under primary accession PRJEB39357. Metagenome-assembled genomes can be found in an Anvi’o result at zenodo open-access repository under http://doi.org/10.5281/zenodo.3939224

## Ethics declarations

The authors declare no competing interests.

## Additional information

Present Address: Willi Gottstein, DSM Delft BV, Delft, Netherlands.

Esther Kuiper, Blaricum, North Holland Province, Netherlands.

