## Supplementary material for "Biodiversity and niche partitioning in an anaerobic benzene degrading culture": Supplemenary_files

### 1 Supplementary Materials and Methods

#### 1.1 Benzene medium

The basal salt medium [69] used in the niche separation experiment was modified as following. Ultra pure water from a Barnstead Nanopure Diamond (Beun De Ronde, The Netherlands) was boiled for 20 minutes. Subsequently the water was flushed with 90% $N_2$ :10% $CO_2$ gas (passed through 0.2 $\mu$m filter) for 20 minutes. Then, we added 1000 mg $KH_2PO_4$, 500 mg $(NH_4)_2SO_4$, 400 mg $MgSO_4$, 100 mg $CaCl_2$ and 800 mg $NaNO_3$ per liter of anoxic water. After the salts were dissolved, we transferred 546 mL of the solution to a 20 min flushed (90% $N_2$ :10% $CO_2$) glass bottle (1 L). After transferred to the glass bottles, the medium was flushed, with the same $N_2$:$CO_2$ mixture as earlier, for 20 min. After flushed, the bottles were closed with Viton stoppers and sealed with crimp-seals. After sealed, the headspace of each bottle was flushed, with the same $N_2$:$CO_2$ mixture as earlier, for other 20 min. Lastly, an overpressure was created by pumping 90% $N_2$ :10% $CO_2$ into each bottle. Next, 2.4 mL Vitamin solution (20 mg Thiamine, 10 mg Riboflavin, 20 mg Nicotinic acid, 50 mg Pyridoxamine, 10 mg Pantothenic acid, 10 mg Vitamin B12, 2 mg Biotin, 10 mg p-Aminobenzoic acid, 5 mg lipoic acid and 5 mg Folic acid dissolved in 100mL sterile anoxic $H_2O$), 2.4 mL DSMZ SL-6 trace element solution (100 mg $ZnSO_4$ 7 $H_2O$, 30 mg $MnCl_2$ 4 $H_2O$, 300 mg $H_3BO_4$, 200 mg $CoCl_2$ 6 $H_2O$, 10 mg $CuCl_2$ 2 $H_2O$, 20 mg $NiCl_2$ 6 $H_2O$ and 30 mg $Na_2MoO_4$ 2 $H_2O$ dissolved in 50mL sterile anoxic $H_2O$), 13.2 mL of 0,5M phosphate stock (3,55 g $Na_2HPO_4$ dissolved in 50mL sterile anoxic $H_2O$) and 36 mL of 1M carbonate stock (4,2 g $NaHCO_3$ dissolved in 50mL sterile anoxic $H_2O$) were added to the basal salt medium. The final pH was adjusted to 7.0. We prepared the 22.55 mM benzene anoxic stock solution (benzene stock solution A) by transferring 30 mL benzene ($\geq$ 99.0%) to a 50 mL serum bottle flushed beforehand with 90% $N_2$ :10% $CO_2$ for 20 min. Subsequently, we flushed benzene with 90% $N_2$:10% $CO_2$ for 30 min. After flushed, the bottle was closed with Viton stoppers and sealed with crimp-seals. After sealed, the headspace of each bottle was flushed for 20 min and incubated overnight at ambient temperature. The next day we opened the serum bottle and flushed the benzene solution with 90% $N_2$:10% $CO_2$ for other 30 minutes. After flushed, the bottle was closed with Viton stoppers and sealed with crimp-seals. Next, the headspace of the benzene stock was flushed for 20 min. Lastly, an overpressure was created by pumping 90% $N_2$:10% $CO_2$ into the bottle. Finally, we diluted the benzene anoxic stock solution A to a 4 mM anoxic benzene stock solution further used to spike the serum bottles to be used in the succession experiment. Prior to adding benzene into the 30mL serum bottles (Sigma-Aldrich, MW, USA) used for the succession experiment, 9.5 ultrapure anoxic water was added to each of them. These bottles were flushed, sealed and had an extra pressure created as described earlier. After the extra pressure was added, all 30 mL serum bottles were autoclaved at 121°C for 20 min. Once autoclaved, 0.5 mL of the 4 mM anoxic benzene stock solution was added to each bottle to a final concentration of 200 $\mu$M in 10 mL of ultrapure anoxic sterile water. We kept an anoxic environment by carrying out inoculations in a Controlled Atmosphere Chamber Model 855-AC (Plas-Labs, Inc. MI, USA) flushed with 50 L per h of 90% $N_2$:10% $CO_2$ gas for 72h. All materials containing plastic used during inoculations were punctured and moved to the chamber 48h prior inoculation. During inoculations, oxygen levels were monitored by an Oxygen Measuring Device GOX 100 (GREISINGER electronic GmbH, Germany). Initially, we inoculated the 600 mL bottles containing basal salt medium, buffers, vitamins and trace elements with approximately 4 x 10 4 cells (see later) of the model benzene consortium per mL. Subsequently, each 30 mL serum bottle containing 10 mL of 200 $\mu$M sterile benzene solution were added of the mixture of benzene microbial consortium cells and basal medium. Orbital shacking of the mixture of cells and basal medium was performed prior inoculation of each serum bottle.

##### 1.2 Benzene determination

Benzene was measured in 150 $\mu$L headspace gas samples, taken through the Viton stoppers of the cultures using a gas-tight Luer-Lok glass syringe and directly injected into a Shimadzu GC-2010 gas chromatograph (Shimadzu, Kyoto, Japan) equipped with a flame ionization detector and an Rtx-1 capillary (30.0 m $\times$ 0.32 mm; Restek, Bellefonte, PA, USA). Helium was used as carrier gas at a flow of 3 mL min-1. The injector and detector temperatures were set to 250°C and 300°C, respectively. The column temperature was kept at 75°C for 2 minutes followed by a gradual increase of 12°C per minute to 100°C. External standards for quantification were prepared in buffered medium in crimp-sealed serum bottles with the same liquid-to-headspace ratio and the medium composition as used for the batch cultures in the succession experiment.

##### 1.3 Determination of cell density

Total cell enumeration in batch cultures was performed using a modified version of a previously described protocol [17]. In a 15 mL polypropylene tube (Greiner, Sigma-Aldrich Co.,Zwijndrecht, The Netherlands) we aliquot 2 mL of cell suspension. Cells were fixed by addition of 50 $\mu$L of formamide to each cell suspension, this suspension was inverted 10 times gently and incubated it 10 min at room temperature. After, we added 4 mL of 10 mM Sodium Pyrophosphated dissolved in filtered and sterilized (autoclave, 121°C for 15 min) Ultrapure water from a Barnstead Nanopure Diamond (Beun De Rond, The Netherlands). Aggregates of cells were disassembled by sonication by placing the polypropylene tubes in a plastic glass filled with water, no more than four at a time. We optimized sonication conditions by varying the amplitude, time and interval of sonication using a SONIPREP 150 Ultrasonic Disintegrator (MSE Ltd., London, UK). Optimal sonication was determined by light microscopy of cell suspensions prior and after sonication and consisted of 3x sonication with 30 sec interval, when tubes were softly inverted, using 15 $\mu$m of amplitude. We stained cells by adding of 0.5 $\mu$L of SYBR Green II (Thermo Fisher Scientific Inc., Reinach, Switzerland), inverting tubes ten times and incubating them for 5 min in the dark. Prior flow cytometry, we fixed, sonicated and stained 2 mL of the same batch of non-inoculated benzene medium used in the succession experiments to measure the presence of aggregates that could be stained by SYBR-Green II. Cell densities, expressed as numbers of cells per mL, were determined by running 200 $\mu$L of cells suspensions and non-inoculated benzene medium as a controu in a Accuri C6 Flow Cytometer System (Accuri Ctometers, Ltd., Cambs, UK) using medium velocity setting and Filter FL1-A (specific for SYBR-Green). Cell densities of samples were used to calculate OTU abundances (see section Section 1.8).

##### 1.4 Nucleic acids extraction

We extracted nucleic acids for amplicon sequencing (samples from the succession experiment), metagenomics and metatranscriptomics. Prior sampling the 115 samples during the succession experiment we homogenized each serum bottle by flipping them five times before collection of 1 mL of microbial suspension were individually added to a 2mL Lysing Matrix E tube (MP Biomedicals, Solon, OH, USA) and snap-freeze on liquid nitrogen. The 1 mL aliquots were Samples were stored at -80 o C prior nucleic acid extraction. Further, we collected samples for metagenomics and metatranscriptomics directly from the original bioreactor [69]. Liquid samples were taken in triplicate from bioreactor outlet. To sample biofilm samples, defined areas of biofilm attached to the bioreactor glass wall were scraping off under a constant $N_2/CO_2$ (80/20%) flow. Subsequently, the liquid phase in the vessel was stirred for 5 minutes at 200 rpm to dislodge the biofilm aggregates followed by liquid phase sampling using a 60 ml syringe via a sampling pipe and viton tubing (Rubber BV, Hilversum, The Netherlands). We added 750 $\mu$L of formamide to each sample and centrifuged the samples at 4000 $\times$ g at 4°C for 20 min in a Hettich® ROTINA 420/420R centrifuge (manufacturer information). The 6 samples, approximately 1 mL each, were added to 2mL Lysing Matrix E tube (MP Biomedicals, Solon, OH, USA). The samples were immediately stored at -80°C until further analyses. Co-extraction of DNA and RNA was performed using a modified CTAB/phenol-chloroform described previously [54]. We prepared crude nucleic acid extractions by adding 0.5 mL phenol:chloroform:isoamyl alcohol (25:24:1) (Sigma-Aldrich, St. Louis, MO, USA) to each 2mL Lysing Matrix E tube containing 1 mL microbial suspension, followed by addition of 0.5 mL of CTAB buffer (5% CTAB, 0.25M phosphate buffer pH 8.0, 0.3M NaCl). The samples were beaten at 5.5 m/s for 30 s in a The FastPrep® FP120 cell disrupter (Thermo Savant, CA, USA), and centrifuged at 16K g for 5 min at 4°C in a microcentrifuge CR 3i (Jouan S.A., France). The aqueous phase was transferred to MaxTract High Density 2 mL tubes (Qiagen Inc, Valencia, CA, USA). A second round of extraction with 0.5 ml CTAB buffer, and beating was performed. An equal volume of chloroform was added to each MaxTract High Density tube and centrifuged at 16K g for 5 min at 4°C. The aqueous phase was transferred to a new 2 mL Eppendorf tube and the nucleic acids were precipitated over night at 4 o C with 2 volumes of 30% (w/v) polyethylene glycol 6000 and 1.6 M NaCl. The crude nucleic acid pellets were resuspended in 30 $\mu$L of DEPC-treated water, and stored at -80°C. Purification of total DNA from the 115 samples of the succession experiment was achieved using a QIAamp DNA Mini Kit (Qiagen Inc, Valencia, CA, USA) following manufacturer's instructions. Further, we co-purified the DNA and RNA from the 6 samples originated from bioreactor using a AllPrep DNA/RNA Mini kit (Qiagen Inc, Valencia, CA, USA) following manufacturer's instructions.

##### 1.5 Determination of nitrate and nitrite

We determined the concentration of nitrate and nitrite using capillary electrophoresis ([30]). In short, samples were centrifuged at 13,000 $\times$ g for 15 min at 4°C. The supernatants were transferred to tubes containing a 0.22 - $\mu$m microspin filter (Ultrafree-MC, Millipore, Bedford, MA, USA) and centrifuged at 12,000xg for 5 min at 4°C. Filtered supernatants were stored at -80°C. The capillary electrophoresis was performed on a Beckman P/ACE™ MDQ (Beckman Coulter, Brea, CA, USA) system at 25 °C with UV detection at 214 nm, capillary length of 50 cm, and separation at 25 kV in reverse mode. Run buffers were derived from the CEofix™ Anions 2 kit (Analis, Suarlée, Belgium). Potassium bromate was used as the internal standard in all samples.

#### 1.6 Vitamin analysis

Vitamins were analyzed by LC-MS/MS using a Prominence XR HPLC system (Shimadzu, Den Bosch, The Netherlands) coupled to a QTRAP 4000 tandem mass spectrometer (AB SCIEX, Framingham, MA, USA). Chromatographic separation was achieved with a Kinetex core-shell column containing a biphenyl stationary phase (2.6 $\mu$m particle size, 100 Å pore size, 100 $\times$ 2.1 mm; Phenomenex, Utrecht, The Netherlands) with an injection volume of 3 $\mu$L, a flow rate of 0.2 mL min-1, and a column temperature of 30°C. Gradient elution with eluent A (0.01% v/v acetic acid in $H_2O$) and eluent B (100% methanol) was as follows: 0-3 min (0% B); 3-8 min (linear increase to 100% B); 8-11 min (100% B). The equilibration time between injections was set to 7 min. Ionization of LC effluent was achieved by electrospray ionization (ESI), switching between positive (0-6.9 min, dwell time 40 ms; 7.4-8.5 min; dwell time 50 ms) and negative mode (6.9-7.4 min, dwell time 50 ms; 8.5-11.6 min, dwell time 50 ms) during the run, with $N_2$ as ionization and collision gas. Ionization settings were as follows: ionization temperature 400°C; ionization voltage 4,000 V (-4,000 V in negative mode); curtain gas 10; nebulizer gas 40; heater gas 50; collision gas 6 (all arbitrary units). Matrix-matched standard solutions for external calibration curves were prepared in culture medium at concentrations from 0.002-0.512 ppm. Quantification was based on chromatographic peak areas. Data acquisition and analysis was done with the AB SCIEX Analyst 1.5.1 software. Compound-specific measurement parameters are given in Table 5. Using this method we were able to measure seven out of the ten vitamins added to the medium at the beginning of the incubation. Vitamins that we not measurable with this method were: thiamine, folic acid, and pyridoxamine. Although we could detect riboflavin, concentrations of the samples from the beginning of the incubation were lower than the actual concentrations added to the medium, and additionally, we also saw degradation of riboflavin in the sterile controls at the end of the incubation. Therefore, the observed decrease in riboflavin concentrations in the cultures might be due to abiotic degradation.

#### 1.7 Metabolite analysis

Metabolite analysis by LC-MS/MS was carried out on a Nexera UHPLC system (Shimadzu, Den Bosch, The Netherlands) coupled to a high-resolution quadrupole time-of-flight mass spectrometer (Q-TOF; maXis 4G, Bruker Daltonics, Wormer, The Netherlands). Compounds were separated on a C18 stationary phase column (1.7 $\mu$m particle size, 150 $\times$ 2.1 mm; ACQUITY UPLC CSH C18, Waters, Etten-Leur, The Netherlands) preceded by a guard column (1.7 $\mu$m particle size, 5 $\times$ 2.1 mm; ACQUITY UPLC CSH C18, Waters) with an injection volume of 20 $\mu$L, a flow rate of 0.2 mL min-1, and column temperature of 30°C. Gradients of eluent A (0.01% v/v acetic acid in 10% v/v methanol) and eluent B (0.01% v/v acetic acid in 100% methanol) were as follows: 0-1 min (5% B); 1-15 min (linear increase to 100% B); 15-18 min (100% B). Compound ionization was carried out by ESI operating in negative mode using $N_2$ as ionization gas with the following settings: capillary voltage 3,500 V; end plate offset 500 V; nebulizer gas pressure ($N_2$) 1 bar; dry gas ($N_2$) 8 L min-1; dry temperature 200°C. Settings for MS analysis were: funnel radio frequency (RF) 200 Vpp (voltage point to point); multipole RF 200 Vpp; collision cell RF 200 Vpp; transfer time 40 $\mu$s; prepulse storage 5 $\mu$s. Internal mass calibration was performed automatically during every measurement by loop injection of 20 $\mu$L of a 2 mM sodium acetate solution in 1:1 v/v ultrapure water-isopropanol [12]. Data acquisition and analysis were done using the Bruker Daltonics software suites Compass 2.7 and DataAnalysis 4.2, respectively. Data-independent target analysis of compounds for which pure standards were available Table 6 was carried out in broadband collision induced dissociation (bbCID) MS/MS mode with a collision energy of 17 eV and an acquisition rate of 1 Hz. Compound identification was based on chromatographic retention time, a mass accuracy threshold ($\leq$ 5 ppm), and an isotopic pattern fit threshold ($\leq$ 50 mSigma) [12]. To confirm the identities of detected compounds in the samples and to test for matrix effects, samples were spiked with standards to a final concentration of 0.15 ppm. A subset of samples (three biological replicates sampled at 0, 18, 28, 32, and 36 days after inoculation) were selected for data-dependent suspect analysis carried out in auto-MS/MS mode by screening MS spectra for masses of putative metabolites and automatically selecting the found masses for MS/MS analysis Table 7. Mass spectra for precursor ions were recorded in a range of 35-800 m/z (excluding 58.91-59.11 m/z corresponding to acetic acid in the eluent) with a spectra rate frequency of 2.5 Hz. Collision energies were increased with increasing masses as follows: 35-100 m/z: 15 V; 100-500 m/z: 25 V; 500-1000 m/z: 50 V. Isolation widths for precursor ions were set to 8 m/z. Acquisition cycle time was set to 2 seconds.

##### 1.8 16S rDNA amplicon sequencing and processing

We performed triplicate PCR reactions to each sample to minimize PCR bias. Each twenty-five $\mu$L reaction consisted of 0.05 $\mu$g of DNA, 0.5 $\mu$L of Phusion Green Hot Start II High-Fidelity Dna Polymerase (Thermo Fisher Scientific, Sweden), 5.0 $\mu$L of 5x Phusion Green HF buffer $MgCl_2$, 2.3 $\mu$L of 25 mM $MgCl_2$ stock solution, 5.0 $\mu$L of 10 $\mu$M primer mix (1:1) and 0.5 $\mu$L of 10 mM nucleotide mix. The thermal cycling protocol was 98°C form 30 sec, 30 cycles of 98°C for 10 sec, 55°C for 30 sec, 72°C for 30 sec and a final 10-min extension at 72°C. We targeted the V3-V4 region of the 16S rRNA gene, primer pairs were: S-D-Bact-0341-b-S-17, 5'-CCTACGGGNGGCWGCAG-3 [29], and the V4 reverse primer S-D-Bact-0785-a-A-21, 5'-GACTACHVGGGTATCTAATCC-3 [29]. The primers were dual barcoded and were compatible with Illumina sequencing platforms as described previouslsy [11]. All amplicons were run in 0.9% (w/v) agarose gels and bands containing expected size were excised from the gel and purified using QIAquick Gel Extraction kit (QIAGEN GmbH, Hilden, Germany). Purified amplicons from each triplicate reaction were pooled together and further quantified using PicoGreen dsDNA assay (Invitrogen). High-throughput sequencing raw data were demultiplexed and processed using a modified version of the Brazilian Microbiome Project 16S profiling analysis pipeline [51]. Quality trimming was done according with the following parameters: quality score > 30, sequence length > 285, no maximum ambiguous bases and no mismatched bases in the primer. Sequences belonging to different samples were demultiplexed using bcl2fastq software version 1.8.4 (Illumina), primers were trimmed using Cutadapt [38] and paired-end reads were joined using PANDAseq [39]. Metadata and demultiplexed sampels were merged using add_qiime_labels.py [11] and sequence headers were changed using bmp-Qiime2Uparse.pl [51]. UPARSE was used to dereplicate, discard OTUs detected less than 4 times and OTU cluster at 97% similarity [19]. We filtered chimeras by reference database search using UCHIME algorithm [20] and the SSU rRNA gene SILVA database release 123 [52]. OTU taxonomy was assigned using the UCLUST algorithm [19] on QIIME [11] using SILVA compatible taxonomy mapping files [52] and aligned using SINA [50]. Taxonomy was manually curated and refined up to genus level based on 95% similarity of reference sequences. The reference tree was calculated using FastTree 2 [49]. We generated a BIOM file using make_otu_table.py on

QIIME [11]. Further, because of the low cell numbers we decided to start the PCR reactions with a low yield DNA (0.05 $\mu$g of DNA per reaction). To avoid the presence of OTUs of external origin in the experimental samples, we added two blank PCR reactions per plate. Every OTU found in the blank PCR reactions was removed from the BIOM file prior to the community analysis.

##### 1.9 Bioreactor Sampling

Liquid samples were taken in triplicate from bioreactor outlet. To sample biofilm, defined areas o attached to the bioreactor glass wall were scraping off under a constant $N_2/CO_2$ (80/20%) flow. Subsequently, the liquid phase in the vessel was stirred for 5 minutes at 200 rpm to dislodge the biofilm aggregates followed by liquid phase sampling using a 60 ml syringe via a sampling pipe and viton tubing (Rubber BV, Hilversum, The Netherlands). The samples were immediately stored at -80°C until further analyses. A phenol-chloroform sumultaneous extraction and purification of DNA and RNA was done as described above.

##### 1.10 Metagenomics analysis, MAG selection and taxonomy assignment

For the Anvi'o v6.1 metagenomics workflow, we used the previously assembled contings to create a corresponding Anvi'o contig databases with the default settings, the trimmed sequences of Biofilm 3 & 4 were mapped to the assembled contings with Bowtie2 v2.3.4.1 and samtools v1.2 [35]. The pipeline uses underline HMMER3 v3.1.b2 [18] for sequence search, NCBI's COGs [60] to annotate genes with function and incorporates Kaiju v1.7.3 [42] web server results to assign taxonomy on the contigs bases on the NCBI's non-redundant protein database (for Bacteria, Archaea, Viruses and database last update: 2017-05-16). During manual refinement we kept the major group of contigs with matched taxonomy on phylum level. We placed all the excluded contings together with all the non-clustered contings from MaxBin into a mixed collection indicated as MAG 000. The refined MAGs were functional annotated eggNOG-mapper v2 [26] with default setting and minimum % of query and subject coverage to 50. On the first round of analysis (on the MAG collection derived from MaxBin2), the quality assessment performed with CheckM tool [47]. The MAGs with completion above 90% and redundancy less than 10% were considered good quality for further analysis. The initial taxonomy assignments were manually assigned by combining multiple approaches, In more detail, we apply the multi-metagenome pipeline [2]and MEGANs [25] LCA algorithm. Also, we used the JSpeciesWS webservie [55]and we reconstructed the phylogeny with PhyloPhlAn [57] on 5847 out of 22055 selected complete bacteria genomes (based on previous taxonomies), plus the good quality MAGs. On the second round on the refined MAG collection we kept the same quality thresholds, but applied on the Anvi'o reported scores. Exception were made on MAGs 13, 35, 37, 58 and 59, because they found highly mixed during manual refinement and we excluded them and on MAG 3 (completion: 98.5 %, redundancy: 12.6 %) which we include. The final taxonomy assignments were done with GTDB-Tk v1.0.2 [14]. Finally, custom blast search with post-filtering in R (sequence identity > 97% and bit score > 200) was performed between a local ribosomal RNA sequence database (SILVA 132 SSURef Nr99)[52] and our metagenomic MAGs (see overview, Supplementary table 2). Similar custom blast search was performed between the OTUs from the succession experiment and the metagenomics MAGs.

#### 1.11 Metatranscriptomes sequencing and analysis

Quality assessment and trimming of Trueseq adapters were performed with Trim Galore v0.6.0 [4], which uses Cutadapt v2.3. [38] and FastQC [3], with a minimum sequence length of 40 bp and a minimum quality of 30 on both ends of the read and as mean quality. All reads with non-IUPAC characters were discarded as were all reads containing more than three Ns. We removed ribosomal RNA reads using SortMeRNA v2.1 [31] and all included databases as indicated by developers.

#### 1.12 Multi-omics analysis details

RNA and DNA were co-extracted from all six biofilm samples to obtain concurrent metagenome and metatranscriptome data sets. Clustering analysis is performed using affinity propagation on resulting Pearson correlation matrix of $K$. A general work-flow to assess the most suitable number of clusters is started with high exemplar preferences values, which led to a very large number of clusters. Application of agglomerative clustering on the resulting affinity propagation clusters using the R-package apcluster [7]. Therefore, a cutoff manually decided and affinity propagation rerun repeatedly to achieve the desirable number of clusters. Furthermore, dimensional reduction with Uniform Manifold Approximation and Projection (UMAP) [6] was performed to visualize the matrix $K$ and the previously obtained cluster assignments. The R package Boruta [33] was used to obtain a reliable ranking of feature importance and to select only discriminative features (in our case KOs) for the classification task derived from the previous clustering analysis. The algorithm is a wrapper around Random Forest [10] that performs randomization tests. Default parameterization was used except the maximal number runs, which was increased to 2000.

#### 1.13 Theoretical scheme of anaerobic benzene degradation

Fig. 12 illustrates the peripheral degradation pathways up to the level of benzoyl CoA intermediates. Dashed lines indicate putative reactions. Compounds surrounded by dashed rectangles indicate hypothesized intermediates. Cofactors and conversion products thereof marked with a question mark indicate hypothesized compounds. Reactions for which enzymes and/or genes have been identified are labeled with numbered dots. Compounds: (12) benzene; (13) toluene; (14) (R)-benzylsuccinate; (15) (R)-benzylsuccinyl-CoA; (16) (E)-phenylitaconyl-CoA; (17) 2-[hydroxy(phenyl)methyl]-succinyl-CoA; (18) benzoylsuccinyl-CoA; (19) benzoate; (20) phenol; (21) phenylphosphate; (22) 4-hydroxybenzoate; (23) 4-hydroxybenzoyl-CoA; (24) benzoyl-CoA; (Fum) fumarate; (SucCoA) succinyl-CoA; (Suc) succinate; (HSCoA) conenzyme A; (AcCoA) acetyl-CoA; (MeTHF) methyl-tetrahydrofolate; (SAM) (S)-adenosyl-methionine; (THF) tetrahydrofolate; (SAH) (S)-adenosyl-homocysteine; (Pi) inorganic phosphate; (Fd) ferredoxin. Enzymes/genes: (R6) benzylsuccinate synthase (BssABCD); (R7) succinyl-CoA:(R)-benzylsuccinate CoA-transferase (BbsEF); (R8) (R)-benzylsuccinyl-CoA dehydrogenase (BbsG); (R9) phenylitaconyl-CoA hydratase (BbsH); (R10) 2-[hydroxy(phenyl)methyl]-succinyl-CoA dehydrogenase (BbsCD); (R11) benzoylsuccinyl-CoA thiolase (BbsAB) [names for (R4-11) identical for A. aromaticum EbN1, Azoarcus spp., Thauera spp., Magnetospirillum spp., and G. metallireducens]; (R12) putative benzene carboxylase (AbcDA, anaerobic enrichment cultures [34, 36]; AbcA, F. palcidus); (R13) benzoate-CoA ligase (*bclA*, A. aromaticum EbN1, Thauera spp., Magnetospirillum spp.; *bzdA*, Azoarcus spp.; *bamY*, G.

metallireducens, S. aromaticivorans); (R13') succinyl-CoA:benzoate CoA transferase (*bct*, G. metallireducens); (R14) putative benzene hydroxylase and NADH:quinone oxidoreductase (*gmet 0231-0232*, G. metallireducens); (R15) phenylphosphate synthase (*ppsABC*, A. aromaticum EbN1, G. metallireducens); (R16) phenylphosphate carboxylase (*ppcABCD*, A. aromaticum EbN1, G. metallireducens); (R17) 4-hydroxybenzoate-CoA ligase (HbcL-1, A. aromaticum EbN1); (R18) 4-hydroxybenzoyl-CoA reductase (*hcrCAB*, A. aromaticum EbN1; PcmRST, G. metallireducens);. The information in this figure is compiled from the following studies on anaerobic benzene degradation [34, 22, 36, 41, 13, 24, 48, 62, 65, 70, 43, 53].

Fig. 13 illustrates the central degradation pathways. Dashed lines indicate putative reactions. Cofactors and conversion products thereof marked with a question mark indicate hypothesized compounds. Reactions for which enzymes and/or genes have been identified are labeled with numbered dots. Enzymes/genes involved in more than one pathway are highlighted in gray. Numbers for identifiers of compounds and genes/enzymes continue from Fig. 12. For clarity, identifiers for compounds and genes/enzymes also occurring in Fig. 12 have not been changed. (A) Upper pathways until hydrolytic ring cleavage. (B) Lower pathways comprising $\beta$-oxidation after hydrolytic ring cleavage for benzoate and subsequent degradation via the TCA cycle during respiration (upper panel) and fermentation based on S. aromaticivorans ([43]) (lower panel). Compounds: (19) benzoate; (24) benzoyl-CoA; (70) 2,3-epoxybenzoyl-CoA; (71) 3,4-dehydroadipyl-CoA semialdehyde; (72) 3,4-dehydroadipyl-CoA; (73) $\beta$-ketoadipyl-CoA; (74) acetyl-CoA; (75) succinyl-CoA; (76) cyclohex-1,5-diene-1-carbonyl-CoA; (77) 6-hydroxycyclohex-1-ene-1-carbonyl-CoA; (78) 6-ketocyclohex-1-ene-1-carbonyl-CoA; (79) 3-hydroxypimelyl-CoA; (95) 3-ketopimelyl-CoA; (96) glutaryl-CoA; (97) cortonyl-CoA; (98) 3-hydroxybutyryl-CoA; (99) acetoacetyl-CoA; (100) acetate; (101) butyryl-CoA; (102) butyrate; (HSCoA) coenzyme A; (Fd) ferredoxin; (Pi) inorganic phosphate. Enzymes/genes: (R13) benzoate-CoA ligase (*bclA*, A. aromaticum EbN1, Thauera spp., Magnetospirillum spp.; *bzdA*, Azoarcus spp.; *bamY*, G. metallireducens, S. aromaticivorans); (R13') succinyl-CoA:benzoate CoA transferase (*bct*, G. metallireducens); (R45) benzoyl-CoA oxygenase (*boxAB*, A. aromaticum EbN1, Azoarcus evansii); (R46) 2,3-epoxybenzoyl-CoA hydrolase (*boxC*, A. aromaticum EbN1, Azoarcus evansii); (R47) 3,4-dehydroadipyl-CoA semialdehyde dehydrogenase (*boxD*, Azoarcus evansii); (R48) $\beta$-ketoadipyl-CoA thiolase (*boxE*, Azoarcus evansii; *ebA2768*, A. aromaticum EbN1); (R49) benzoyl-CoA reductase (*bcrCBAD*, Thauera spp., Magnetospirillum spp., A. aromaticum EbN1; *bzdNOPQ*, Azoarcus spp.); (R49') benzoyl-CoA reductase (*bamBCDEFGHI*, G. metallireducens, S. aromaticivorans); (R50) cyclohex-1,5-diene-1-carbonyl-CoA hydratase (*dch*, Thauera spp., Magnetospirillum spp., A. aromaticum EbN1; *bzdW* Azoarcus spp.; *bamR* G. metallireducens, S. aromaticivorans); (R51) 6-hydroxycyclohex-1-ene-1-carbonyl-CoA dehydrogenase (*had*, Thauera spp., Magnetospirillum spp., A. aromaticum EbN1; *bzdX*, Azoarcus spp.; BamQ, G. metallireducens, S. aromaticivorans); (R52) 6-ketocyclohex-1-ene-1-carbonyl-CoA hydrolase (*oah*, Thauera spp., Magnetospirillum spp., A. aromaticum EbN1, *bzdY*, Azoarcus spp.; *bamA*, G. metallireducens, S. aromaticivorans); (R66) 3-hydroxypimelyl-CoA dehydrogenase (*fadB*, A. aromaticum EbN1; *gmet_2169*, G. metallireducens; *synarDRAFT_1233*, S. aromaticivorans); (R67) 3-ketopimelyl-CoA thiolase (*ebA2314*, A. aromaticum EbN1; *synarDRAFT_3465*, S. aromaticivorans) (R68) glutaryl-CoA dehydrogenase (*gcdH*, A. aromaticum EbN1; *bamM*, G. metallireducens; *synarDRAFT_2257*); (R69) enoyl-CoA hydratase (*fadB*, A. aromaticum EbN1; *gmet_2169*, G. metallireducens; *synarDRAFT_1515*, S. aromaticivorans); (R70) acetoacetyl-CoA reductase (*c2A173*, A. aromaticum EbN1; *gmet_1717*, G. metallireducens; *synarDRAFT_1514*, S. aromaticivorans) ; (R71) acetoacetyl-CoA thiolase (*fadA*, A. aromaticum EbN1; *gmet_1719*, G. metallireducens; *synarDRAFT_3465*, S. aromaticivorans); (R72) acyl-CoA synthase (*synarDRAFT_1015*); (R73) butyryl-CoA dehydrogenase (*synarDRAFT_1017*); (R74) acyl-CoA synthase (*synarDRAFT_1015*) [names for (R72-74) correspond to S. aromaticivorans]. The information in this figure is compiled from the following studies on anaerobic benzene degradation [9, 13, 22, 44, 43, 24, 36, 41, 53, 48]. Names for (R66-74) were derived from the KEGG PATHWAY Database (http://www.genome.jp/kegg/pathway.html) for A. aromaticum *ebN1*, G. metallireducens, and from the genome analysis of S. aromaticivorans by Nobu et al. [43].

##### 1.14 Selection of pathways for the characterization of anaerobic benzene metabolism

We used the scheme of anaerobic benzene degradation to identify the key enzymes and metabolites in the KEGG database. As such, we availed of so-called, custom pathways to screen our MAGs for their KEGG counterparts (Section 1.13). Importantly, not all supposed reactions are integrated in the current KEGG database. Further, we simplified the custom pathway of the TCA-cycle and included only the key enzymes. Each custom-made pathway visualized in heat-maps using the following scheme, absence: -1, presence: 0, after applying log2 transformation to transcription value for each KO. we used the R pachages: RColorBrewer, pheatmap and apcluster (Fig. 14 - Fig. 26). An overview was obtained by transformation of the data into into a graph representation, accounting only the mRNAs transcription and average the scores based on the custom-made pathway selection. In the graph, we further split each of the three initial step into two parts, the activation of benzene ring and the degradation to benzoyl-CoA. Briefly, Benzenestart 01, 02, 03, corresponds to anaerobic activation and conversion of benzene to benzoyl-CoA with intermediates benzylsuccinate, benzoate and hydroxybenzoate respectively. Then, Benzene mid aerobic is an aerobic hybrid pathway that convert benzoyl-CoA to acetyl-CoA and succinyl-CoA, which both of which fuel the TCA-cycle. In the other hand, Benzene mid anaerobic goes from benzoyl-CoA to 3-hydroxypimelyl-CoA anaerobically. The Benzene lower continues the anaerobic conversion to cortonyl-CoA. After that, we discriminate two types, Fermentation 1 to acetate and Fermentation 2 to butyrate and Respiration to acetyl-CoA that enters the TCA-cycle. Moreover, we also include a custom selection called Denitrification and dissimilatory nitrate reduction to ammonium (DNRA). In more detail, Benzene start 01 represents the upper path of Fig. 12 and correspond to reactions R05598, R05588, R05584, R05599, R05575 and R05587 from KEGG. On main Fig. 2 a&b, the first reaction (R05598) corresponds to the node Benzene open 01, while the rest reactions to Benzene CoA syn 01 (detailed heatmap: Fig. 14). Similarly, Benzene start 02 represents the middle path of Fig. 12 and correspond to R04986 (Benzene open 02) and R01422 (Benzene CoA syn 02)(detailed heatmap: Fig. 15). Benzene start 03 represents the lower path of Fig. 12 and correspond to reactions R03543 (Benzene open 03), R05625, R01300 and R05316 (Benzene CoA syn 03)(detailed heatmap: Fig. 16). Benzene mid aerobic represents the aerobic hybrid pathway of Fig. 13 A left side and correspond to reactions R09555, R09556, R09554 and R00829. (detailed heatmap: Fig. 18). Benzene mid anaerobic represents the anaerobic pathway of Fig. 13. Reactions at the right side correspond to reactions R09555, R09556, R09554 and R00829. (detailed heatmap: Fig. 18). Benzene lower represents the anaerobic conversion to cortonyl-CoA of Fig. 13 B at the left side and corresponds to reactions R05305 and R02488. (detailed heatmap: Fig. 19). Fermentation 1 represents the path to acetate of Fig. 13 B left side and corresponds to a discriminate reaction R0_72. (detailed heatmap: Fig. 20). Fermentation2 represents the path to butyrate of Fig. 13 B left side and corresponds to a discriminate reaction R0_73. (detailed heatmap: Fig. 21). Respiration represents the path to acetyl-CoA of Fig. 13 B left side and corresponds to reactions R02685, R03026, R01779 and R00238 (detailed heatmap: Fig. 22). TCA represents the TCA-cycle of Fig. 13 B at the left side and corresponds to the following custom reaction and KEGG reactions R02164_S, R02164_F, R0_glutamic_1, R0_glutamic_2, R0_citrate_1, R0_citrate_2, R00342 and R00360 (detailed heatmap: Fig. 23). Denitrification is a custom selection to represent the usage of nitrate and corresponds to R0_NarGHI, R0_NapAB, R0_NirK, R0_NirS, R0_NorCB and R0_NosZ reaction (detailed heatmap: Fig. 24). Key reactions of the glyxoylate shunt are R00479 and R00472 (detailed heatmap: Fig. 26).

### 2 Supplementary Results & Discussion

#### 2.1 Nitrogen cycling

We scanned the MAGs for genes encoding enzymes involved in dissimilatory reduction of nitrate or nitrite to ammonium (DNRA) or for denitrification (sequential reduction of nitrate to dinitrogen gas). DNRA enzyme systems are made up of a nitrate reductase (NarGHI or NapAB to reduce nitrate to nitrite) and a membrane bound cytochrome *c*-type nitrite reductase (NrfAH or NrfABCD-type to reduce nitrite to ammonium). Both nitrate reductases are membrane bound, but the NarGHI enzyme has its catalytic site inside the cell, whereas the one from the NapAB enzyme faces the outside of the membrane. This feature has implications for the bioenergetics such that NarGHI contributes to the generation of a *pmf* while NapAB is non-energetic. Denitrification pathways on the other hand usually also include one of these types of nitrate reductase, but the characteristic enzymes that discriminate denitrification from DNRA are a soluble nitrite reductase (NirK- or NirS-type), a nitric oxide reductase (NorCB or qNor), and a nitrous oxide reductase (NosZ). The *nirS* gene encodes a heme $cd_1$-type nitrite reductase, which requires iron for its hemes. The NirK-type enzyme is a trimer with copper as catalytic center. NorCB is a 2-subunit enzyme with heme-*c* containing NorC as the electron donor to the catalytic center of NorB. The qNor enzyme is a 1-subunit enzyme as a result of the fact that *norC* and *norB* paralogous sequences were fused somewhere during evolution. This enzyme receives electrons from quinol rather than from cytochrome *c* in the case of NorCB. Most of the 47 MAGs with their MAGs that we ultimately selected have the potential to perform DNRA (eight MAGs), denitrification (31 MAGs), or combinations of that (five MAGs). Only three MAGs express a nitrate reductase without an accompanying nitrite reductase. It may be that they do not have that genetic potential at all, or that a contig with that genetic information did not meet the criteria for proper binning. DNRA with nitrate as initial electron acceptor is likely to occur in five MAGs (these have a NarGHI or NapAB-type nitrate reductase and NrfABCD), with nitrite in eight MAGs (only NrfA). Notably, genes encoding the alternative nitrite reductase NrfABCD were not found in either of the MAGs. The five MAGs that have the genetic potential to perform both DNRA and denitrification are not restricted to certain phyla although three of them belong to the Chloroflexi. Two of the five MAGs lack a nitrate reductase and hence they use nitrite as initial oxidant. The reason for having both types of nitrate or nitrite reducing capacities is not clear. Typical differences between the two is that DNRA as compared to denitrification produces less of the cytotoxic intermediate NO, is faster in terms of rates of electron transfer between donor and acceptor and is more potent in electron accepting as nitrate reduction to ammonium requires 8 electrons rather than the 5 needed to reduce nitrate to dinitrogen during denitrification. This may be important for redox balancing during free energy transduction. The trade-off for using DNRA on the other hand is that the generation of the proton motive force during the full reduction of 1 nitrate molecule to ammonium is 25% less efficient as compared to denitrification (18 charge separations during transfer of 8 electrons during DNRA versus 15 per 5 electrons during denitrification). Organisms that have both trades can opt for the best solution at every change in their growth condition. A characteristic feature of denitrification is that an N-oxide ion is reduced to an N-oxide gas, a step catalyzed by a NirS or NirK-type enzyme. Only MAGs that have either type of reductase are true denitrifiers. These may be split up in three groups, those that have the *nirS* gene (one MAG), those with *nirK* (23 MAGs) and those that have both types (12 MAGs). The latter organisms are therefore versatile with regard to metal availability. Most of the *nirS* genes are found in the proteobacteria (nine MAGs), *nirK* is more prevailing in all other phyla. Further inspection of the genetic potential to carry out denitrification identified six MAGs, five of which proteobacteria, that have a full complement of genes to make all four enzymes of denitrification. The nitrate reductase most frequently found in the MAGs is of the NarGHI-type (25 MAGs). The NapAB-type is rare and found in only three MAGs, all of which are proteobacteria. Two of these have even both types of nitrate reductase. The denitrifying MAGs further display a variety of gene combinations, the most abundant one is the combination of genes encoding a nitrate and nitrite reductase (11 MAGs), or just a single nitrite reductase (23 MAGs). The fact that more than half of the denitrifiers seem to be able to reduce nitrite to nitric oxide, but do not have the means to convert this cytotoxic molecule any further as they lack the potential to make either type of nitric oxide reductase, may seem quite peculiar for an organism. From a community perspective, however, this might make sense since NO is a freely diffusible gas that may be converted into harmless nitrous oxide by other community members that do have *nor* genes (18 MAGs). Notably as well is the observation that there are five MAGs with a gene encoding qNor (one MAG) or with a *norCB* gene cluster (four MAGs) without a gene encoding an accompanying nitrite reductase. It might well be that these MAGs have found a niche to efficiently detoxify the community on the one hand, while using NO as electron acceptor on the other hand. We were also aware of the fact that a paralogue of the qNor protein was described in a previous study [5]. This paralogue is called nitric oxide dismutase (NOD) and it shares a high degree of homology with canonical qNor proteins. But where qNor reduces nitric oxide to nitrous oxide, NOD dismutates 2 molecules of NO into an oxygen molecule and a dinitrogen molecule without the need for electrons. This reaction has implications for the enzyme make up. The electron pathway in qNor involves a low spin heme *b* for transfer of electrons to the high spin heme in the reaction center. Both hemes are ligated via positionally conserved histidine residues next to one another in helix 10 of the enzyme. NOD does not require electrons so it lost the heme and the allocated histidine during evolution. The absence of this histidine is characteristic for NOD enzymes and discriminates them from the Nor enzymes. NOD enzymes that share this property were found in MAGs 34 and 71, although the latter did not meet the criteria for selection into the MAG list. Nevertheless, these MAGs may well be able to carry out NO dismutation to produce some oxygen for themselves or for community members. Indeed, it was shown that the benzene degrading culture produced oxygen once that it was pulsed with nitrite [5, 69, 64]. Some MAGs may use locally produced traces of oxygen for their mono- or dioxygenases for breakdown of benzene under an otherwise anaerobic atmosphere. Indeed, we noticed the upregulation of mRNAs encoding such oxygenases in some of the MAGs. The list of 47 MAGs also includes a member of the Planctomycetes, order Brocadiales, which is very similar to *Candidatus* Kuenenia Stuttgartiensis. The latter MAG carries out the process termed anammox, anaerobic ammonium oxidation. All enzymes for this free energy transducing process are located within a so-called anammoxosome, a unique type of organelle. The key enzyme in this process is a hydrazine synthase that combines NO and ammonium to form hydrazine that is subsequently oxidized to dinitrogen by a hydrazine dehydrogenase. Hydrazine synthase is made up of three subunits. The sequences of all 3 of them were deduced from this MAG and they had high resemblance with those from *Candidatus* Kuenenia Stuttgartiensis. It is, therefore, fair to say that a MAG from the Brocadiacea is a member of the community in the benzene degrading culture. Also, in this case, a MAG has found its own niche in the continuous culture as it has its unique type of autotrophic metabolism with ammonium as the free energy source and carbon dioxide as the carbon source. This suggestion is further corroborated by the fact that a MAG from the Brocadiaceae has been isolated in pure culture some years ago ([69]).

##### 2.2 Species and taxonomy

MAGs were assigned to the following phyla: 33 Proteobacteria, 17 Bacteroidetes, 15 Chloroflexi, 11 Actinobacteria, seven Planctomycete, five Verrucomicrobia, four Acidobacteriota, four Myxococcus, three Gemmatimonadetes, two Hydrogenedentota, one of each: Armatimonadota, Firmicutes, Spirochaetota, Zixibacteria, Deinococcota and three unknown. Our binning methodology and verification ultimately resulted in 47 assembled genomes with a high-quality standard. The most dominant phylum is that of the Proteobacteria (16 MAGs). Five of these are $\alpha$-protobacteria with the order Rhizobiales as the most prevailing one (four, one of which close to *Ochrobactrum anthropi*). The 5th one is *Paracoccus saliphilus* from the order Rhodobacterales. *Rhizobiales* and *Paracoccus* MAG are known as typical nitrogen fixing organisms with a versatile metabolism, usually able to denitrify under oxygen depletion. The $\beta$-proteobacteria contribute with seven MAGs, six of which are from the order Burkholderiales (some closely related to *Candidimonas bauzanensis*, *Comamonas testosteroni*, and *Hydrogenophaga intermedia*), and the 7th from the Rhodocyclales (*Propionivibrio sp.*). Both the Burkholderiales and Rhodocyclales are described as dominant phylotypes in enrichment cultures of anaerobic benzene-degrading microcosms and are suggested to use the methylation pathway for anaerobic benzene activation [63, 1, 69]. There are further three $\gamma$-proteobacteria, order Xanthomonadales and one $\delta$-proteobacterium. Xanthomonadales have the genetic potential to make type IV secretion systems, which are able to secrete toxins directly into their prokaryotic target species. Hence, they apparently feed on rival bacterial cells [58]. The community also contains 10 MAGs from the phylum Bacteroidetes, with classes Ignavibacteria (seven), Bacteroidia (two), and Chitinophagia (one, *Niastella yeongjuensis*). Members from the Bacteroidetes were regarded as putative scavengers during syntrophic breakdown of benzene [61]. It has also been observed that many types of Bacteroidales have gene clusters encoding so-called type VI secretion systems, which is apparent in gut microbiomes [15]. It has been suggested that enteric pathogens use type VI secretion systems to antagonize symbiotic gut *E. coli*, facilitating colonization and disease progression.

The phylum Chloroflexi is represented by eight MAGs, three of which belong to families *Anaerolineaceae* (three, one is close to *Bellilinea caldifistulae*), *Ardenticatenaceae* (four, one close to *Ardenticatena sp.*) and Thermomicrobia (one). It is an ecologically and physiologically diverse group of bacteria, which have been detected in an increasingly wide range of anaerobic habitats including sediments, hot springs, methanogenic anaerobic sludge digesters where they are highly abundant. Chloroflexi seem to play an important role in filamentous scaffolding around which flocs are formed to ferment intracellular compounds from lysed bacterial cells to low molecular weight substrates to support their growth and that of others in a community [59, 68]. *Anaerolineaceae* bacteria occur in marine sediments. Culture-independent analyses revealed *Anaerolineaceae* as abundant primary fermenters in anaerobic digesters treating waste activated [40]. We also noticed the presence of three MAGs belonging to the phylum Gemmatimonadetes, two of which were in the genus *Gemmatimonas* (*Gemmatimonas phototrophica*). The type species is *Gemmatimonas aurantiaca*. This bacterium has been identified as a member of one of the top nine phyla found in soils; yet, there are currently only six cultured isolates. Gemmatimonadetes have been found in a variety of arid soils, such as grassland, prairie, and pasture soil, as well as eutrophic lake sediments and alpine soils. The metabolic pathways and enzymes of this bacterium are unique and it is able to grow by both aerobic and anaerobic respiration. Two MAGs with relatively high abundance are from the phylum Planctomycetes, 1 from class Brocadiae (close to *Candidatus* Kuenenia Stuttgartiensis), and the other from class Phycisphaerae (*Phycisphaera sp.*). *Candidatus* Kuenenia Stuttgartiensis has a special type of free energy metabolism that takes place in a so-called anammoxosome. There, ammonium is used as free-energy source during oxidation to dinitrogen gas with nitrite as electron acceptor and hydrazine as intermediate compound. Carbon dioxide serves as the carbon donor for cell biomass formation. Phycisphaerae belong to the same phylum, but are not able to perform anammox. They are sometimes found together in anaerobic ammonium oxidizing cultures [66]. The MAG with contigs that belong to this Phycisphaerae was, however, not selected for further analyses as it did not meet the criteria. We found only a single MAG of the Firmicutes, and this one belongs to the family *Peptococcaceae*, closely related to *Desulfonispora thiosulfatigenes*. *Peptococcaceae* are also well known for their role in anaerobic benzene degradation and it has been suggested that they use the carboxylation pathway [63, 1, 69, 64, 5].

#### 2.3 Expression levels mRNAs

We inspected the mRNAs that were expressed at relatively high levels in each of the MAGs. It shows that all Chloroflexi (MAGs 1, 5, 26, 31, 39, 41, 42) express relatively high levels of mRNAs encoding peptidases, solute binding proteins and ABC-type transporters. This property is shared with 1 of the Bacteroidetes (MAG 18). High levels of messengers encoding the key enzyme of the aerobic middle pathway for benzene degradation are found exclusively in most of the $\beta$-proteobacteria (MAGs 19, 36, 47, 56 and 68). Some of these also express mRNA for making another key enzyme of benzene degradation, 3-octaprenyl-4-hydroxybenzoate carboxy-lyase (MAGs 19, 25 and 68). We also noted some upregulation of mRNAs encoding superoxide dismutase (SODs) in species that have their genomes in MAGs 19, 29, and 47 all of which proteobacteria. Secretion systems of types II, III and VI may be operational in some species as well since we found relatively high concentrations of mRNAs related to type II in members of Armatimonadetes (MAG 22) and Actinobacteria (MAG 66), to type III in MAG 22 as well, and to type VI in members of Acidobacteria (MAG 7) and Ignavibacteria (MAG 10). Some species (a Chloroflexi, MAG 31 and Ignavibacteria, MAG 49) have high levels of mRNA encoding reverse transcriptase. These enzymes are typical for certain types of bacteriophages, entities that rely on host cells to propagate. The fate of the host cell is lysis. The species in MAG 29, a $\gamma$-proteobacterium, has a relatively high concentration of mRNA to make a chemotaxis protein. Perhaps it uses that to find its way to certain nutrients. Messengers indicative for making a flagellum for movement were found in a member of the *Peptococcaceae* (MAG 9), and in a $\delta$-proteobacterium (MAG 14). The latter one, though, has also mRNAs indicative or a biofilm lifestyle so perhaps it could switch from one state to another. Biofilm mRNAs were also apparent in the member of the Armatimonades (MAG 22), and the $\gamma$-proteobacterium in MAG 34. These findings are in perfect agreement with the prediction of their preferred location on the basis of comparison of mRNA levels of each species between biofilm and liquid (see Table). We here hypothesize that the concentration of the mRNAs is a reflection of their abundance in either location. Overall it shows that only 5 species seem to prevail exclusively in the liquid, 3 of which are Bacteroidetes, the 4th one a Verrucomicrobium and the 5th one in a member of the *Peptococcaceae*. All others feel more comfortable in the biofilm or switch from one location to the other. Relative high levels of expression of mRNAs encoding nitrous oxide reductases and haem copper oxidases were also apparent. The first type of terminal oxidoreductase may well be important in Ignavibacteria (MAGs 20 and 101), and Chloroflexi (MAGs 26 and 31), the second one in $\beta$-proteobacteria (MAG 25, 36, 68 and a Verrumicrobia (MAG 30). Specific and more exclusive mRNAs are also apparent in a number of species. Here is a brief description of the potential enzymes and the species that make it. Subtilase by an Acidobacteria (MAG 7), a key enzyme of glycerol uptake and metabolism by a Chloroflexi (MAG 42), fatty acid uptake and metabolism by an Ignavibacteria (MAG 49) and a Verrumicrobia (MAG 12), enzymes for methylamine metabolism in an a-proteobacterium (MAG 16, their genes are clustered), enzymes for formate metabolism in yet another a-protobacterium (MAG 4, genes are clustered). Last but not least, a protocatechuate 4,5-dioxygenase in 2 different $\beta$-proteobacteria (MAGs 19 and 68, successively). The enzyme catalyzes the oxygen dependent ring opening of 3,4-dihydroxybenzoate to yield 4-carboxy-2-hydroxymuconate semialdehyde.

The analyses of the list of relatively high concentrations of mRNAs revealed some interesting features that could shed some light on the observation that so many different species live together in a bioreactor with benzene as main carbon and free energy source and nitrate as the electron acceptor. We have good reasons to believe that there are only a few primary consumers of the benzene, the most dominant of which is a member of the *Peptococcaceae*. Another niche within the community is occupied by a member of the Planctomycetes that is closely related to *Candidatus* Kuenenia stuttgartiensis. The latter is specialized in anaerobic ammonia oxidation (Anammox) for its free energy transduction. Carbon dioxide rather than benzene serves as the carbon source in this species. Hence, there is no competition for food between the benzene degraders and the ammonium oxidizer. Such a autotrophic niche could make a significant contribution to the overall carbon flux and the carbon produced by it may enter into other heterotrophic cycles in the system. Additionally, autotrophic processes can deplete electron acceptors which then are not available anymore for benzene degradation [27, 32, 23]. Overall, the Planctomycetes may play an important role on ramifications for benzene degradation. Other trades of growth are obvious in the Chloroflexi. All of them have relatively high levels of mRNAs encoding both peptidases, solute binding proteins and ABC transporters suggestive for their ability to cut larger peptides or sugar polymers into smaller molecules that can be transported to the inside of the cell via these dedicated transport systems. Once inside the cell, these molecules may serve as carbon- and free energy source. Yet other species appear to make enzymes for fatty acid uptake and metabolism. The question then becomes apparent what do they scavenge on. We speculate the source of these peptides, sugar polymers and fatty acids are the remnants of dead cells that lysed either passively or actively. Such biomass scavengers may play an important role in anaerobic systems and may influence the carbon turnover and interfere with anaerobic/syntrophic benzene degradation. They may produce hydrogen through the fermentation of biomass-derived compounds which might in turn inhibit benzene fermentation. Moreover, important co-factors like vitamins that other members of the community including primary benzene-degraders might not be able to produce themselves could be produced by the scavenging members of the community [16, 28, 67, 45]. Likely candidates for active lysis are the suggested bacteriophages. In this view it is also notable that some species are able to recruit secretion systems of either type II, III or VI. These systems allow them to penetrate a host cell and inject toxins or other types of effector molecule in them, ultimately resulting in cell death [37]. We inspected the locus of the gene encoding the tube protein of the secretion system of type VI in MAG 7. It appears to be clustered along with genes encoding the tip protein, baseplate, ATPase and contractile sheath. In addition, a gene encoding lysozyme was found sandwiched between them, and this might be the specific effector molecule of the TSS system in order to make holes in the membrane of the target cell. A similar type of clustering of genes encoding the subunits of the TSS VI secretion system was found in MAG 10. MAGs 7 and 10 may be considered apex predators that occupy the highest trophic levels within the food web. Alternatively, cells may lyse by the species-specific bacteriophages. It has been estimated that about half of a community's species may be lysed by phages [56]. The observation of high levels of mRNAs in some of the MAGs encoding reverse transcriptase may add thought to this. The suggested niches are a likely explanation for the variety of species in the culture. Yet, survival is not only dependent on the rate and efficiency of free energy metabolism, but certainly also of the growth rate. Would it be slower than the dilution rate, than the species would flush out of the culture. So, the only explanation for the fact that there are still slow growers in the culture is the presence of the biofilm, which would retain those organisms that have such a low growth rate. One clear example concerns the Anammox species that doubles every 20 days [66]. Indeed, we do see mRNAs popping up in some species that indicate their prevalence in the biofilm. Yet others favor a free-living life style with a flagellum that would allow them to swim to more nutrient rich areas. Some species appear to be specialists in certain types of metabolism and complement other types that we rationalized before. These are the ones for example that have the enzymes either to metabolize methylamine, or formate or glycerol. These molecules are likely to be formed during anaerobic digestion of the peptides and carbohydrates. This suggestion is further corroborated by the finding that the genes encoding the mRNAs for the respective enzymes are clustered on their respective genomes, suggestive for coordinated expression of them.

##### 2.3.1 The oxygen paradox

The culture is anoxic with a completely anaerobic atmosphere. Yet we find the high-level expression of enzymes that require oxygen for their activity like terminal oxidases, monooxygenases and dioxygenases in some of the species. The species with its genome in MAGs 25, 36, 56, 68, 69 and 92 have expressed their genes encoding the high affinity $cbb_3$-type oxidase. Cytochrome *c* oxidases of the Verrumicrobial-type are in MAG 30, and $\beta$-proteotype in 48. This seems peculiar in terms of efficiency of making proteins that are supposedly not active under anaerobic conditions. The only likely hypothesis is that there is a local production of oxygen where some species can benefit from. As already mentioned, the most likely oxygen producing enzyme in the (dark) culture is the NO dismutase, the genes of which were identified both in MAG 34 and 71. Unfortunately, the latter did not meet the criteria for further analyses, but it is evident that there is a second copy of the NO dismutase gene in the culture. NO and $N_2O$ are freely diffusible gases and may well be exchanged between the species. The NOD may even be fueled by externally produced NO. This is what is to be expected since many species appear to have a NirS- or NirK-type nitrite reductase to produce NO but not an accompanying NO reductase to consume it again. The assumption that there is a continuous flux of oxygen produced by the NOD members then also explains the expression of oxygenases in MAGs 19 and 68, both of which from *Burkholderia* species of the $\beta$-proteobacteria. They apparently rely on aerobic mid-pathways for degradation of benzene. Both MAGs have high expression of mRNAs encoding 3-octaprenyl-4-hydroxybenzoate carboxy-lyase, of benzoyl CoA oxygenase and of protocatechuate 4,5-dioxygenase. Recruitment of their enzymes allows them to convert benzene to benzoate and next either to epoxy benzoyl-CoA or to 4-carboxy-2-hydroxymuconate-6-semialdehyde. The exact nature of the suggested syntrophic interaction between the species with MAG 34 and the *Burkholderia* species remains to be elucidated. If oxygen would be a freely diffusible gas in the culture, one would also expect defense mechanisms against it. Indeed, we see upregulation of genes encoding Fe/Mn-type SODs in some of the proteobacteria. Notably, these are also the species that have the monooxygenases. Could it be that they are in close proximity of the oxygen producers such that they can optimally profit from the oxygen flux. If so, also the steady state concentration of oxygen is expected to be very low in the remaining part of the pot.

#### 2.4 Classification scheme on benzene metabolism

We summarized the previous obtained results into a simplified view of microbial community structure. We classified each MAG as high/medium/low/none, for their potential to be a primary consumer based on their capacity to activate benzene and further degrade it. We classified further their functional diversity based on the assigned functional groups from before (See main results) and if they are dominant or non-dominant members of the community (See main results). Also, we looked in which phase on the bioreactor they found with higher actively (biofilm, liquid or both) based on the transcriptomic profiles - based on the previous results (See main results). Therefore, in relation to classification for primary consumer of benzene, if a MAG does not contain any KOs present or transcribed in the following selections: Benzene open 01, Benzene open 02, Benzene open 03, Benzoyl CoA syn 01, Benzoyl CoA syn 02, Benzoyl CoA syn 03, then it is assigned into the class "none". if it contains present or transcribed KOs, but the selections were not sequential coupled (open 01–CoA syn 01, open 02–CoA syn

02 and open 03–CoA syn 03), then it assigned as class "low" and if the selection were sequential coupled then assigned as class "medium". For a MAG to assigned as class "high" it must contain transcribed KOs on at lease one of the three sequential couples and transcribed KOs at least in one of the sequential next step (Benzene mid aerobic or Benzene mid anaerobic).

##### 2.5 Benzene degradation pathways

MAG 16 is the only one with genes encoding enzymes for benzene degradation to benzoyl-CoA through benzyl succinate. These enzymes are benzylsuccinate CoA-transferase, (R)-benzylsuccinyl-CoA dehydrogenase, E-phenylitaconyl-CoA hydratase and benzoylsuccinyl-CoA thiolase Fig. 14. Another few MAGs appear to degrade benzene through benzoate using UbiD/UbiX-related carboxylases, benzoate-CoA ligase and benzoyl-CoA reductase. Relatively high transcription levels of their mRNAs are found in MAGs 6 and 9, and lower levels in MAGs 19, 25 and 68 Fig. 15. We did not find MAGs that have a full suite of genes for benzene degradation via the hydroxybenzene route. Apparently, benzene is converted to benzoyl-CoA through benzoate and/or hydroxybenzoate routes by most of the MAGs that belong to the primary consumers. The route through benzyl succinate may only be used by MAG 16. Further metabolism of benzoyl-CoA may proceed through a strict anaerobic route or via one that includes an oxygen dependent benzoyl-CoA oxygenase. The anaerobic route requires genes that encode benzoyl-CoA reductase, cyclohexa-1,5-dienecarbonyl-CoA hydratase, 6-hydroxycyclohex-1-ene-1-carbonyl-CoA dehydrogenase, 6-oxocyclohex-1-enecarbonyl- CoA hydrolase. Their transcripts were found at relatively high levels in MAGs 4, 6, 9 and 18 and to a lesser extent in MAGs 14, 19, 25, 68 and 101. Interestingly, genes (BamB/BamC) related to benzoyl-CoA reductase were found exclusively in MAG 9 on an even higher transcription level than the rest Fig. 17. The route that includes benzoyl-CoA oxygenase may be operative in a few Proteobacterial MAGs (MAGs 4, 19, 25, 36, 47, 56 and 68) as they contain genes for benzoyl-CoA 2,3-epoxidase, benzoyl-CoA-dihydrodiol lyase, 3,4-dehydroadipyl-CoA semialdehyde dehydrogenase, acetyl-CoA acyltransferase and 3-oxo-5,6-didehydrosuberyl-CoA/3-oxoadipyl-CoA thiolase. Also, although no complete sequential genes to complete the pathway found in any MAG, it is surprising that in this anoxic environment we find genes indicating toluene monooxygenase and phenol hydroxylase. Those proteins oxidise benzene to catechol and catechol 2,3-dioxygenase activates oxidative ring cleavage of catechol, which is then further converted to pyruvate and acetyl-CoA by enzymes of the lower pathway - indicates a possible syntrophic breakdown of benzene by aerobic degradation [5] (See Fig. 18). Overall, this result indicates that only a few MAGs activate and further degrade benzene as the primary consumers, while the majority of the community does not and occupies other niches, ultimately resulting in a complex and extended food web.

**Table 1.** Basic genome information, quality and relative RNA & DNA abundance of all MAGs. The grey rows are the filtered out MAGs based on the quality and manual refinement

| | Genome information | | | | MAG quality | | Abundance | | | |
|---|---|---|---|---|---|---|---|---|---|---|
| MAGs | total length | num contigs | N50 | GC content | Completion % | Redundancy % | DNA | RNA - Biofilm | RNA - Liquid | Orf count |
| 000 | 60499047 | 15403 | 8098 | 60.47 | 100.00 | 842.25 | 0.35 | 2.16 | 3.71 | 64790 |
| 001 | 3435559 | 60 | 121528 | 59.27 | 94.37 | 1.41 | 24.34 | 5.57 | 0.66 | 3329 |
| 002 | 1944298 | 36 | 56800 | 36.61 | 84.51 | 2.82 | 15.27 | 3.71 | 2.57 | 1727 |
| 003 | 3959694 | 208 | 30344 | 40.77 | 98.59 | 12.68 | 7.01 | 27.07 | 4.20 | 3715 |
| 004 | 3822053 | 27 | 206048 | 65.67 | 100.00 | 2.82 | 4.22 | 0.35 | 0.02 | 3739 |
| 005 | 3739161 | 57 | 149802 | 54.03 | 95.77 | 2.82 | 4.60 | 3.01 | 0.23 | 3610 |
| 006 | 3311055 | 120 | 60502 | 64.13 | 97.18 | 7.04 | 3.57 | 1.94 | 0.58 | 3377 |
| 007 | 3335773 | 15 | 392789 | 53.74 | 95.77 | 1.41 | 3.16 | 0.20 | 0.02 | 3089 |
| 008 | 1249458 | 26 | 26585 | 62.65 | 47.89 | 0.00 | 2.99 | 3.29 | 0.27 | 1153 |
| 009 | 4248075 | 93 | 87433 | 44.35 | 98.59 | 0.00 | 2.97 | 36.07 | 75.89 | 4063 |
| 010 | 3685673 | 287 | 23711 | 33.34 | 97.18 | 1.41 | 2.17 | 0.47 | 0.04 | 3522 |
| 011 | 253085 | 19 | 65316 | 59.63 | 0.00 | 0.00 | 2.18 | 2.76 | 1.96 | 256 |
| 012 | 4597136 | 78 | 79543 | 62.51 | 94.37 | 2.82 | 1.43 | 0.55 | 0.00 | 3791 |
| 013 | 3008466 | 28 | 331130 | 58.98 | 97.18 | 0.00 | 1.49 | 1.03 | 0.06 | 2470 |
| 014 | 8076467 | 148 | 81157 | 69.94 | 95.77 | 0.00 | 1.26 | 0.10 | 0.01 | 7282 |
| 015 | 242034 | 32 | 51870 | 57.42 | 0.00 | 0.00 | 1.60 | 0.61 | 0.22 | 246 |
| 016 | 4189325 | 81 | 178727 | 62.81 | 97.18 | 5.63 | 1.11 | 0.20 | 0.03 | 3974 |
| 017 | 2684626 | 610 | 6114 | 69.99 | 87.32 | 4.23 | 0.98 | 0.16 | 0.05 | 2945 |
| 018 | 3086012 | 19 | 283578 | 32.96 | 94.37 | 1.41 | 1.03 | 0.98 | 1.99 | 2707 |
| 019 | 4074397 | 166 | 48779 | 69.79 | 91.55 | 7.04 | 0.93 | 0.10 | 0.06 | 4037 |
| 020 | 3687606 | 59 | 107749 | 40.58 | 97.18 | 1.41 | 0.89 | 1.17 | 1.37 | 3009 |
| 021 | 3038481 | 56 | 54077 | 59.94 | 88.73 | 0.00 | 0.83 | 0.08 | 0.03 | 758 |
| 021.1 | 822707 | 39 | 23312 | 59.08 | 0.00 | 0.00 | 0.84 | 1.15 | 0.50 | 2562 |
| 022 | 3247912 | 35 | 94516 | 63.41 | 94.37 | 1.41 | 0.76 | 0.13 | 0.01 | 2934 |
| 023 | 4665751 | 234 | 41764 | 61.10 | 100.00 | 7.04 | 0.71 | 0.20 | 0.06 | 4582 |
| 024 | 3552934 | 445 | 13239 | 67.68 | 85.92 | 7.04 | 0.65 | 0.08 | 0.03 | 3681 |
| 025 | 3756427 | 64 | 109280 | 70.77 | 92.96 | 0.00 | 0.65 | 0.06 | 0.02 | 3504 |
| 026 | 3639465 | 40 | 253413 | 61.13 | 95.77 | 0.00 | 0.54 | 0.12 | 0.00 | 3166 |
| 027 | 2308863 | 797 | 3858 | 66.45 | 83.10 | 14.08 | 0.52 | 0.73 | 0.66 | 2936 |
| 028 | 7069549 | 1158 | 9164 | 71.03 | 74.65 | 8.45 | 0.43 | 0.08 | 0.01 | 7201 |
| 029 | 2957553 | 179 | 30579 | 68.65 | 97.18 | 8.45 | 0.40 | 0.11 | 0.06 | 2670 |
| 030 | 4181919 | 23 | 212925 | 66.55 | 98.59 | 1.41 | 0.33 | 0.15 | 0.32 | 3339 |
| 031 | 4931782 | 51 | 149493 | 59.49 | 97.18 | 1.41 | 0.31 | 0.11 | 0.01 | 4337 |
| 032 | 3328800 | 100 | 101615 | 36.55 | 100.00 | 0.00 | 0.31 | 0.04 | 0.01 | 2817 |
| 033 | 2779610 | 39 | 77228 | 53.96 | 83.10 | 1.41 | 0.28 | 0.12 | 0.00 | 2398 |
| 034 | 2554880 | 270 | 16787 | 62.66 | 91.55 | 9.86 | 0.25 | 0.05 | 0.01 | 2572 |
| 035 | 3845035 | 144 | 57260 | 56.22 | 92.96 | 0.00 | 0.23 | 0.13 | 0.00 | 3539 |

| | | | | | | | | | | |
|---|---|---|---|---|---|---|---|---|---|---|
| 036 | 4639052 | 75 | 136744 | 70.10 | 94.37 | 4.23 | 0.21 | 0.10 | 0.03 | 4369 |
| 037 | 3655018 | 572 | 11089 | 66.75 | 97.18 | 2.82 | 0.25 | 0.07 | 0.01 | 3990 |
| 038 | 5538605 | 73 | 111952 | 67.75 | 100.00 | 2.82 | 0.21 | 0.03 | 0.00 | 4551 |
| 039 | 2737729 | 173 | 23317 | 71.08 | 92.96 | 2.82 | 0.20 | 0.02 | 0.00 | 2722 |
| 041 | 4421002 | 227 | 28430 | 63.52 | 94.37 | 1.41 | 0.20 | 0.07 | 0.00 | 3911 |
| 042 | 3428056 | 72 | 60132 | 68.40 | 90.14 | 2.82 | 0.19 | 0.03 | 0.00 | 3182 |
| 043 | 3659643 | 298 | 22797 | 72.94 | 63.38 | 1.41 | 0.25 | 0.12 | 0.01 | 2981 |
| 044 | 5193204 | 420 | 20074 | 76.37 | 38.03 | 1.41 | 0.21 | 0.05 | 0.00 | 4122 |
| 045 | 5148383 | 324 | 49748 | 62.30 | 98.59 | 36.62 | 0.16 | 0.07 | 0.05 | 5182 |
| 046 | 2515780 | 20 | 101247 | 46.99 | 87.32 | 0.00 | 0.31 | 0.10 | 0.03 | 2159 |
| 047 | 4353019 | 111 | 83634 | 61.42 | 98.59 | 5.63 | 0.14 | 0.07 | 0.02 | 4008 |
| 048 | 4940754 | 322 | 36850 | 61.17 | 98.59 | 0.00 | 0.14 | 0.08 | 0.01 | 4443 |
| 049 | 3998798 | 253 | 42236 | 35.87 | 97.18 | 1.41 | 0.20 | 0.09 | 0.03 | 3621 |
| 050 | 3005714 | 199 | 26951 | 61.50 | 95.77 | 2.82 | 0.20 | 0.10 | 0.01 | 2596 |
| 051 | 4088151 | 870 | 8436 | 70.83 | 98.59 | 38.03 | 0.12 | 0.03 | 0.00 | 4835 |
| 052 | 3442950 | 118 | 62564 | 65.04 | 87.32 | 11.27 | 0.14 | 0.35 | 1.27 | 3333 |
| 053 | 5493707 | 338 | 54079 | 68.09 | 95.77 | 91.55 | 0.13 | 0.07 | 0.04 | 5398 |
| 054 | 2864095 | 278 | 18657 | 69.50 | 49.30 | 11.27 | 0.13 | 0.10 | 0.07 | 2879 |
| 055 | 3455486 | 403 | 14004 | 70.78 | 35.21 | 11.27 | 0.12 | 0.04 | 0.01 | 3317 |
| 056 | 5495738 | 515 | 30096 | 63.45 | 100.00 | 7.04 | 0.08 | 0.04 | 0.02 | 5384 |
| 057 | 2617295 | 341 | 13652 | 66.00 | 78.87 | 4.23 | 0.26 | 0.58 | 0.48 | 2830 |
| 058 | 5192471 | 184 | 83902 | 46.87 | 97.18 | 1.41 | 0.21 | 0.20 | 0.14 | 4207 |
| 059 | 3146082 | 29 | 339977 | 50.56 | 95.77 | 0.00 | 0.21 | 0.10 | 0.00 | 2698 |
| 060 | 9936556 | 1032 | 18963 | 60.53 | 92.96 | 22.54 | 0.11 | 0.07 | 0.00 | 8748 |
| 061 | 1654255 | 372 | 7287 | 62.23 | 57.75 | 23.94 | 0.17 | 0.45 | 0.57 | 1969 |
| 062 | 5005696 | 178 | 48554 | 55.90 | 98.59 | 1.41 | 0.12 | 0.02 | 0.01 | 4898 |
| 063 | 6354922 | 1647 | 6194 | 74.93 | 100.00 | 38.03 | 0.09 | 0.01 | 0.00 | 7567 |
| 064 | 3202270 | 281 | 22637 | 67.43 | 100.00 | 1.41 | 0.11 | 0.01 | 0.00 | 3140 |
| 065 | 6744885 | 2039 | 8458 | 50.26 | 85.92 | 43.66 | 0.13 | 0.07 | 0.03 | 8033 |
| 066 | 2817429 | 255 | 24209 | 65.56 | 91.55 | 1.41 | 0.13 | 0.02 | 0.01 | 2970 |
| 067 | 3530919 | 315 | 16384 | 40.32 | 97.18 | 5.63 | 0.13 | 0.08 | 0.10 | 3132 |
| 068 | 4322486 | 485 | 13567 | 68.46 | 95.77 | 4.23 | 0.11 | 0.05 | 0.04 | 4388 |
| 069 | 2926801 | 173 | 30327 | 35.28 | 94.37 | 1.41 | 0.10 | 0.13 | 0.41 | 2663 |
| 070 | 3525412 | 94 | 55493 | 55.04 | 94.37 | 1.41 | 0.14 | 0.09 | 0.01 | 3056 |
| 071 | 5925308 | 635 | 40486 | 70.34 | 92.96 | 29.58 | 0.11 | 0.01 | 0.00 | 5783 |
| 072 | 8758907 | 625 | 39721 | 57.95 | 94.37 | 28.17 | 0.10 | 0.02 | 0.00 | 7748 |
| 073 | 2705742 | 1295 | 2274 | 64.66 | 35.21 | 9.86 | 0.10 | 0.01 | 0.00 | 3576 |
| 074 | 3806583 | 556 | 13834 | 67.89 | 66.20 | 11.27 | 0.08 | 0.02 | 0.01 | 4059 |
| 075 | 7109454 | 1484 | 8254 | 63.41 | 97.18 | 45.07 | 0.07 | 0.11 | 0.00 | 7136 |
| 076 | 2140732 | 272 | 11740 | 69.90 | 90.14 | 2.82 | 0.09 | 0.03 | 0.00 | 2104 |
| 077 | 4673905 | 672 | 15886 | 67.63 | 70.42 | 25.35 | 0.08 | 0.07 | 0.02 | 4505 |
| 078 | 2641288 | 777 | 4822 | 67.84 | 33.80 | 4.23 | 0.16 | 0.43 | 0.44 | 3147 |

| | | | | | | | | | | |
|---|---|---|---|---|---|---|---|---|---|---|
| 079 | 5792266 | 2118 | 3367 | 72.49 | 70.42 | 26.76 | 0.08 | 0.01 | 0.00 | 7247 |
| 080 | 3128734 | 1245 | 2943 | 62.78 | 64.79 | 43.66 | 0.08 | 0.02 | 0.01 | 4077 |
| 081 | 5709272 | 1125 | 10355 | 57.80 | 70.42 | 26.76 | 0.06 | 0.05 | 0.01 | 6001 |
| 082 | 5577205 | 1316 | 6874 | 71.38 | 83.10 | 32.39 | 0.06 | 0.01 | 0.00 | 6295 |
| 083 | 5283378 | 1030 | 9475 | 70.63 | 69.01 | 32.39 | 0.07 | 0.02 | 0.00 | 5778 |
| 084 | 2261235 | 265 | 13840 | 73.14 | 91.55 | 5.63 | 0.10 | 0.01 | 0.01 | 2392 |
| 085 | 3682898 | 1079 | 4545 | 65.41 | 50.70 | 14.08 | 0.07 | 0.03 | 0.03 | 3842 |
| 086 | 3074666 | 117 | 42493 | 39.11 | 97.18 | 1.41 | 0.08 | 0.03 | 0.04 | 2741 |
| 087 | 5847895 | 994 | 9481 | 69.31 | 94.37 | 14.08 | 0.08 | 0.03 | 0.01 | 5942 |
| 088 | 2578089 | 976 | 2995 | 69.82 | 40.85 | 9.86 | 0.06 | 0.01 | 0.00 | 3210 |
| 089 | 3563143 | 1664 | 2337 | 63.32 | 69.01 | 21.13 | 0.08 | 0.04 | 0.00 | 4352 |
| 090 | 2598559 | 21 | 170057 | 67.04 | 95.77 | 0.00 | 0.12 | 0.04 | 0.00 | 2334 |
| 091 | 6576024 | 2551 | 3277 | 60.50 | 91.55 | 32.39 | 0.05 | 0.03 | 0.03 | 7841 |
| 092 | 3044225 | 955 | 3959 | 69.89 | 92.96 | 5.63 | 0.06 | 0.03 | 0.03 | 3717 |
| 093 | 2464376 | 1032 | 2622 | 65.26 | 63.38 | 50.70 | 0.06 | 0.05 | 0.02 | 3113 |
| 094 | 6323288 | 1399 | 8343 | 65.75 | 64.79 | 9.86 | 0.06 | 0.03 | 0.00 | 6159 |
| 095 | 6168783 | 2301 | 3349 | 62.18 | 90.14 | 64.79 | 0.05 | 0.03 | 0.01 | 6785 |
| 096 | 2125011 | 468 | 6710 | 65.84 | 95.77 | 4.23 | 0.06 | 0.02 | 0.01 | 2334 |
| 097 | 2821991 | 1438 | 2096 | 71.17 | 39.44 | 2.82 | 0.05 | 0.02 | 0.01 | 3804 |
| 098 | 2994759 | 1384 | 2223 | 59.24 | 36.62 | 8.45 | 0.05 | 0.04 | 0.02 | 4057 |
| 099 | 6977932 | 3497 | 2130 | 63.52 | 47.89 | 11.27 | 0.05 | 0.04 | 0.00 | 8821 |
| 100 | 2774929 | 1278 | 2398 | 66.79 | 66.20 | 57.75 | 0.05 | 0.02 | 0.01 | 3562 |
| 101 | 3233764 | 60 | 74637 | 35.73 | 95.77 | 0.00 | 0.10 | 0.06 | 0.06 | 2843 |
| 102 | 6777504 | 2482 | 3593 | 59.17 | 94.37 | 50.70 | 0.05 | 0.04 | 0.00 | 8119 |
| 103 | 4174659 | 2117 | 2127 | 67.18 | 81.69 | 33.80 | 0.05 | 0.03 | 0.00 | 5111 |
| 104 | 2222925 | 1197 | 1898 | 66.91 | 45.07 | 22.54 | 0.05 | 0.02 | 0.00 | 3137 |
| 105 | 4015081 | 2335 | 1741 | 62.23 | 61.97 | 28.17 | 0.04 | 0.02 | 0.00 | 5295 |
| 106 | 3005903 | 1644 | 1889 | 61.53 | 67.61 | 8.45 | 0.05 | 0.03 | 0.00 | 3923 |
| 107 | 2106025 | 1202 | 1818 | 73.40 | 50.70 | 2.82 | 0.04 | 0.01 | 0.00 | 2854 |
| 108 | 4845334 | 2986 | 1576 | 65.74 | 54.93 | 18.31 | 0.04 | 0.03 | 0.01 | 6673 |
| 109 | 7986166 | 4519 | 1768 | 57.44 | 83.10 | 76.06 | 0.04 | 0.03 | 0.01 | 11251 |
| 110 | 2600002 | 1655 | 1546 | 54.49 | 59.15 | 14.08 | 0.04 | 0.05 | 0.01 | 3839 |
| 111 | 3688075 | 2205 | 1699 | 70.10 | 67.61 | 38.03 | 0.04 | 0.01 | 0.00 | 5152 |
| Sums | 500649046 | 91303 | | | | | 100.00 | 100.00 | 100.00 | 517768 |

**Table 4.** Environmental variables measured throughout the succession experiment. Cell No: equivalent cell number, see materials and methods for details.

| | Sample ID | Days | Cells No (Log10 mL) | Benzene (mM) | Nitrate (mM) | Nitrite (mM) | Nicotinic acid (pM) | Pantothenic acid (pM) | p-Aminobenzoic acid (pM) | Biotin (pM) | Vitamin B12 (pM) |
|---|---|---|---|---|---|---|---|---|---|---|---|
| 1 | B1III | 0 | 4.43 | 0.09 | 8.60 | 0.02 | 150.41 | 152.32 | 467.95 | 147.05 | 25.86 |
| 2 | C1I | 0 | 3.97 | 0.11 | 8.50 | 0.04 | 134.34 | 16.68 | 294.85 | 118.22 | 21.89 |
| 3 | C1III | 0 | 4.23 | 0.09 | 8.38 | 0.04 | 102.20 | 21.37 | 352.55 | 156.76 | 24.40 |
| 4 | A2I | 7 | 4.94 | 0.11 | 8.28 | 0.04 | 150.41 | 39.34 | 64.05 | 49.88 | 19.27 |
| 5 | A2II | 7 | 4.83 | 0.12 | 8.24 | 0.04 | 37.92 | 20.29 | 0.00 | 59.92 | 28.32 |
| 6 | A2III | 7 | 4.85 | 0.11 | 8.22 | 0.04 | 0.00 | 6.93 | 0.00 | 53.12 | 12.49 |
| 7 | A2IV | 7 | 4.80 | 0.10 | 8.22 | 0.04 | 0.00 | 236.42 | 283.88 | 69.31 | 63.64 |
| 8 | B2I | 7 | 5.48 | 0.10 | 8.02 | 0.04 | 0.00 | 14.08 | 563.73 | 92.63 | 92.83 |
| 9 | B2II | 7 | 5.19 | 0.11 | 7.98 | 0.04 | 0.00 | 8.99 | 366.40 | 93.93 | 63.64 |
| 10 | B2III | 7 | 5.40 | 0.11 | 7.96 | 0.04 | 0.00 | 0.00 | 158.68 | 91.66 | 92.83 |
| 11 | B2IV | 7 | 5.62 | 0.11 | 7.96 | 0.04 | 221.11 | 7.54 | 571.81 | 127.94 | 118.52 |
| 12 | C2I | 7 | 5.61 | 0.11 | 7.94 | 0.26 | 0.00 | 0.00 | 778.95 | 119.19 | 107.42 |
| 13 | C2II | 7 | 5.06 | 0.10 | 7.90 | 0.12 | 0.00 | 32.77 | 738.56 | 94.90 | 94.58 |
| 14 | C2III | 7 | 6.05 | 0.09 | 7.90 | 0.12 | 0.00 | 44.76 | 96.94 | 114.66 | 116.77 |
| 15 | C2IV | 7 | 5.39 | 0.11 | 7.84 | 0.16 | 0.00 | 0.00 | 0.00 | 102.03 | 100.42 |
| 16 | A3I | 14 | 5.05 | 0.08 | 7.32 | 0.48 | 0.00 | 4.94 | 0.00 | 45.67 | 5.67 |
| 17 | A3II | 14 | 5.13 | 0.08 | 7.30 | 0.54 | 0.00 | 0.00 | 0.00 | 3.85 | 0.00 |
| 18 | A3III | 14 | 5.19 | 0.08 | 7.26 | 0.56 | 0.00 | 0.00 | 0.00 | 0.00 | 0.00 |
| 19 | A3IV | 14 | 5.14 | 0.08 | 7.22 | 0.56 | 0.00 | 0.00 | 0.00 | 34.01 | 0.00 |
| 20 | B3I | 14 | 5.25 | 0.10 | 7.78 | 0.18 | 0.00 | 0.00 | 0.00 | 0.00 | 0.00 |
| 21 | B3II | 14 | 5.65 | 0.09 | 7.78 | 0.24 | 0.00 | 0.00 | 0.00 | 44.37 | 0.00 |
| 22 | B3III | 14 | 5.01 | 0.11 | 7.70 | 0.06 | 0.00 | 0.00 | 0.00 | 68.67 | 0.00 |
| 23 | B3IV | 14 | 5.26 | 0.10 | 7.68 | 0.28 | 0.00 | 0.00 | 0.00 | 37.25 | 0.00 |
| 24 | C3I | 14 | 5.95 | 0.11 | 7.68 | 0.30 | 0.00 | 0.00 | 0.00 | 36.60 | 0.00 |
| 25 | C3II | 14 | 6.00 | 0.10 | 7.68 | 0.30 | 0.00 | 0.00 | 0.00 | 85.83 | 69.48 |
| 26 | C3III | 14 | 5.93 | 0.06 | 7.20 | 0.56 | 0.00 | 0.00 | 0.00 | 5.73 | 5.39 |
| 27 | C3IV | 14 | 6.08 | 0.11 | 7.66 | 0.34 | 0.00 | 0.00 | 0.00 | 17.26 | 0.00 |
| 28 | A4I | 18 | 5.24 | 0.05 | 7.18 | 0.56 | 0.00 | 0.00 | 0.00 | 0.00 | 0.00 |
| 29 | A4II | 18 | 5.30 | 0.10 | 7.60 | 0.36 | 0.00 | 0.00 | 0.00 | 2.68 | 0.00 |
| 30 | A4III | 18 | 5.22 | 0.05 | 7.16 | 0.56 | 0.00 | 0.00 | 0.00 | 0.00 | 0.00 |
| 31 | A4IV | 18 | 5.35 | 0.07 | 7.14 | 0.56 | 0.00 | 0.00 | 0.00 | 0.00 | 0.00 |
| 32 | B4I | 18 | 5.32 | 0.08 | 7.12 | 0.56 | 0.00 | 0.00 | 0.00 | 0.00 | 0.00 |
| 33 | B4II | 18 | 5.57 | 0.08 | 7.12 | 0.56 | 0.00 | 0.00 | 0.00 | 0.00 | 0.00 |
| 34 | B4III | 18 | 5.35 | 0.10 | 7.56 | 0.42 | 0.00 | 0.00 | 0.00 | 0.00 | 0.00 |
| 35 | B4IV | 18 | 5.22 | 0.08 | 7.12 | 0.56 | 0.00 | 0.00 | 0.00 | 96.84 | 106.84 |
| 36 | C4I | 18 | 5.92 | 0.10 | 7.54 | 0.40 | 0.00 | 0.00 | 0.00 | 7.45 | 0.00 |
| 37 | C4II | 18 | 5.90 | 0.10 | 7.54 | 0.52 | 0.00 | 273.60 | 0.00 | 111.74 | 98.67 |
| 38 | C4III | 18 | 6.18 | 0.10 | 7.52 | 0.44 | 0.00 | 0.00 | 0.00 | 66.72 | 0.00 |

| | | | | | | | | | | | |
|---|---|---|---|---|---|---|---|---|---|---|---|
| 39 | C4IV | 18 | 6.15 | 0.12 | 7.52 | 0.46 | 0.00 | 0.00 | 0.00 | 0.00 | 0.00 |
| 40 | A5I | 22 | 5.64 | 0.09 | 7.50 | 0.46 | 0.00 | 0.00 | 0.00 | 0.00 | 0.00 |
| 41 | A5II | 22 | 5.30 | 0.01 | 6.84 | 0.62 | 0.00 | 0.00 | 877.04 | 45.34 | 77.65 |
| 42 | A5III | 22 | 4.56 | 0.10 | 7.50 | 0.46 | 0.00 | 0.00 | 0.00 | 0.00 | 0.00 |
| 43 | A5IV | 22 | 5.31 | 0.07 | 7.10 | 0.56 | 0.00 | 0.00 | 0.00 | 0.00 | 0.00 |
| 44 | B5I | 22 | 5.87 | 0.04 | 7.12 | 0.58 | 0.00 | 0.00 | 0.00 | 3.43 | 0.00 |
| 45 | B5II | 22 | 5.74 | 0.08 | 7.10 | 0.58 | 0.00 | 0.00 | 0.00 | 0.00 | 0.00 |
| 46 | B5III | 22 | 5.53 | 0.09 | 7.46 | 0.48 | 0.00 | 0.00 | 0.00 | 74.17 | 94.58 |
| 47 | B5IV | 22 | 5.87 | 0.03 | 6.78 | 0.62 | 0.00 | 0.00 | 0.00 | 0.00 | 0.00 |
| 48 | C5I | 22 | 6.15 | 0.08 | 7.08 | 0.56 | 0.00 | 0.00 | 0.00 | 9.39 | 0.00 |
| 49 | C5II | 22 | 5.55 | 0.08 | 7.08 | 0.62 | 0.00 | 0.00 | 0.00 | 0.00 | 0.00 |
| 50 | C5III | 22 | 5.47 | 0.05 | 7.06 | 0.60 | 0.00 | 0.00 | 0.00 | 0.00 | 0.00 |
| 51 | C5IV | 22 | 5.80 | 0.11 | 7.46 | 0.48 | 0.00 | 0.00 | 0.00 | 0.00 | 0.00 |
| 52 | A6I | 26 | 5.50 | 0.01 | 6.78 | 0.62 | 0.00 | 0.00 | 0.00 | 0.00 | 0.00 |
| 53 | A6II | 26 | 5.29 | 0.01 | 6.76 | 0.62 | 0.00 | 0.00 | 0.00 | 0.00 | 0.00 |
| 54 | A6III | 26 | 5.30 | 0.01 | 6.76 | 0.62 | 0.00 | 0.00 | 0.00 | 0.00 | 0.00 |
| 55 | A6IV | 26 | 5.62 | 0.09 | 7.00 | 0.60 | 0.00 | 0.00 | 0.00 | 0.00 | 81.74 |
| 56 | B6I | 26 | 6.22 | 0.03 | 6.76 | 0.62 | 0.00 | 0.00 | 0.00 | 0.00 | 0.00 |
| 57 | B6II | 26 | 6.12 | 0.01 | 6.76 | 0.64 | 0.00 | 0.00 | 0.00 | 0.00 | 0.00 |
| 58 | B6III | 26 | 5.30 | 0.01 | 6.74 | 0.64 | 0.00 | 0.00 | 0.00 | 0.00 | 0.00 |
| 59 | B6IV | 26 | 6.10 | 0.09 | 7.44 | 0.44 | 0.00 | 638.87 | 0.00 | 103.32 | 101.00 |
| 60 | C6I | 26 | 6.27 | 0.01 | 6.74 | 0.64 | 0.00 | 0.00 | 0.00 | 0.00 | 0.00 |
| 61 | C6II | 26 | 6.33 | 0.09 | 6.98 | 0.62 | 0.00 | 0.00 | 0.00 | 0.00 | 0.00 |
| 62 | C6III | 26 | 6.10 | 0.02 | 6.72 | 0.68 | 0.00 | 0.00 | 0.00 | 0.00 | 0.00 |
| 63 | C6IV | 26 | 6.20 | 0.11 | 7.40 | 0.48 | 0.00 | 0.00 | 0.00 | 0.00 | 0.00 |
| 64 | A7I | 30 | 5.16 | 0.01 | 6.72 | 0.64 | 0.00 | 0.00 | 0.00 | 0.00 | 0.00 |
| 65 | A7II | 30 | 5.18 | 0.04 | 6.92 | 0.62 | 0.00 | 0.00 | 0.00 | 0.00 | 0.00 |
| 66 | A7III | 30 | 5.19 | 0.02 | 6.68 | 0.64 | 0.00 | 0.00 | 0.00 | 0.00 | 0.00 |
| 67 | A7IV | 30 | 5.07 | 0.01 | 6.66 | 0.64 | 0.00 | 0.00 | 0.00 | 0.00 | 0.00 |
| 68 | B7I | 30 | 5.99 | 0.10 | 7.40 | 0.50 | 0.00 | 0.00 | 0.00 | 0.00 | 0.00 |
| 69 | B7II | 30 | 5.91 | 0.11 | 7.40 | 0.50 | 1311.23 | 0.00 | 1494.43 | 134.74 | 136.03 |
| 70 | B7III | 30 | 5.85 | 0.01 | 6.62 | 0.64 | 0.00 | 0.00 | 0.00 | 0.00 | 0.00 |
| 71 | B7IV | 30 | 5.52 | 0.07 | 6.88 | 0.62 | 0.00 | 0.00 | 0.00 | 0.00 | 0.00 |
| 72 | C7I | 30 | 6.21 | 0.01 | 6.60 | 0.64 | 0.00 | 0.00 | 0.00 | 0.00 | 0.00 |
| 73 | C7II | 30 | 5.86 | 0.08 | 6.86 | 0.62 | 0.00 | 0.00 | 0.00 | 0.00 | 0.00 |
| 74 | C7III | 30 | 5.86 | 0.10 | 7.34 | 0.50 | 0.00 | 0.00 | 0.00 | 0.00 | 0.00 |
| 75 | C7IV | 30 | 5.50 | 0.01 | 6.58 | 0.64 | 0.00 | 0.00 | 0.00 | 0.00 | 0.00 |
| 76 | A8I | 34 | 5.45 | 0.01 | 6.58 | 0.64 | 0.00 | 0.00 | 0.00 | 0.00 | 0.00 |
| 77 | A8II | 34 | 5.32 | 0.01 | 6.48 | 0.66 | 0.00 | 0.00 | 0.00 | 0.00 | 0.00 |
| 78 | A8III | 34 | 5.33 | 0.06 | 6.86 | 0.60 | 0.00 | 0.00 | 0.00 | 0.00 | 0.00 |
| 79 | A8IV | 34 | 5.43 | 0.01 | 6.48 | 0.64 | 0.00 | 0.00 | 0.00 | 0.00 | 0.00 |
| 80 | B8I | 34 | 6.53 | 0.01 | 6.42 | 0.66 | 0.00 | 0.00 | 0.00 | 0.00 | 0.00 |

| | | | | | | | | | | | |
|---|---|---|---|---|---|---|---|---|---|---|---|
| 81 | B8II | 34 | 6.25 | 0.01 | 6.38 | 0.66 | 0.00 | 0.00 | 0.00 | 0.00 | 0.00 |
| 82 | B8III | 34 | 6.24 | 0.01 | 6.32 | 0.78 | 0.00 | 0.00 | 0.00 | 0.00 | 0.00 |
| 83 | B8IV | 34 | 6.02 | 0.11 | 7.34 | 0.50 | 0.00 | 0.00 | 0.00 | 121.46 | 129.61 |
| 84 | C8I | 34 | 6.15 | 0.01 | 6.28 | 0.72 | 0.00 | 0.00 | 0.00 | 0.00 | 0.00 |
| 85 | C8II | 34 | 6.05 | 0.13 | 7.32 | 0.52 | 0.00 | 0.00 | 0.00 | 125.67 | 122.60 |
| 86 | C8III | 34 | 5.92 | 0.01 | 6.20 | 0.72 | 0.00 | 0.00 | 0.00 | 0.00 | 0.00 |
| 87 | C8IV | 34 | 6.12 | 0.01 | 6.20 | 0.66 | 0.00 | 0.00 | 0.00 | 0.00 | 0.00 |

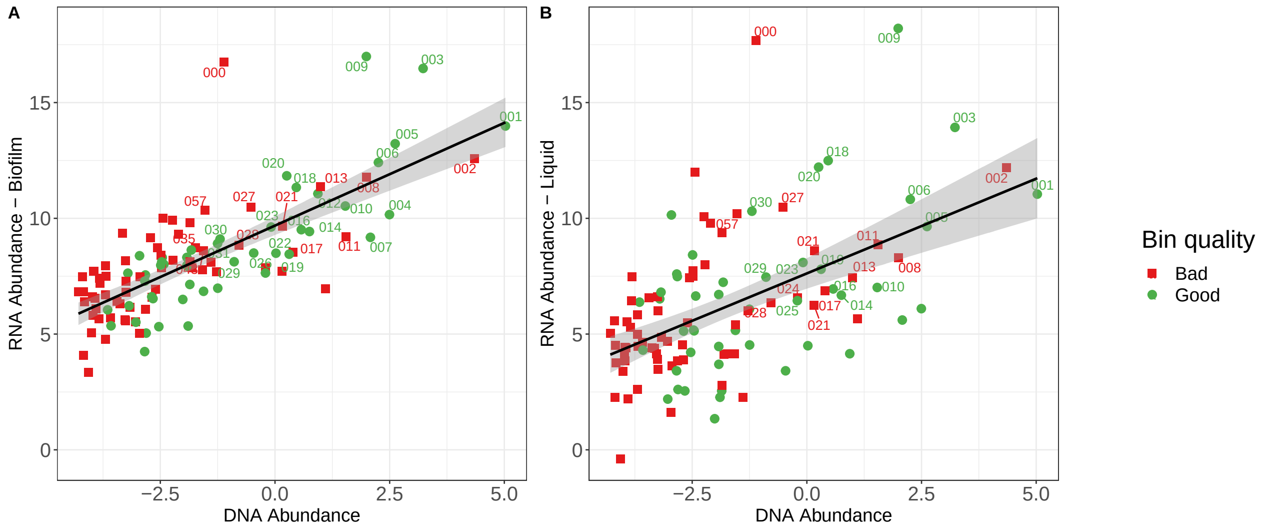


**Figure 1.** Relation of MAGs RNA abundance with DNA abundance (biofilm samples). A) for RNA derived from biofilm. B) for RNA derived from liquid. The color of the points indicates the quality of MAGs. The positive correlation varying from moderate to weak, respectively (biofilm: $r^2 = 0.556$, liquid: $r^2 = 0.286$). The most transcribed MAGs, 3 and 9, were not the most abundant ones as judged by their DNA

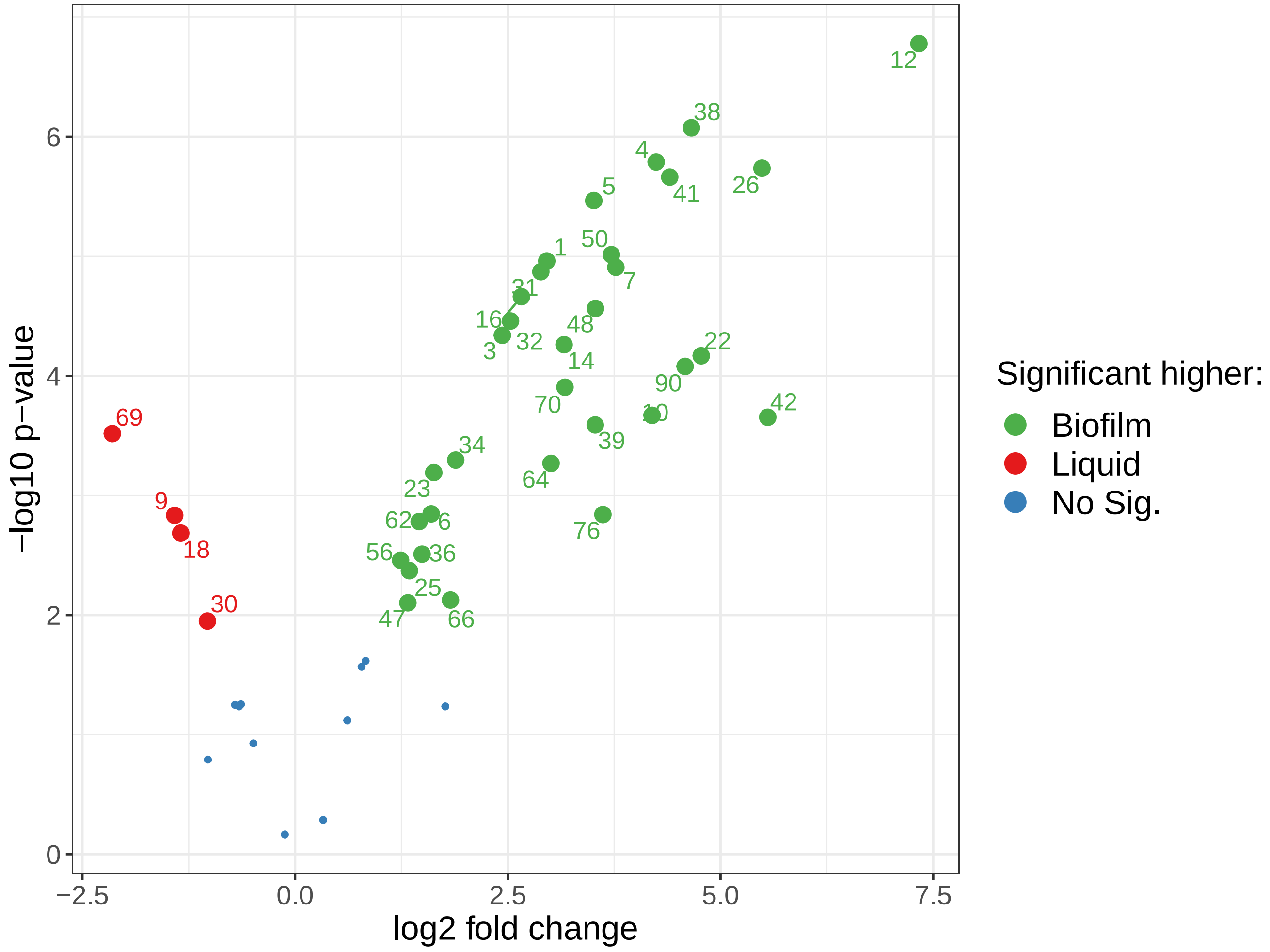


**Figure 2.** Volcano plot indicate the significant different RNA content of MAGs between Between biofilm and liquid phase. Right side with green color the MAGs are higher in biofilm samples, while the left side with red indicated the liquid phase.


[4] S. Andrews. *Trim Galore, a wrapper tool around Cutadapt and FastQC to consistently apply quality and adapter trimming to FastQ files. Available at* http://www.bioinformatics.babraham.ac.uk/projects/trim_galore. 2012. URL: http://www.bioinformatics.babraham.ac.uk/projects/trim_galore/.

[5] Siavash Atashgahi et al. "A benzene-degrading nitrate-reducing microbial consortium displays aerobic and anaerobic benzene degradation pathways". In: *Scientific Reports* 8.1 (Mar. 2018). DOI: 10.1038/s41598-018-22617-x. URL: https://doi.org/10.1038/s41598-018-22617-x.

[6] Etienne Becht et al. "Dimensionality reduction for visualizing single-cell data using UMAP". In: *Nature Biotechnology* 37.1 (Dec. 2018), pp. 38–44. DOI: 10.1038/nbt.4314. URL: https://doi.org/10.1038/nbt.4314.

[7] U. Bodenhofer, A. Kothmeier, and S. Hochreiter. "APCluster: an R package for affinity propagation clustering". In: *Bioinformatics* 27.17 (July 2011), pp. 2463–2464. ISSN: 1460-2059. DOI: 10.1093/bioinformatics/btr406. URL: http://dx.doi.org/10.1093/bioinformatics/btr406.

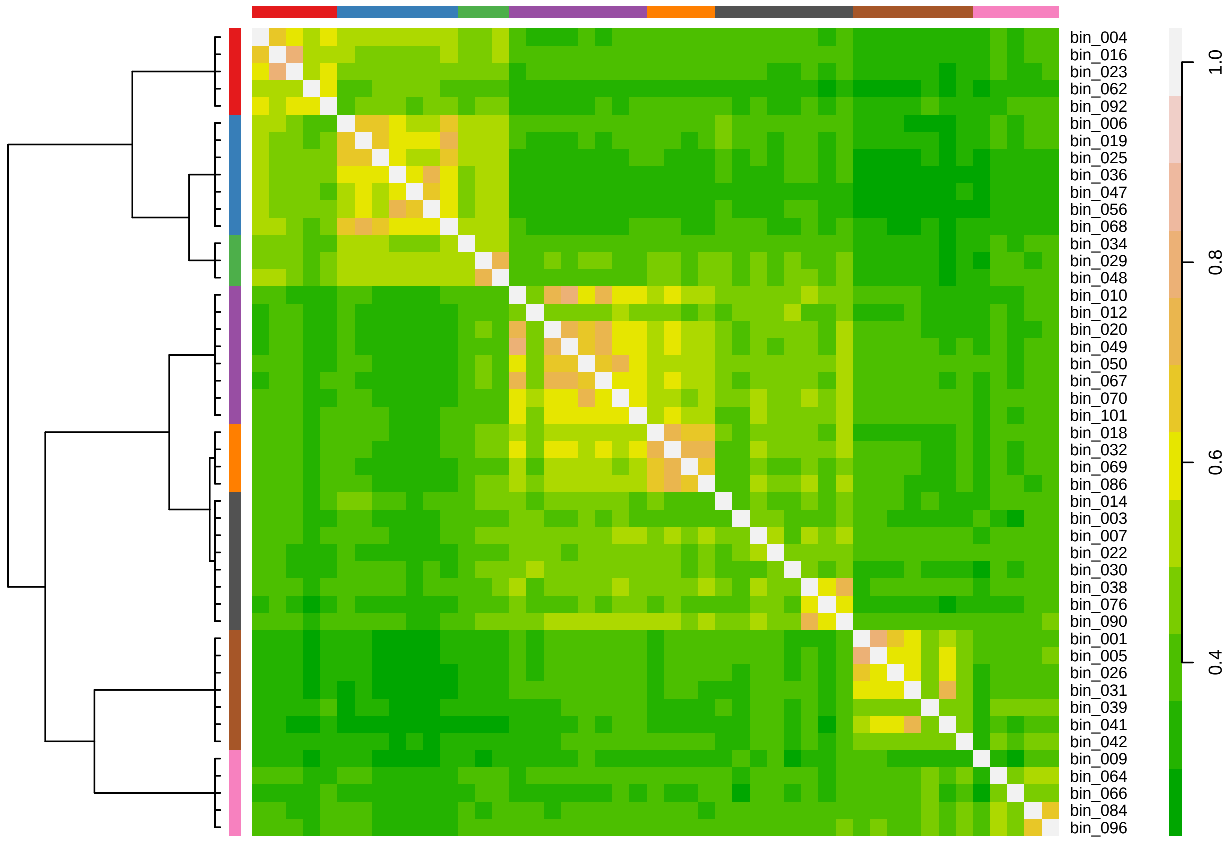


**Figure 3.** Affinity propagation heatmap on clustered MAGs. 8 clusters were determined, which could further assigned on 3 group as the dendrogram on the left side indicates

[8] Ulrich Bodenhofer et al. "msa: an R package for multiple sequence alignment". In: *Bioinformatics* (Aug. 2015), btv494. DOI: 10.1093/bioinformatics/btv494. URL: https://doi.org/10.1093/bioinformatics/btv494.

[9] Matthias Boll et al. "Anaerobic degradation of homocyclic aromatic compounds via arylcarboxyl-coenzyme A esters: organisms, strategies and key enzymes". In: *Environmental Microbiology* 16.3 (Dec. 2013), pp. 612–627. DOI: 10.1111/1462-2920.12328. URL: https://doi.org/10.1111/1462-2920.12328.

[10] Leo Breiman. "Random Forests". In: *Machine Learning* 45.1 (2001), pp. 5–32. ISSN: 0885-6125. DOI: 10.1023/a:1010933404324. URL: http://dx.doi.org/10.1023/A:1010933404324.

[11] J. G. Caporaso et al. "Global patterns of 16S rRNA diversity at a depth of millions of sequences per sample". In: *Proceedings of the National Academy of Sciences* 108.Supplement_1 (June 2010), pp. 4516–4522. DOI: 10.1073/pnas.1000080107. URL: https://doi.org/10.1073/pnas.1000080107.

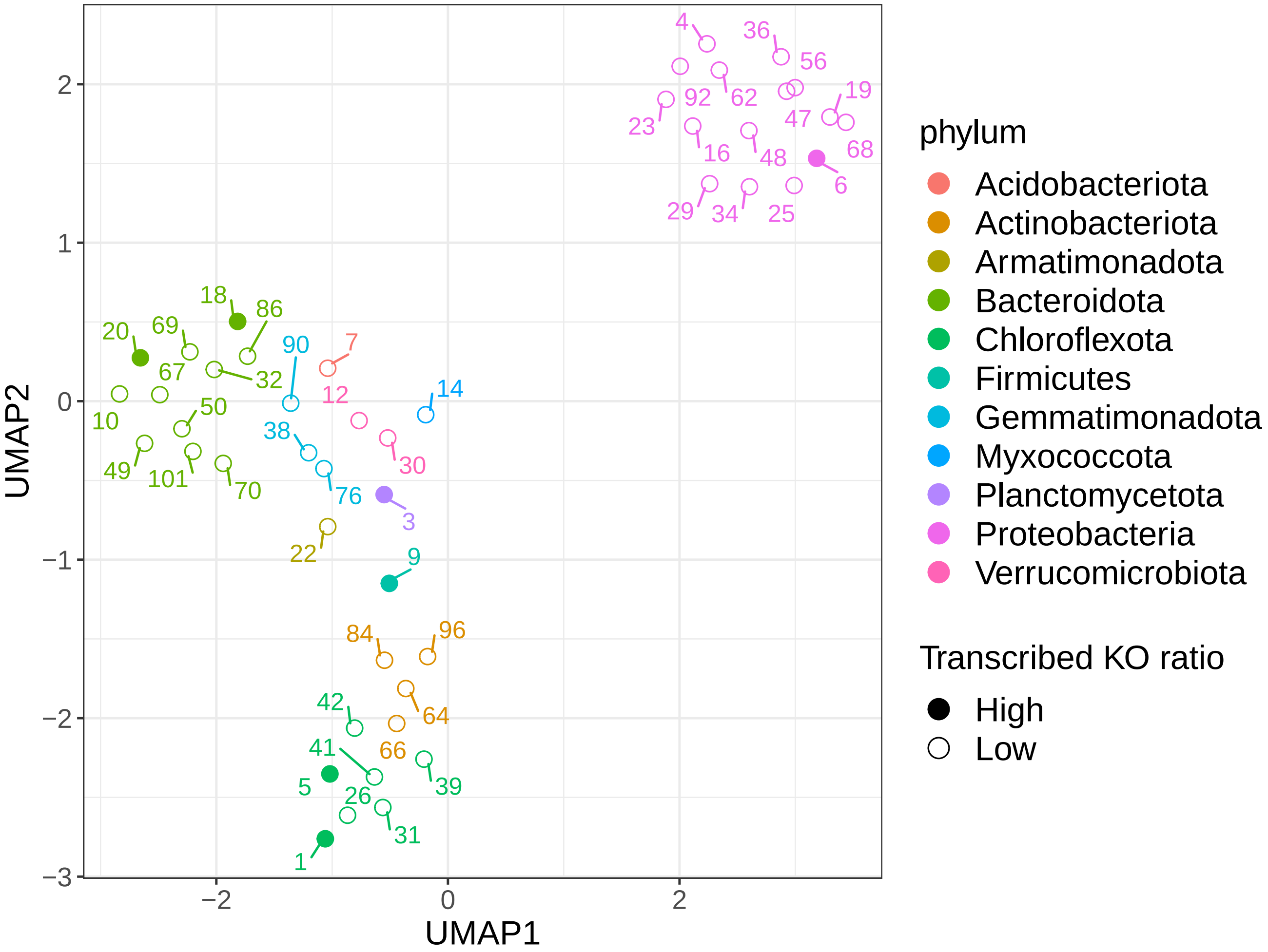


**Figure 4.** Functional landscape with UMAP. Colored based on phylum taxonomic assignment.

[12] Andrea Carboni et al. “A method for the determination of fullerenes in soil and sediment matrices using ultra-high performance liquid chromatography coupled with heated electrospray quadrupole time of flight mass spectrometry”. In: *Journal of Chromatography A* 1433 (Feb. 2016), pp. 123–130. DOI: 10.1016/j.chroma.2016.01.035. URL: https://doi.org/10.1016/j.chroma.2016.01.035.

[13] Manuel Carmona et al. “Anaerobic Catabolism of Aromatic Compounds: a Genetic and Genomic View”. In: *Microbiology and Molecular Biology Reviews* 73.1 (Mar. 2009), pp. 71–133. DOI: 10.1128/mmbr.00021-08. URL: https://doi.org/10.1128/mmbr.00021-08.

[14] Pierre-Alain Chaumeil et al. “GTDB-Tk: a toolkit to classify genomes with the Genome Taxonomy Database”. In: *Bioinformatics* (Nov. 2019). Ed. by John Hancock. DOI: 10.1093/bioinformatics/btz848. URL: https://doi.org/10.1093/bioinformatics/btz848.

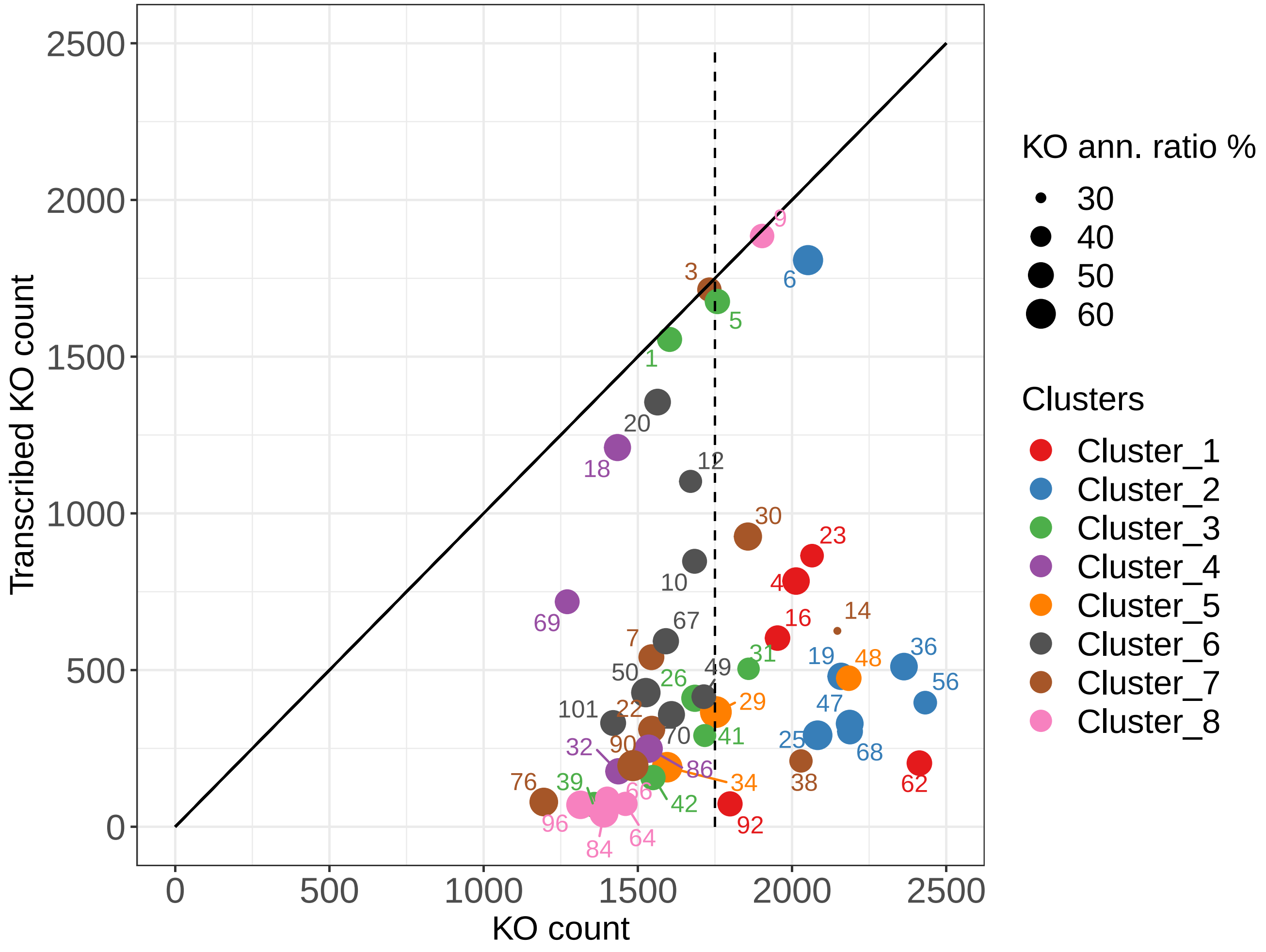


**Figure 5.** Relationship between MAGs transcribed KOs and the total KOs count. Size of the point indicates the KO annotation ratio and color the clustering result. t-test on KO count: (group C (Mean = 2082.6, SD = 238.7) and group (A + B) (Mean = 1593.3 %, SD = 214.8); p-value = 4.381e-07). t-test on KO annotation ratio: (group C (Mean = 53.3 %, SD = 6.2) and group (A + B) (Mean = 50 %, SD = 6.1); p-value = 0.099)

**Table 2.** Taxonomy assignment overview of the selected MAGs.The first column indicates the number of the MAG. Columns 2-4 is the GTDB taxonomy, keep in mind the recent change proposed in [46] in comparision to NCBI taxonomy. Columns 5-6 are the significant sequence matches (>90%) between the 16S RNA genes of MAGs from the bioreactor and the succession experiment. Columns 7-8 are the taxonomy of significant sequence matches (>97%) between the 16S RNA genes of MAGs from the bioreactor and the Silva database, 132 SSU ref. Nr 99 version (The taxonomic levels of domain and phylum were matched and therefore excluded).
*U = uncultured

| | **GTDB Taxonomy of MAGs** | | | **Between experiments** | | **Silva (version 132 SSU ref. Nr99) taxonomy of succession experiment** | |
|---|---|---|---|---|---|---|---|
| MAGs | Phylum | Class | Genus | OTUs | Identity | Identity | Bacteria taxonomy |
| 001 | Chloroflexota | Anaerolineae | UBA7227 | | | 100.00 | Anaerolineae;Anaerolineales;Anaerolineaceae;U; |
| 003 | Planctomycetota | Brocadiae | Kuenenia | | | 99.68 | Brocadiae;Brocadiales;Brocadiaceae;Candidatus Kuenenia; |
| 004 | Proteobacteria | Alphaproteobacteria | Pseudorhodoplanes | OTU525650078 | 93.70 | 100.00 | Alphaproteobacteria;Rhizobiales;Xanthobacteraceae;U; |
| 005 | Chloroflexota | Anaerolineae | OLB14 | | | 97.63 | Anaerolineae;Anaerolineales;Anaerolineaceae;U; |
| 006 | Proteobacteria | Gammaproteobacteria | UTPRO2 | | | | |
| 007 | Acidobacteriota | Blastocatellia | OLB17 | | | 98.81 | Blastocatellia (Subgroup 4);Blastocatellales;Blastocatellaceae;OLB17; |
| 009 | Firmicutes | Thermincolia | | OTU624837510 | 100.00 | 97.17 | Clostridia;Clostridiales;*Peptococcaceae*;Thermincola; |
| 010 | Bacteroidota | Ignavibacteria | Ignavibacterium | | | 98.59 | Ignavibacteria;Ignavibacteriales;PHOS-HE36; |
| 012 | Verrucomicrobiota | Verrucomicrobiae | | OTU226057524 | 90.35 | | |
| 014 | Myxococcota | Polyangia | | | | | |
| 016 | Proteobacteria | Alphaproteobacteria | Hyphomicrobium | | | | |
| 018 | Bacteroidota | Bacteroidia | | OTU942897628 | 100.00 | 98.34 | Bacteroidia;Flavobacteriales;Cryomorphaceae;U; |
| 019 | Proteobacteria | Gammaproteobacteria | SCN-69-89 | | | | |
| 020 | Bacteroidota | Ignavibacteria | | | | | |
| 022 | Armatimonadota | Fimbriimonadia | | OTU276191148 | 100.00 | | |
| 023 | Proteobacteria | Alphaproteobacteria | Hyphomicrobium | | | | |
| 025 | Proteobacteria | Gammaproteobacteria | PALSA-1003 | | | | |
| 026 | Chloroflexota | Anaerolineae | Promineofilum | | | 97.75 | Anaerolineae;Ardenticatenales;U; |
| 029 | Proteobacteria | Gammaproteobacteria | | OTU826979979 | 100.00 | 98.37 | Gammaproteobacteria;Xanthomonadales;Xanthomonadaceae;Arenimonas; |
| 030 | Verrucomicrobiota | Verrucomicrobiae | Didemnitutus | | | 100.00 | Verrucomicrobiae;Opitutales;Opitutaceae;Lacunisphaera; |
| 031 | Chloroflexota | Anaerolineae | OLB15 | | | | |
| 032 | Bacteroidota | Bacteroidia | OLB10 | | | 98.16 | Bacteroidia;Sphingobacteriales;AKYH767; |
| 034 | Proteobacteria | Gammaproteobacteria | | | | | |
| 036 | Proteobacteria | Gammaproteobacteria | Hydrogenophaga | | | | |
| 038 | Gemmatimonadota | Gemmatimonadetes | SCN-70-22 | OTU170800226 | 100.00 | 99.46 | Gemmatimonadetes;Gemmatimonadales;Gemmatimonadaceae;U; |
| 039 | Chloroflexota | Ellin6529 | Palsa-1032 | | | 97.13 | Chloroflexi;KD4-96; |
| 041 | Chloroflexota | Anaerolineae | ZC4RG36 | | | | |
| 042 | Chloroflexota | Chloroflexia | | | | | |
| 047 | Proteobacteria | Gammaproteobacteria | Pusillimonas | OTU607377144 | 95.75 | 98.86 | Gammaproteobacteria;Betaproteobacteriales;Burkholderiaceae;Eoetvoesia; |
| 048 | Proteobacteria | Gammaproteobacteria | Dokdonella | | | | |
| 049 | Bacteroidota | Ignavibacteria | | | | | |
| 050 | Bacteroidota | UBA10030 | 2-02-FULL-55-14 | | | | |
| 056 | Proteobacteria | Gammaproteobacteria | Comamonas | | | 97.49 | Gammaproteobacteria;Betaproteobacteriales;Burkholderiaceae;Comamonas; |
| 062 | Proteobacteria | Alphaproteobacteria | Ochrobactrum | | | 97.82 | Alphaproteobacteria;Rhodobacterales;Rhodobacteraceae;Defluviimonas; |
| 064 | Actinobacteriota | Acidimicrobiia | | | | | |
| 066 | Actinobacteriota | Actinobacteria | Cryobacterium | OTU167856346 | 96.71 | 98.42 | Actinobacteria;Micrococcales;Microbacteriaceae;Leifsonia; |
| 067 | Bacteroidota | Ignavibacteria | UTCHB3 | | | | |
| 068 | Proteobacteria | Gammaproteobacteria | SCN-69-89 | | | 98.86 | Gammaproteobacteria;Betaproteobacteriales;Rhodocyclaceae;Azoarcus; |
| 069 | Bacteroidota | Bacteroidia | | | | | |
| 070 | Bacteroidota | UBA10030 | UBA6688 | | | | |
| 076 | Gemmatimonadota | Gemmatimonadetes | | | | | |
| 084 | Actinobacteriota | Thermoleophilia | | | | | |
| 086 | Bacteroidota | Bacteroidia | Ferruginibacter | OTU171842870 | 100.00 | 97.28 | Bacteroidia;Chitinophagales;Chitinophagaceae;Ferruginibacter; |
| 090 | Gemmatimonadota | Gemmatimonadetes | Fen-1231 | OTU758662476 | 100.00 | | |
| 092 | Proteobacteria | Alphaproteobacteria | | OTU623980803 | 100.00 | 97.04 | Alphaproteobacteria;Rhodobacterales;Rhodobacteraceae;Defluviimonas; |
| 096 | Actinobacteriota | Thermoleophilia | 67-14 | | | 97.91 | Thermoleophilia;Solirubrobacterales;67-14; |
| 101 | Bacteroidota | Ignavibacteria | | | | | |

**Table 3.** Empirical Bayes statistics for differential expression MAGs

| MAGs | logFC | AveExpr | t-statistic (t) | P.Value | adj.P.Val | log-odds |
|---|---|---|---|---|---|---|
| 012 | 7.3325 | 9.5170 | 20.3567 | 0.0000 | 0.0000 | 8.1131 |
| 038 | 4.6581 | 11.0428 | 16.0428 | 0.0000 | 0.0000 | 6.6907 |
| 004 | 4.2434 | 11.3044 | 14.5723 | 0.0000 | 0.0000 | 6.0784 |
| 026 | 5.4871 | 10.7020 | 14.3695 | 0.0000 | 0.0000 | 5.9877 |
| 041 | 4.4030 | 11.1860 | 13.9812 | 0.0000 | 0.0000 | 5.8094 |
| 005 | 3.5107 | 11.5273 | 13.0433 | 0.0000 | 0.0000 | 5.3520 |
| 050 | 3.7171 | 11.4175 | 11.2162 | 0.0000 | 0.0001 | 4.3371 |
| 001 | 2.9578 | 11.8363 | 10.9766 | 0.0000 | 0.0001 | 4.1902 |
| 007 | 3.7702 | 11.4672 | 10.8237 | 0.0000 | 0.0001 | 4.0945 |
| 031 | 2.8883 | 11.8763 | 10.6430 | 0.0000 | 0.0001 | 3.9796 |
| 016 | 2.6599 | 12.0536 | 9.8980 | 0.0000 | 0.0001 | 3.4827 |
| 048 | 3.5305 | 11.8638 | 9.6105 | 0.0000 | 0.0001 | 3.2802 |
| 032 | 2.5321 | 12.0779 | 9.2221 | 0.0000 | 0.0001 | 2.9968 |
| 003 | 2.4361 | 11.9092 | 8.8423 | 0.0000 | 0.0001 | 2.7079 |
| 014 | 3.1613 | 12.0269 | 8.6381 | 0.0001 | 0.0002 | 2.5475 |
| 022 | 4.7732 | 10.9894 | 8.3853 | 0.0001 | 0.0002 | 2.3438 |
| 090 | 4.5838 | 10.9320 | 8.1231 | 0.0001 | 0.0002 | 2.1263 |
| 042 | 5.1844 | 10.5614 | 7.7161 | 0.0001 | 0.0003 | 1.7755 |
| 070 | 3.1721 | 11.5333 | 7.6134 | 0.0001 | 0.0003 | 1.6843 |
| 010 | 4.1957 | 11.2011 | 7.0016 | 0.0002 | 0.0005 | 1.1179 |
| 039 | 3.5275 | 11.5724 | 6.7917 | 0.0002 | 0.0005 | 0.9139 |
| 069 | -2.1483 | 12.5489 | -6.5826 | 0.0003 | 0.0006 | 0.7053 |
| 034 | 1.8881 | 12.4637 | 6.0496 | 0.0005 | 0.0010 | 0.1494 |
| 064 | 3.0082 | 11.7744 | 6.0140 | 0.0005 | 0.0010 | 0.1109 |
| 023 | 1.6300 | 12.3365 | 5.7996 | 0.0006 | 0.0012 | -0.1242 |
| 076 | 3.6178 | 11.9505 | 5.0890 | 0.0013 | 0.0024 | -0.9494 |
| 006 | 1.5996 | 12.5047 | 5.0697 | 0.0014 | 0.0024 | -0.9728 |
| 009 | -1.4153 | 12.6498 | -5.0349 | 0.0014 | 0.0024 | -1.0152 |
| 062 | 1.4584 | 12.4299 | 4.9322 | 0.0016 | 0.0026 | -1.1413 |
| 018 | -1.3443 | 12.6528 | -4.7360 | 0.0020 | 0.0032 | -1.3867 |
| 036 | 1.4923 | 12.6101 | 4.4083 | 0.0030 | 0.0045 | -1.8093 |
| 056 | 1.2397 | 12.6138 | 4.3022 | 0.0034 | 0.0050 | -1.9496 |
| 025 | 1.3439 | 12.5750 | 4.1487 | 0.0041 | 0.0059 | -2.1555 |
| 066 | 1.8263 | 12.5122 | 3.7248 | 0.0072 | 0.0099 | -2.7415 |
| 047 | 1.3257 | 12.5873 | 3.6746 | 0.0077 | 0.0103 | -2.8125 |
| 030 | -1.0303 | 12.8495 | -3.4048 | 0.0111 | 0.0145 | -3.1998 |
| 029 | 0.8291 | 12.7212 | 2.8561 | 0.0240 | 0.0305 | -4.0099 |
| 019 | 0.7818 | 12.7347 | 2.7753 | 0.0270 | 0.0334 | -4.1310 |
| 067 | -0.6359 | 12.7577 | -2.2820 | 0.0558 | 0.0647 | -4.8695 |
| 020 | -0.7065 | 12.6776 | -2.2794 | 0.0560 | 0.0647 | -4.8734 |
| 049 | 1.7660 | 12.0417 | 2.2745 | 0.0565 | 0.0647 | -4.8806 |
| 086 | -0.6579 | 12.8018 | -2.2575 | 0.0579 | 0.0648 | -4.9058 |
| 084 | 0.6140 | 12.4948 | 2.0753 | 0.0759 | 0.0830 | -5.1740 |
| 092 | -0.4895 | 12.7721 | -1.7739 | 0.1186 | 0.1267 | -5.6041 |
| 101 | -1.0248 | 12.2585 | -1.5715 | 0.1593 | 0.1664 | -5.8784 |
| 096 | 0.3304 | 12.3395 | 0.6851 | 0.5149 | 0.5261 | -6.8202 |
| 068 | -0.1201 | 12.7129 | -0.4236 | 0.6844 | 0.6844 | -6.9773 |

**Table 5.** Ionization modes, mass transitions, operating parameters, limits of quantification, and chromatographic retention times for the measured vitamins.

| Compound | Ionization mode | Precursor ion > product ion (m/z) | Declustering potential (V) | Collision energy (V) | Collision exit potential (V) | LOQ (ppm) * | Chromatographic retention time (min) |
|---|---|---|---|---|---|---|---|
| 4-Aminobenzoic acid | negative | 136 $[M-H]^-$ > 92 | -52 | -16 | -3 | 0.004 | 7.2 |
| Biotin | positive | 245 $[M+H]^+$ > 227 | 63 | 21 | 11 | 0.002 | 8.2 |
| Lipoic acid | negative | 205 $[M-H]^-$ > 171 | -40 | -11 | -10 | 0.016 | 9.5 |
| Nicotinic acid | positive | 124 $[M+H]^+$ > 80 | 75 | 30 | 12 | 0.016 | 2.4 |
| Pantothenic acid | positive | 220 $[M+H]^+$ > 90 | 72 | 23 | 5 | 0.004 | 6.3 |
| Riboflavin | positive | 377 $[M+H]^+$ > 243 | 96 | 33 | 13 | 0.032 | 8.5 |
| Vitamin $B_{12}$ | positive | 700 $[M+2Na]^+$ > 658 | 113 | 39 | 33 | 0.016 | 7.6 |

*LOQ: Limit of quantification (defined as the concentration of the lowest standard with a signal/noise ratio >10 and an accuracy of 70-130%).

**Table 6.** Mass transitions, limits of detection, and chromatographic retention times for compounds used for data-independent target analysis.

| Compound | Precursor ion > product ion (m/z) | LOD (ppm) * | Chromatographic retention time (min) |
|---|---|---|---|
| 4-Hydroxybenzoate | 137 $[M-H]^-$ > 93 | 0.01 | 10.1 |
| Benzoate | 121 $[M-H]^-$ > 72 | 0.01 | 12.6 |
| Benzylsuccinate | 207 $[M-H]^-$ > 163 | 0.01 | 12.9 |

*Limit of detection (defined as the concentration of the lowest standard with an identifiable chromatographic peak of the precursor ion and, if applicable, product ion).

**Table 7.** Mass ranges used for data-dependent auto-MS/MS analysis of suspected metabolites.

| Compound | Targeted mass range (m/z) |
|---|---|
| 2-Hydroxy-cyclohexanecarboxylate | 143.01-143.21 |
| 2-Oxo-cyclohexanecarboxylate | 141.96-142.16 |
| Catechol | 108.93-109.13 |

**Table 8.** Definitions used in this paper.

| | **Defined as a niche that:** |
|---|---|
| **Primary consumer** | Feeds from the main free energy source and as such produces the initial biomass |
| **Syntrophy** | Depends on another member by cross-feed of compounds |
| **Scavenging** | Feeds from metabolic left-overs or on the contents of dead cells of others with a return to the public-good |
| **Cheating** | Feeds from metabolic left-overs of others without a return to the public-good |
| **Predation** | Actively attacks and feeds on other member of the community |

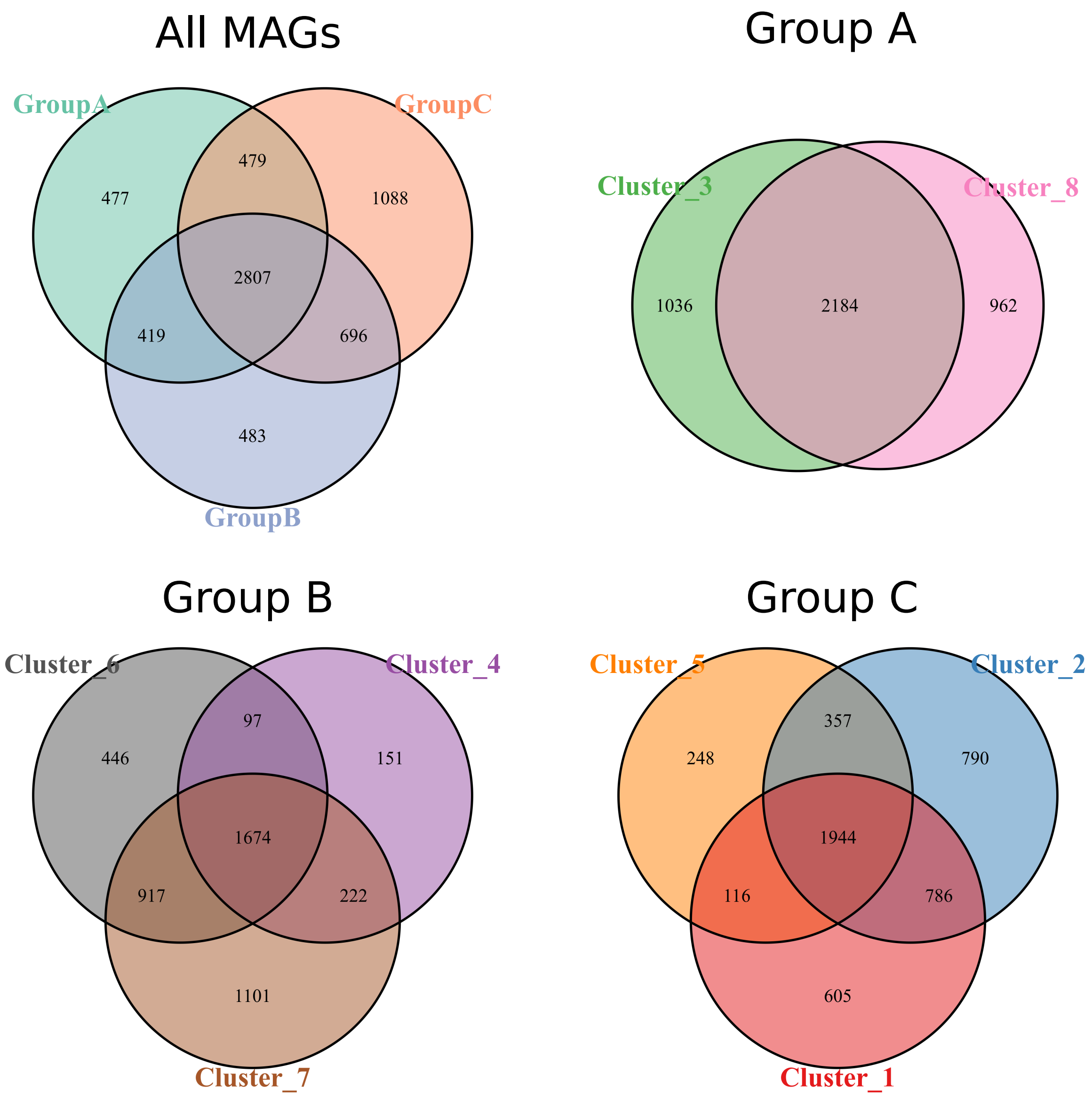


**Figure 6.** Venn diagram based on KO Ids. All MAGs separated on three groups (top left), MAGs of group A separated on two clusters (top right), MAGs of group B separated on three clusters (bottom left), MAGs of group C separated on three clusters (bottom right).

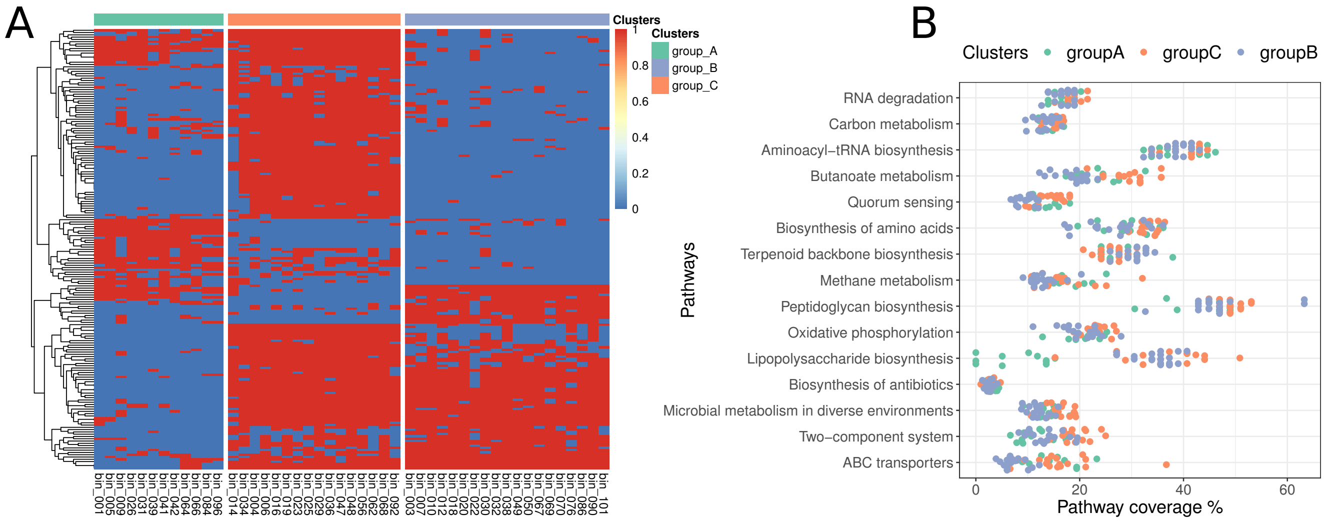


**Figure 7.** Feature selection result to discriminate the three different MAGs groups. A) An overview heatmap for the presence/absence (red/blue) KO that discriminate the 3 functional group of MAGs. B) The pathway coverage % of KEGG pathways that discriminate the 3 functional group of MAGs. The major discriminator involves genes for lipopolysaccharide biosynthesis, which were found specifically more in MAGs from group C and B, as opposed to those from group A

**Table 9.** Taxonomy assignment of more significant correlated OTUs of the succession experiment with cultures stage, selection is based on random forest variable importance and PERMANOVA analysis

| OTUs | Phylum | Class | Order | Family | Genus (Species) |
|---|---|---|---|---|---|
| OTU851020915 | Proteobacteria | Alphaproteobacteria | Rhizobiales | Brucellaceae | Ochrobactrum |
| OTU501058114 | Proteobacteria | Gammaproteobacteria | Pseudomonadales | Pseudomonadaceae | Pseudomonas (stutzeri) |
| OTU474979920 | Proteobacteria | Betaproteobacteria | Rhodocyclales | Rhodocyclaceae | Azoarcus |
| OTU152847281 | Proteobacteria | Gammaproteobacteria | Xanthomonadales | Xanthomonadaceae | Thermomonas |
| OTU596667673 | Proteobacteria | Gammaproteobacteria | Enterobacteriales | Enterobacteriaceae | Escherichia-Shigella |
| OTU819133079 | Bacteroidetes | Flavobacteria | Flavobacteriales | Flavobacteriaceae | Chryseobacterium |
| OTU888962565 | Proteobacteria | Betaproteobacteria | Burkholderiales | Oxalobacteraceae | Undibacterium |
| OTU27733144 | Proteobacteria | Betaproteobacteria | Burkholderiales | Comamonadaceae | Comamonas |
| OTU344128085 | Proteobacteria | Betaproteobacteria | Burkholderiales | Comamonadaceae | unclassified_Comamonadaceae |
| OTU32483051 | Proteobacteria | Alphaproteobacteria | Rhizobiales | Rhodobiaceae | unclassified_Rhodobiaceae |
| OTU121064001 | Proteobacteria | Betaproteobacteria | Burkholderiales | Burkholderiaceae | Limnobacter |
| OTU858316386 | Firmicutes | Clostridia | Clostridiales | *Peptococcaceae* | Thermincola |
| OTU758662476 | Gemmatimonadetes | Gemmatimonadetes | Gemmatimonadales | Gemmatimonadaceae | Gemmatimonas |
| OTU244476266 | unclassified_Bacteria | unclassified_Bacteria | unclassified_Bacteria | unclassified_Bacteria | unclassified_Bacteria |
| OTU575006217 | unclassified_Bacteria | unclassified_Bacteria | unclassified_Bacteria | unclassified_Bacteria | unclassified_Bacteria |
| OTU101622224 | Verrucomicrobia | Opitutae | Opitutales | Opitutaceae | Opitutus |

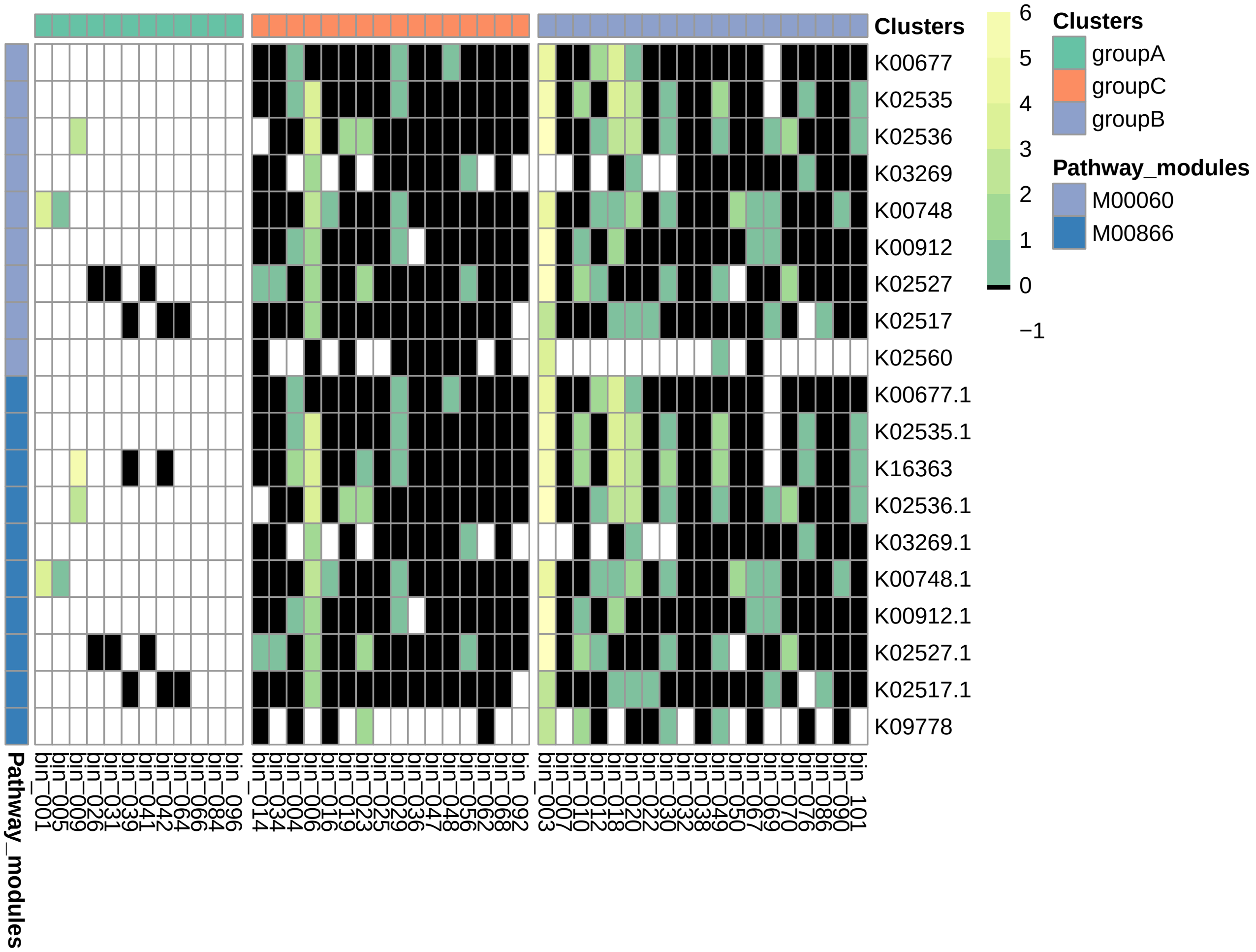


**Figure 8.** Heatmap of lipid A biosynthesis, which is involved in lipopolysaccharide biosynthesis. The presence/absence is represented by black/white colors and the strength of transcription level is represented by the color gradient. KEGG module-M00060: LpxL-LpxM type, KEGG module-M00866: non-LpxL-LpxM type. More specifically those genes encoding a hydrophobic anchor called lipid A, an essential molecule for membrane synthesis, which are discriminators for MAGs from group A compared to group C and B

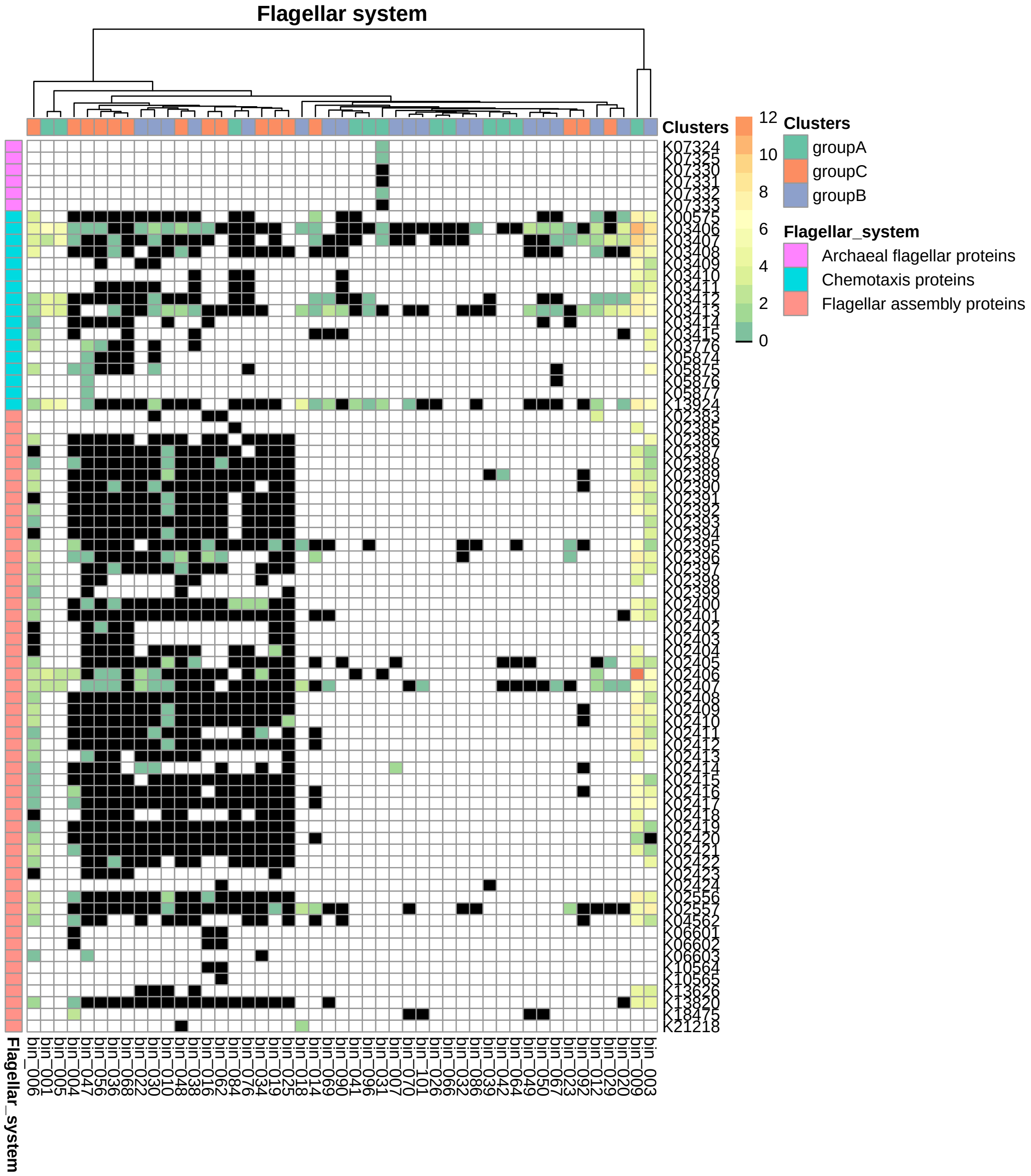


**Figure 9.** Heatmap of flagellar system, with the presence and absence (black/white) of the KOs and color scale indicating the strength of transcription.

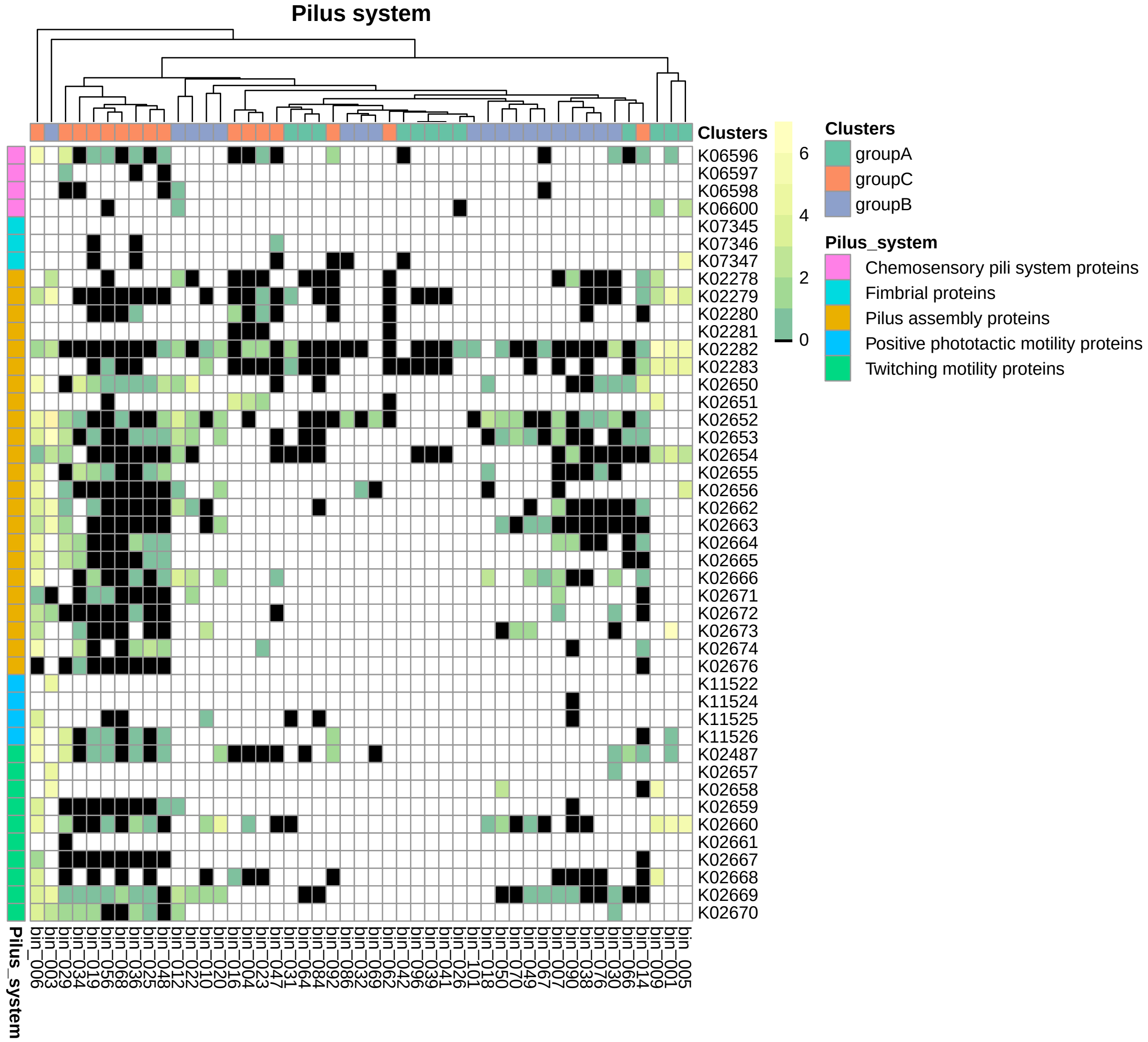


**Figure 10.** Heatmap of pilus system, with the presence and absence (black/white) of the KOs and color scale indicating the strength of transcription.

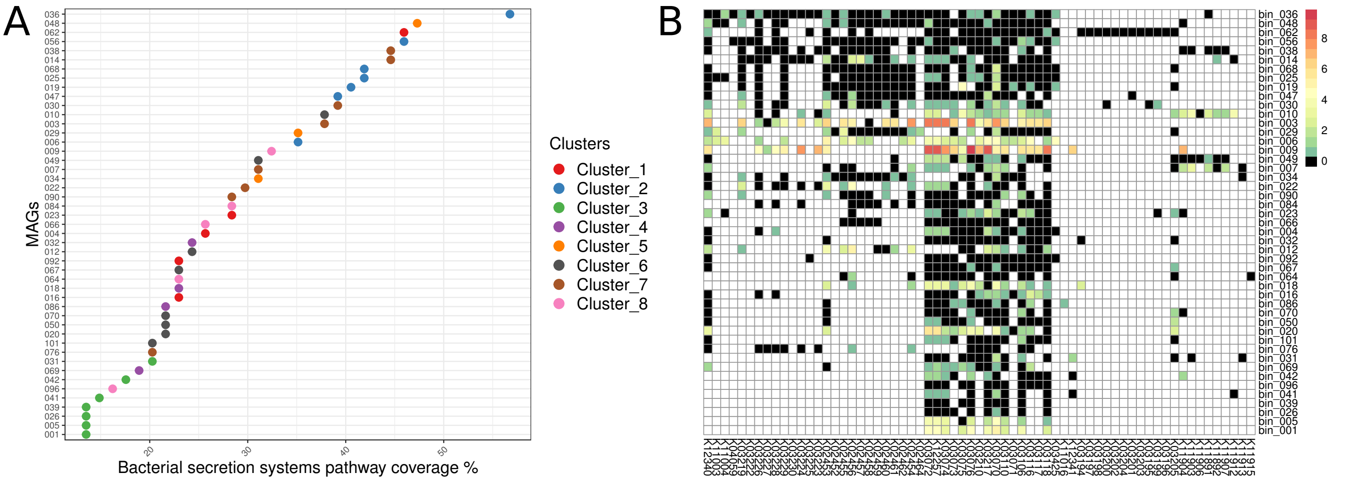


**Figure 11.** A) The pathway coverage % of MAGs on bacteria secretion systems and B) the heatmap of MAGs with the corresponding KOs, with the presence and absence (black/white) of the KOs and color scale indicating the strength of transcription.

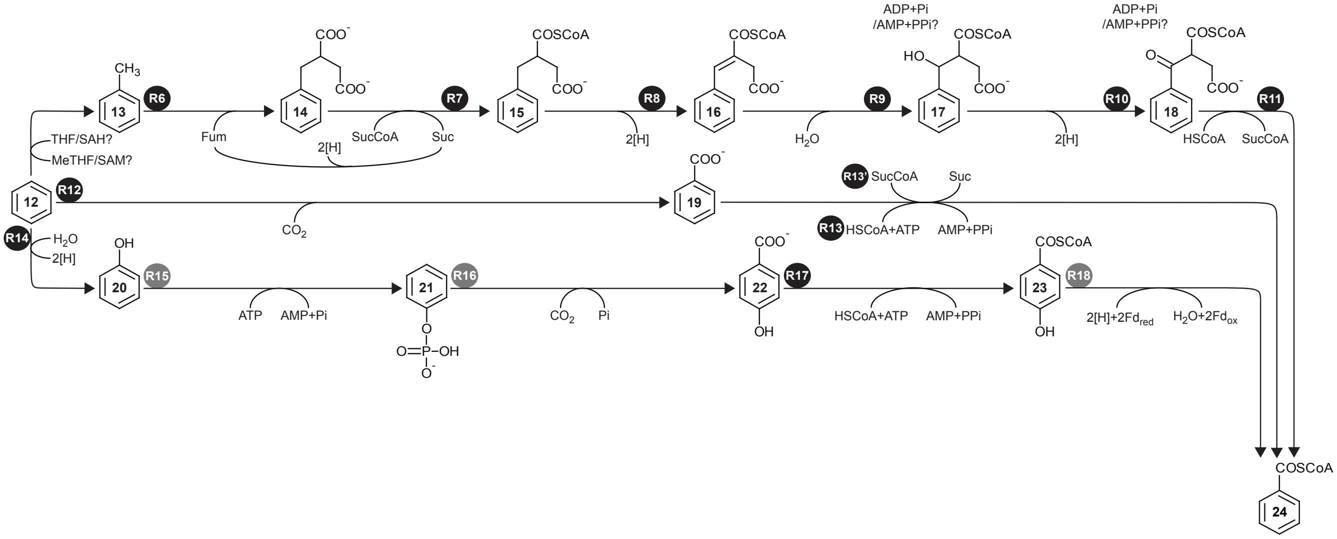


**Figure 12.** The anaerobic benzene peripheral degradation pathways up to the level of benzoyl CoA intermediates. See Supp. M&M section Section 1.13 for details.

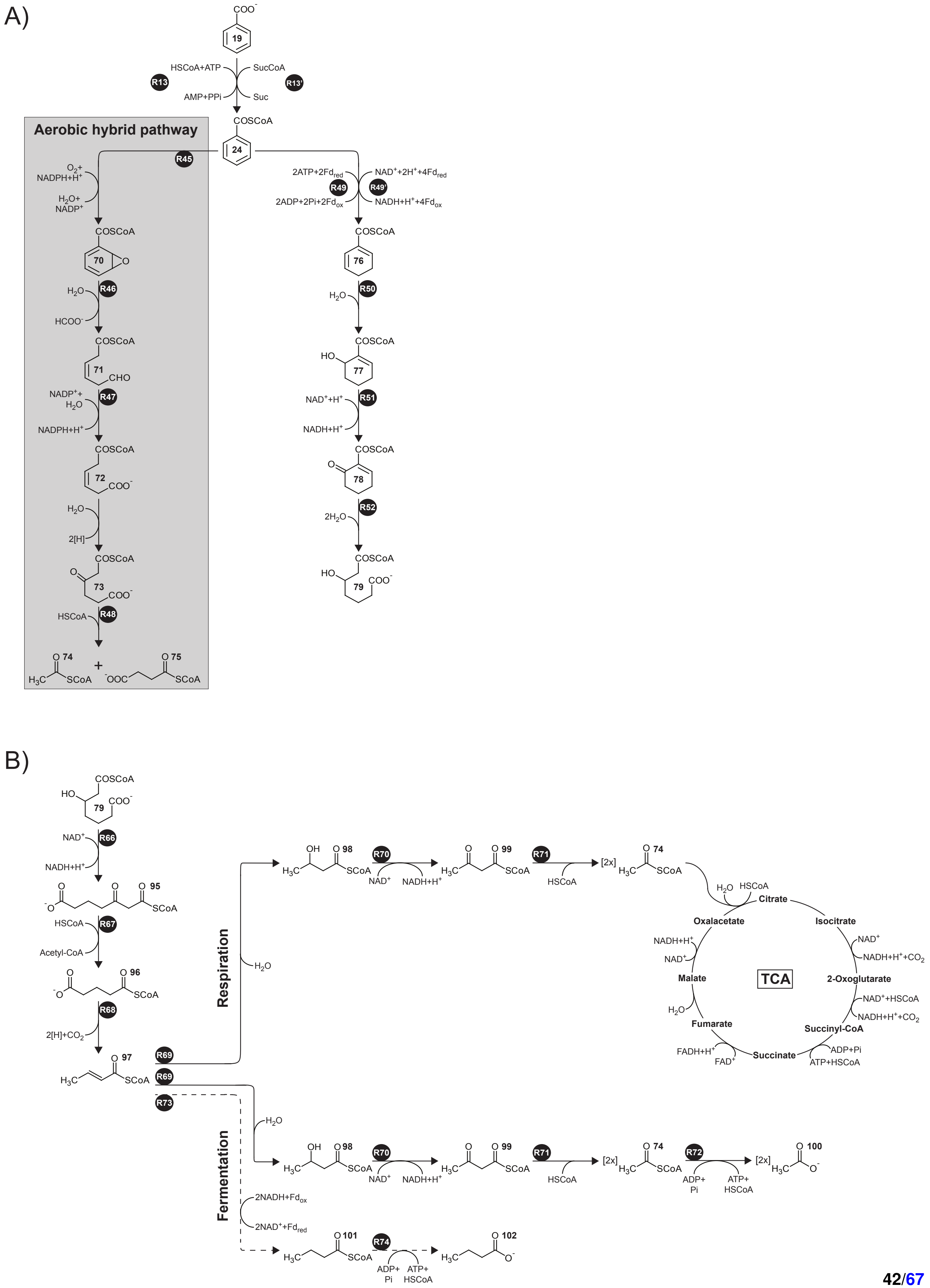




**Figure 13.** The anaerobic benzene central degradation pathways. See Supp. M&M section Section 1.13 for details.

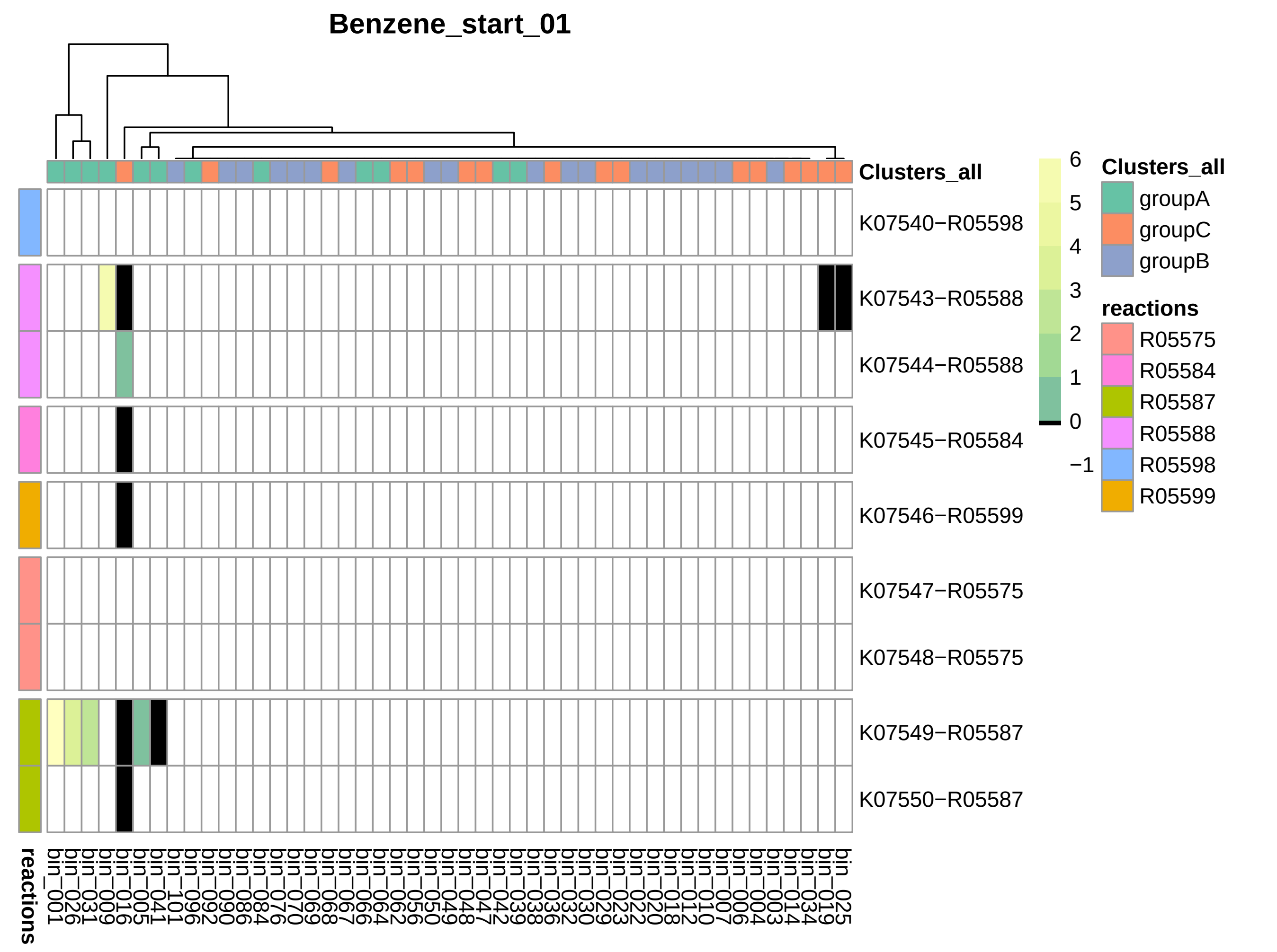


**Figure 14.** Heatmap represent opening of benzene ring and conversion to benzoyl-CoA through bezylsuccinate, with the presence and absence (black/white) of the KOs and color scale indicating the strength of transcription. This selection is the same with KEGGs module M00418

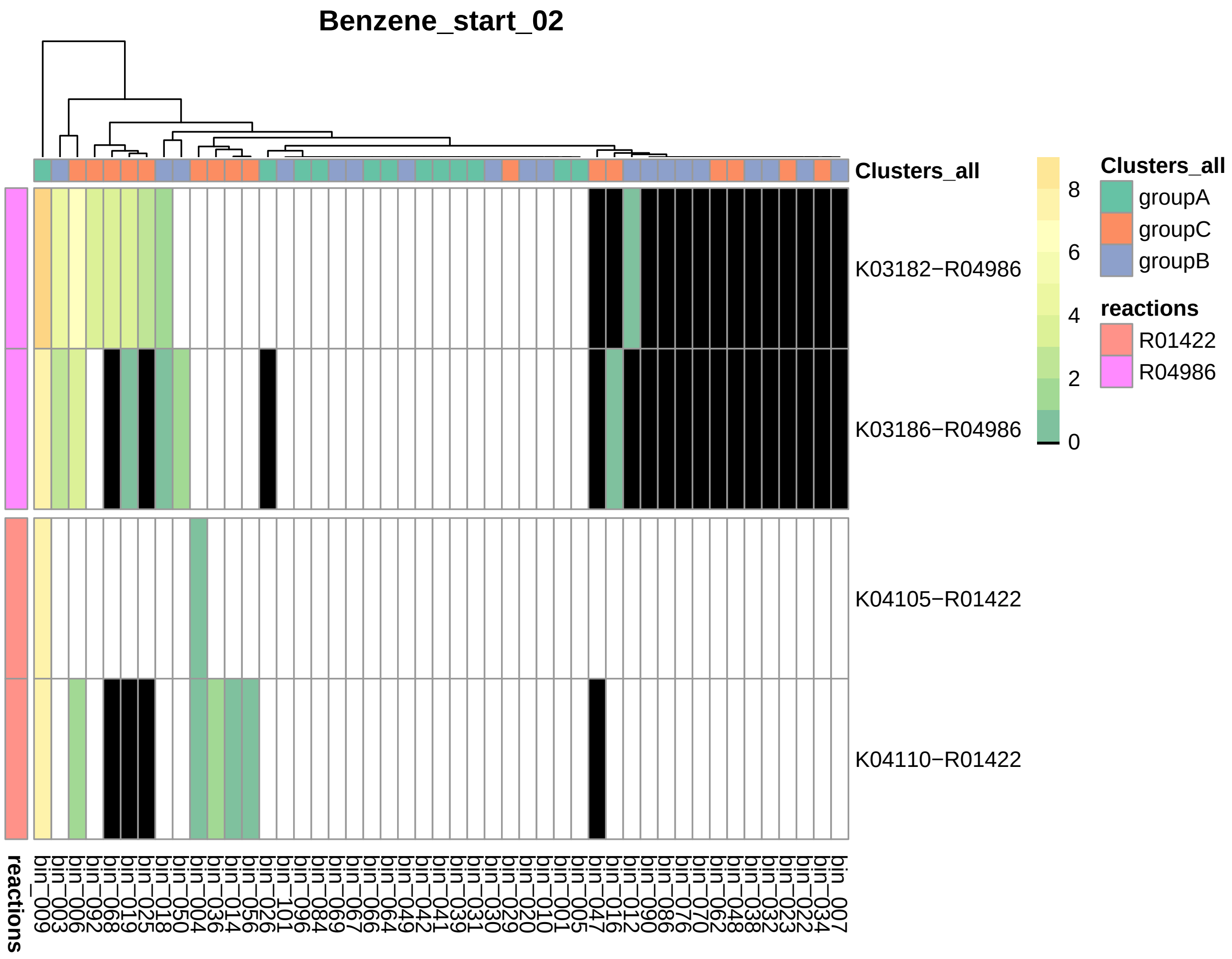


**Figure 15.** Heatmap represent opening of benzene ring and conversion to benzoyl-CoA through benzoate, with the presence and absence (black/white) of the KOs and color scale indicating the strength of transcription.

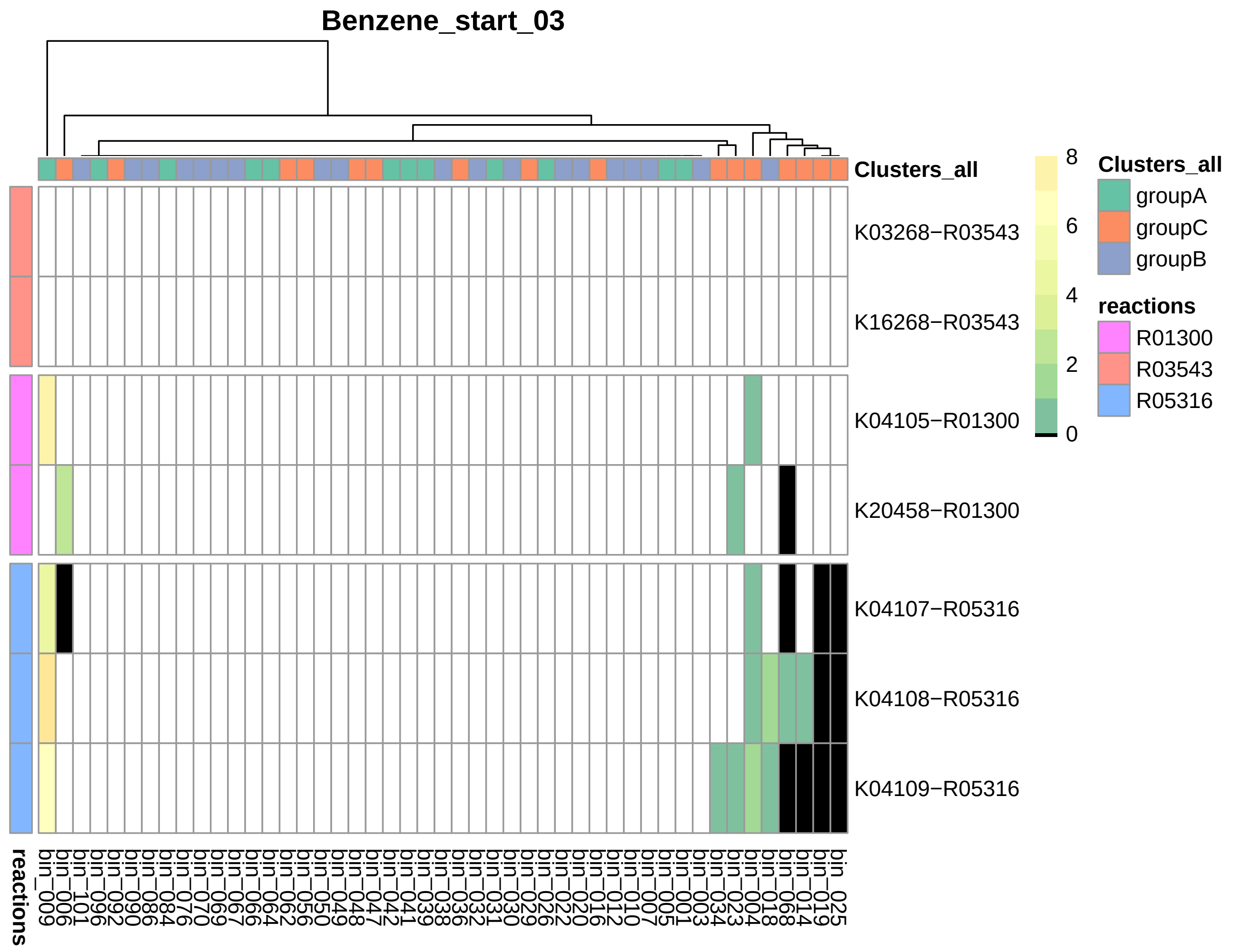


**Figure 16.** Heatmap represent opening of benzene ring and conversion to benzoyl-CoA through hydroxybenzene, with the presence and absence (black/white) of the KOs and color scale indicating the strength of transcription.

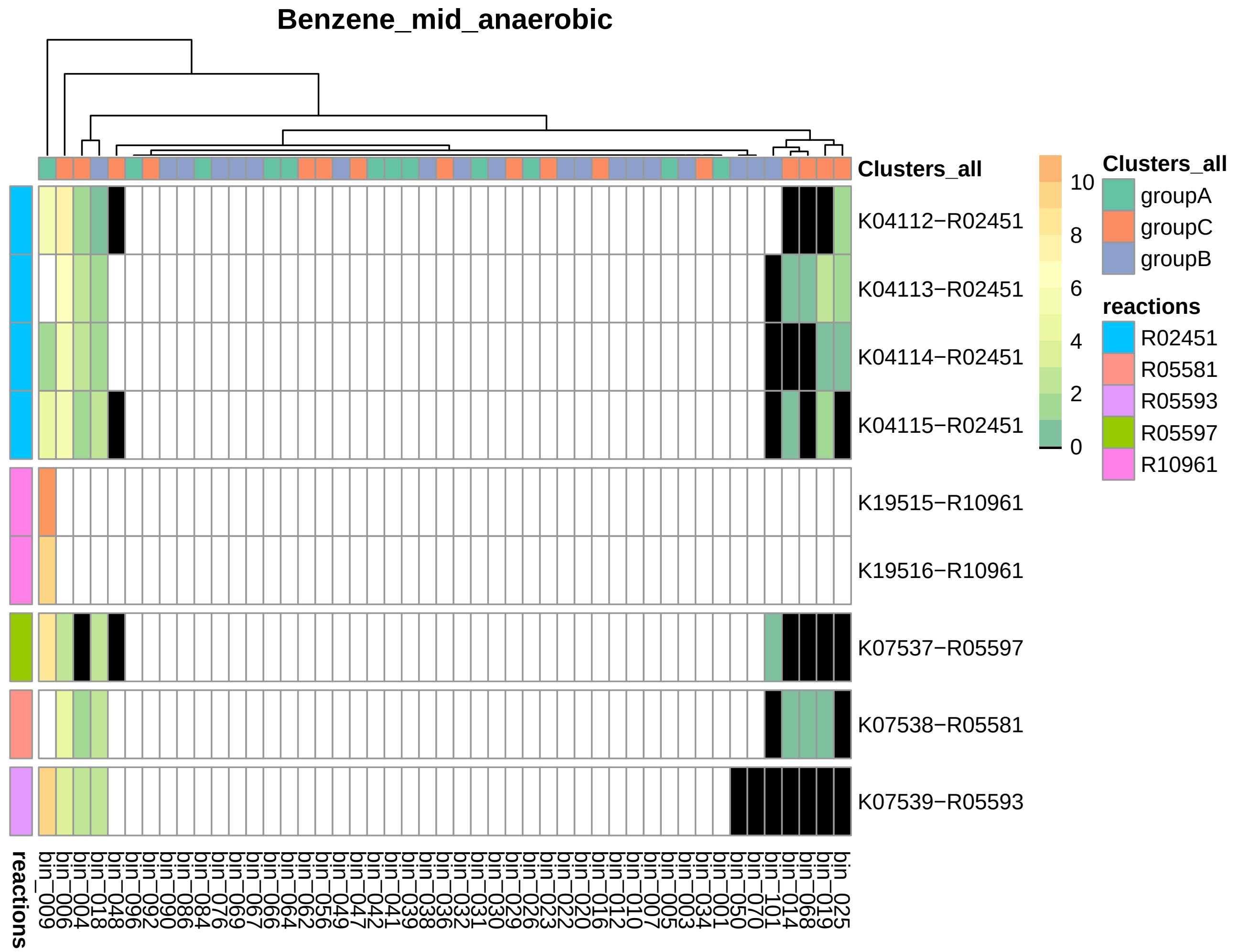


**Figure 17.** Heatmap represent anaerobic conversion of benzoyl-CoA to hydroxipimelyl-CoA, with the presence and absence (black/white) of the KOs and color scale indicating the strength of transcription. This selection is the same with KEGGs module M00541

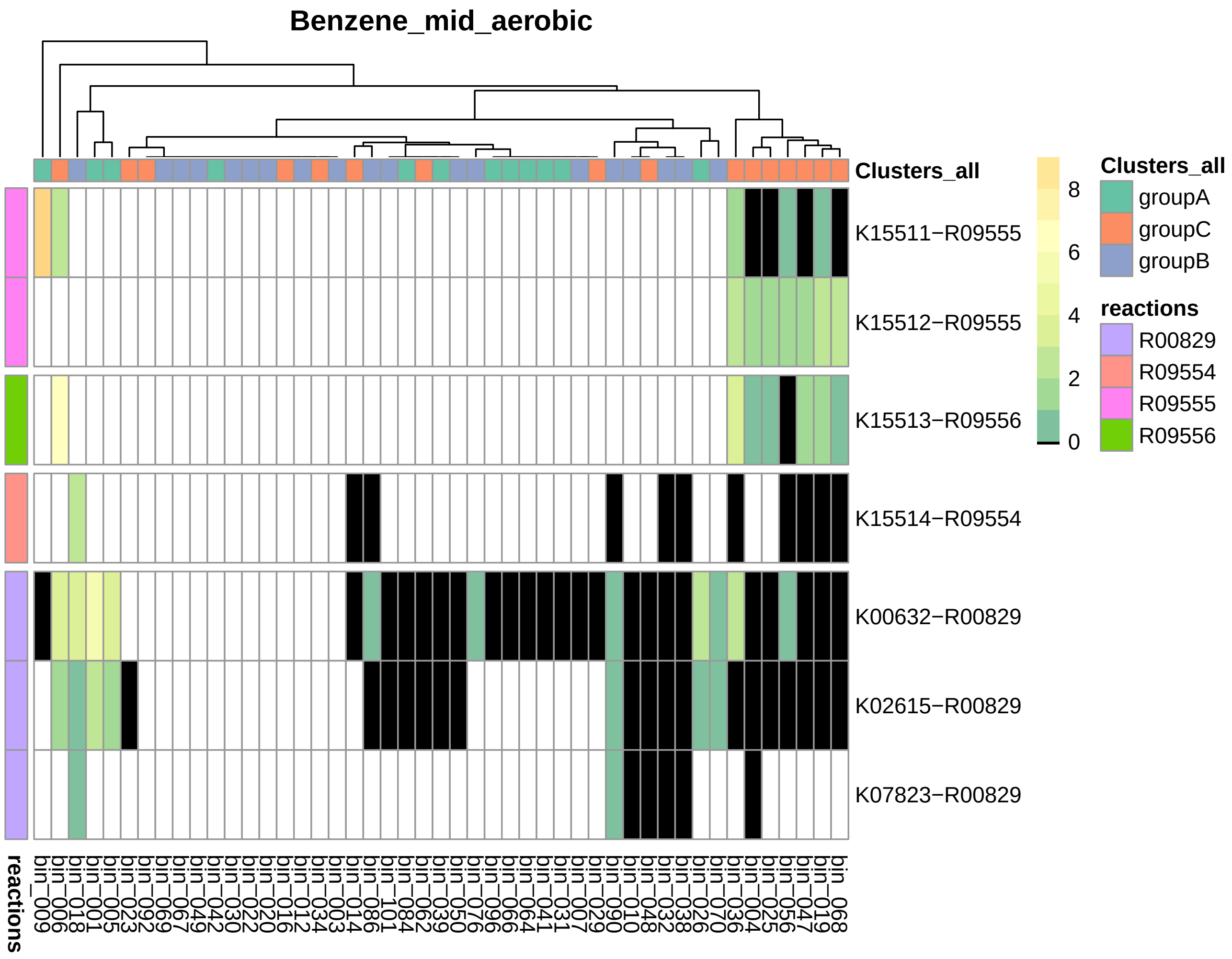


**Figure 18.** Heatmap represent semi-aerobic conversion of benzoyl-CoA to acetyl-CoA and succinyl-CoA, with the presence and absence (black/white) of the KOs and color scale indicating the strength of transcription.

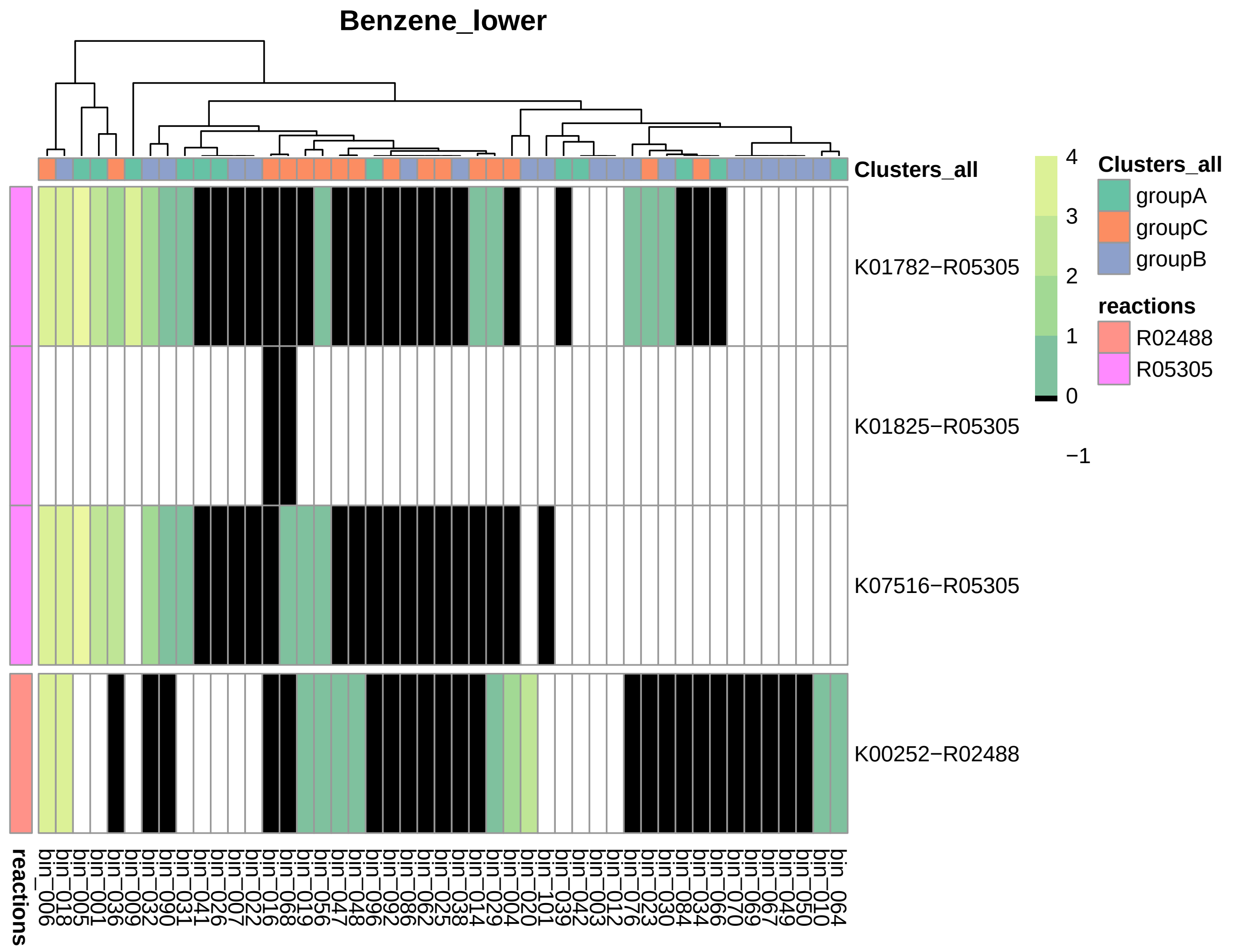


**Figure 19.** Heatmap represent conversion of hydroxipimelyl-CoA to crotonyl-CoA, with the presence and absence (black/white) of the KOs and color scale indicating the strength of transcription.

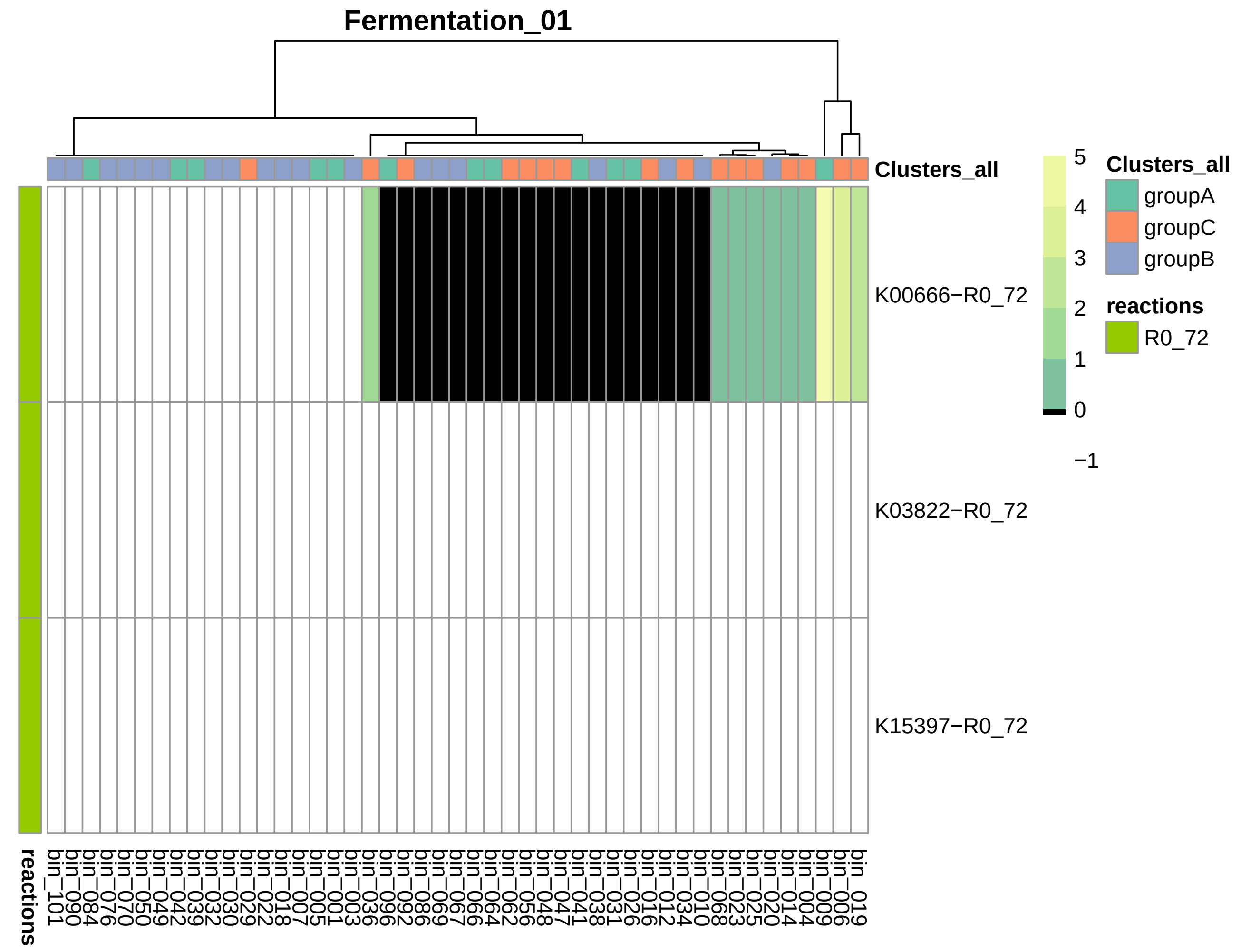


**Figure 20.** Heatmap on key KOs to identify fermatation of crotonyl-CoA to acetate, with the presence and absence (black/white) of the KOs and color scale indicating the strength of transcription.

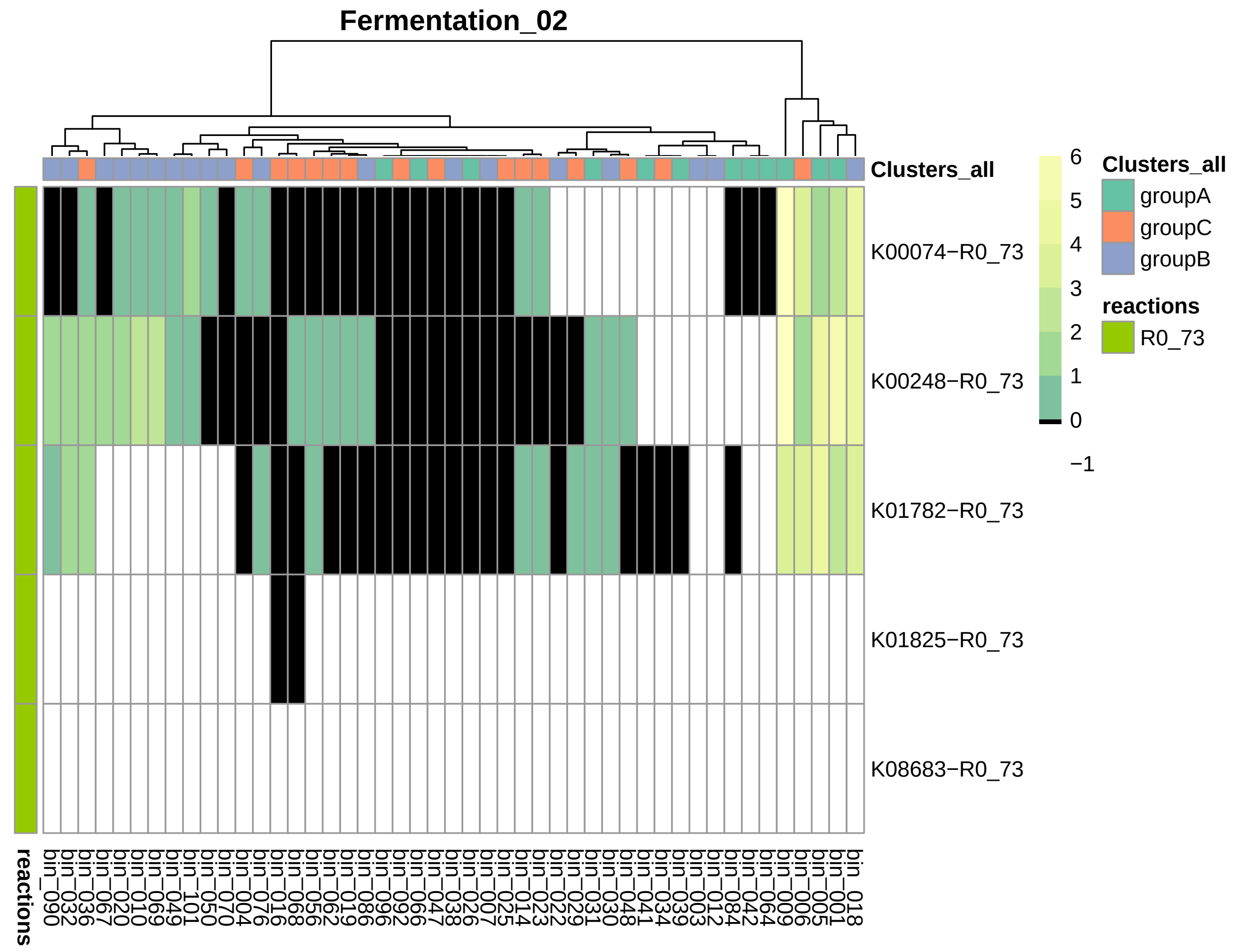


**Figure 21.** Heatmap of key KOs to identify fermentation of crotonyl-CoA to butyrate, with the presence and absence (black/white) of the KOs and color scale indicating the strength of transcription.

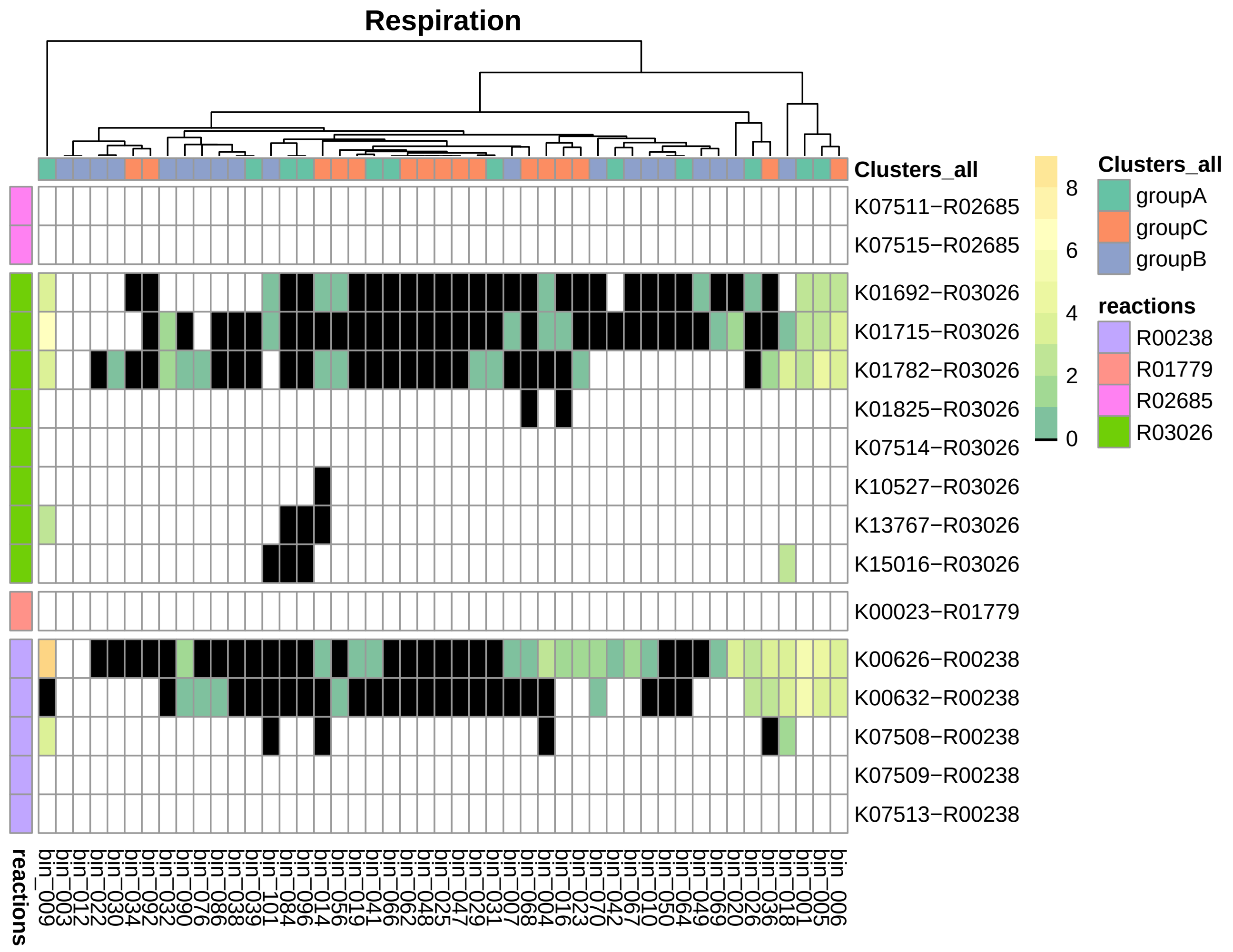


**Figure 22.** Heatmap of key KOs to identify fermentation of crotonyl-CoA to acetyl-CoA, with the presence and absence (black/white) of the KOs and color scale indicating the strength of transcription.

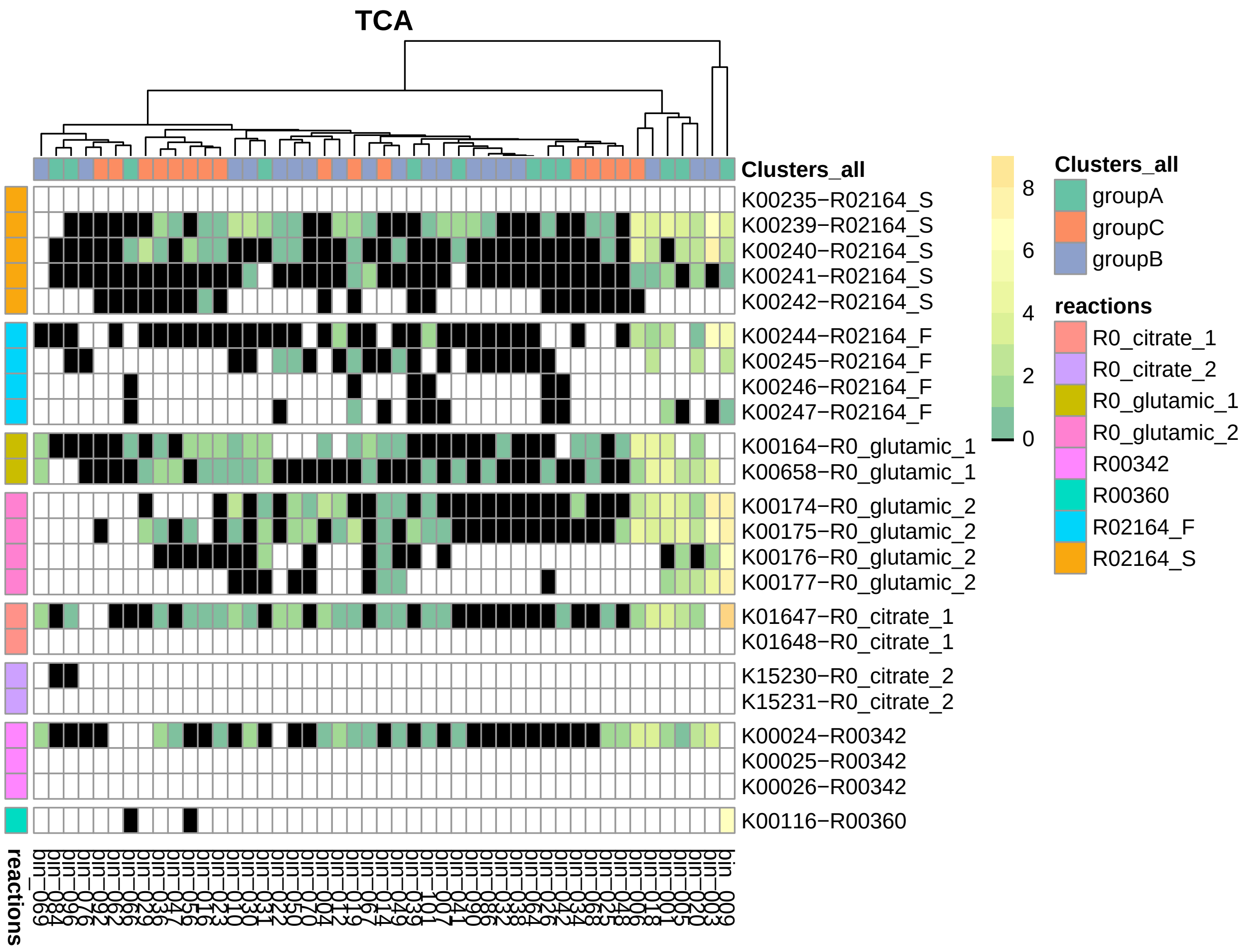


**Figure 23.** Heatmap of selected KOs to identify TCA, with the presence and absence (black/white) of the KOs and color scale indicating the strength of transcription.

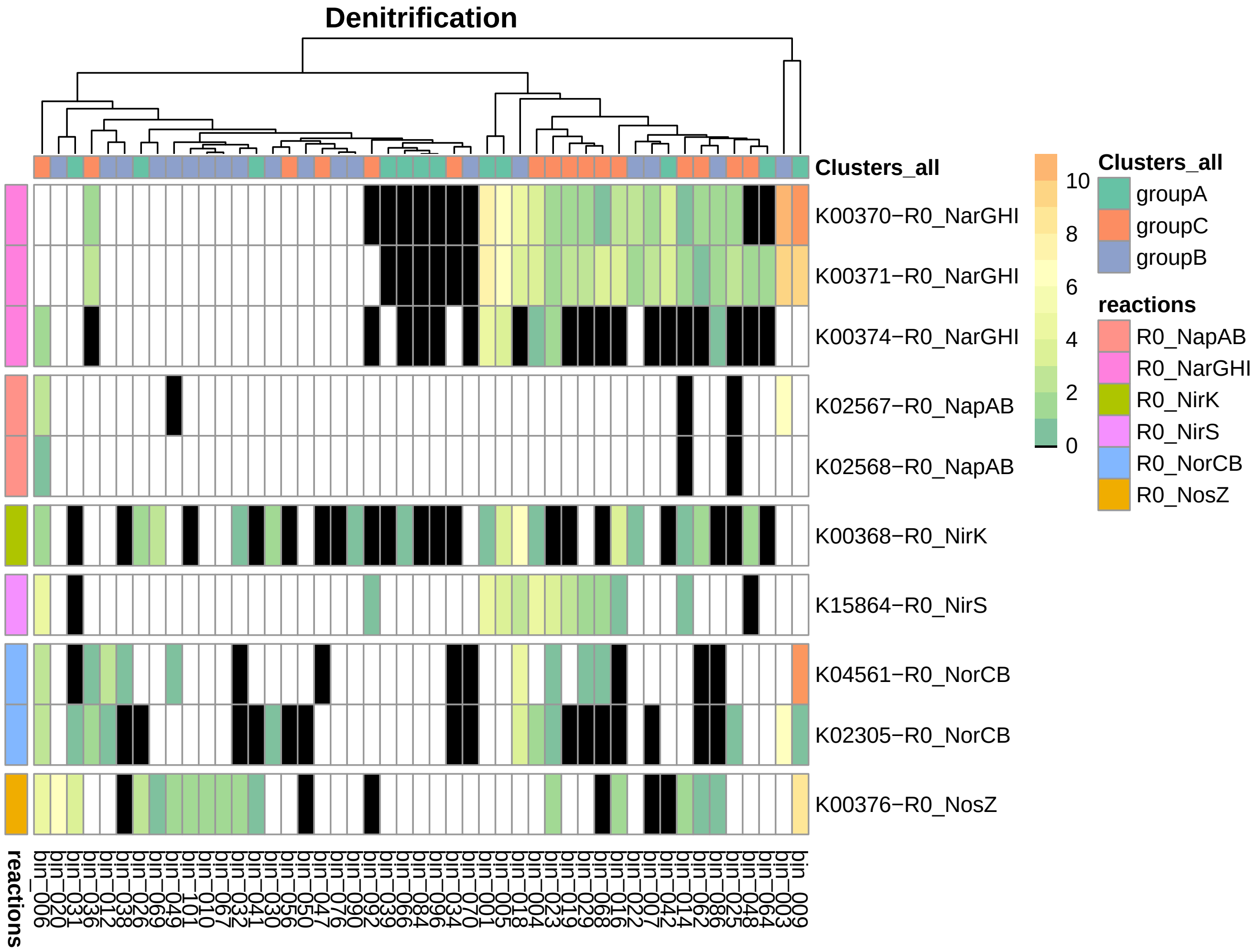


**Figure 24.** Heatmap of KOs to identify denitrification, with the presence and absence (black/white) of the KOs and color scale indicating the strength of transcription.

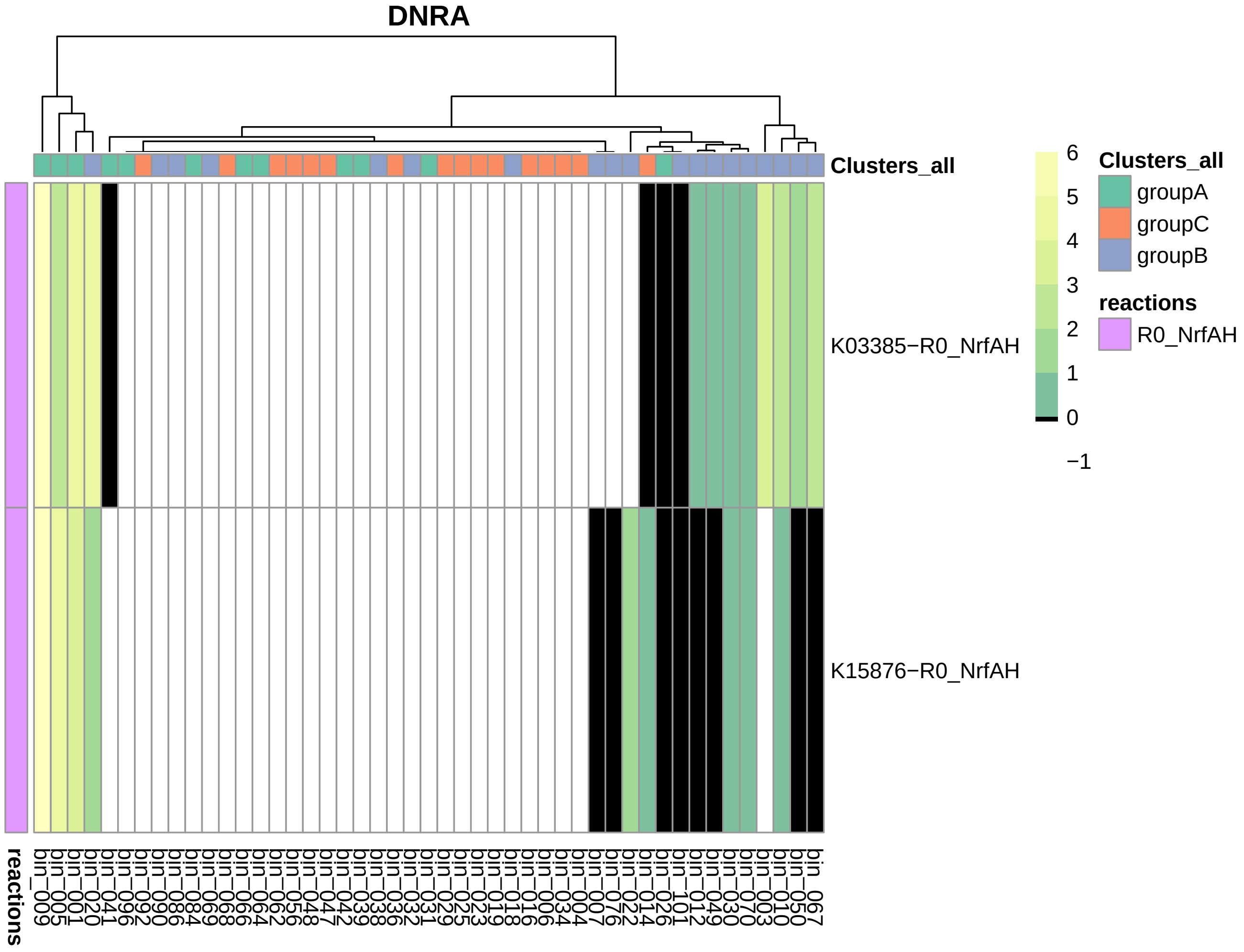


**Figure 25.** Heatmap of KOs to identify DNRA, with the presence and absence (black/white) of the KOs and color scale indicating the strength of transcription.

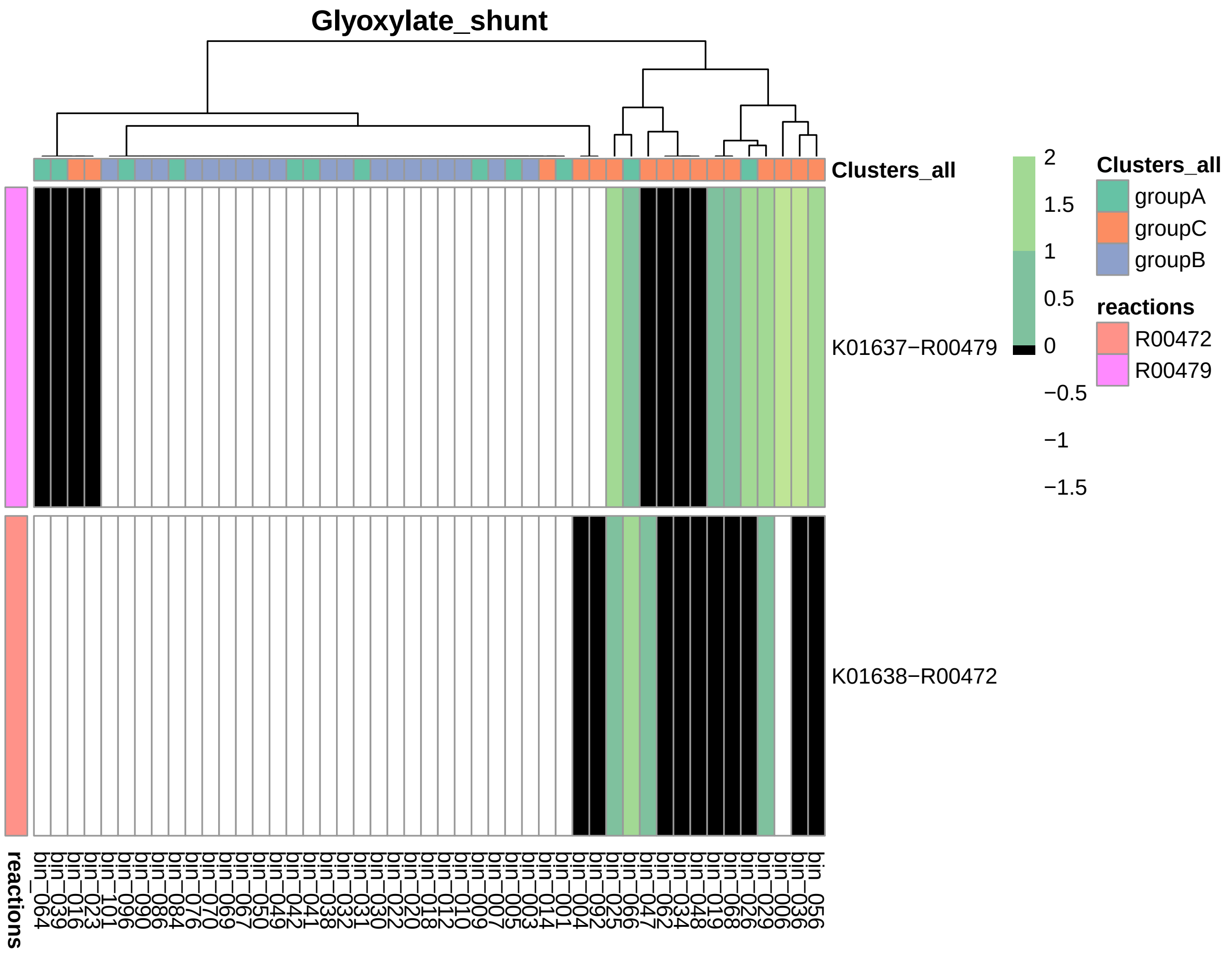


**Figure 26.** Heatmap of KOs to identify glyoxylate shunt, with the presence and absence (black/white) of the KOs and color scale indicating the strength of transcription.

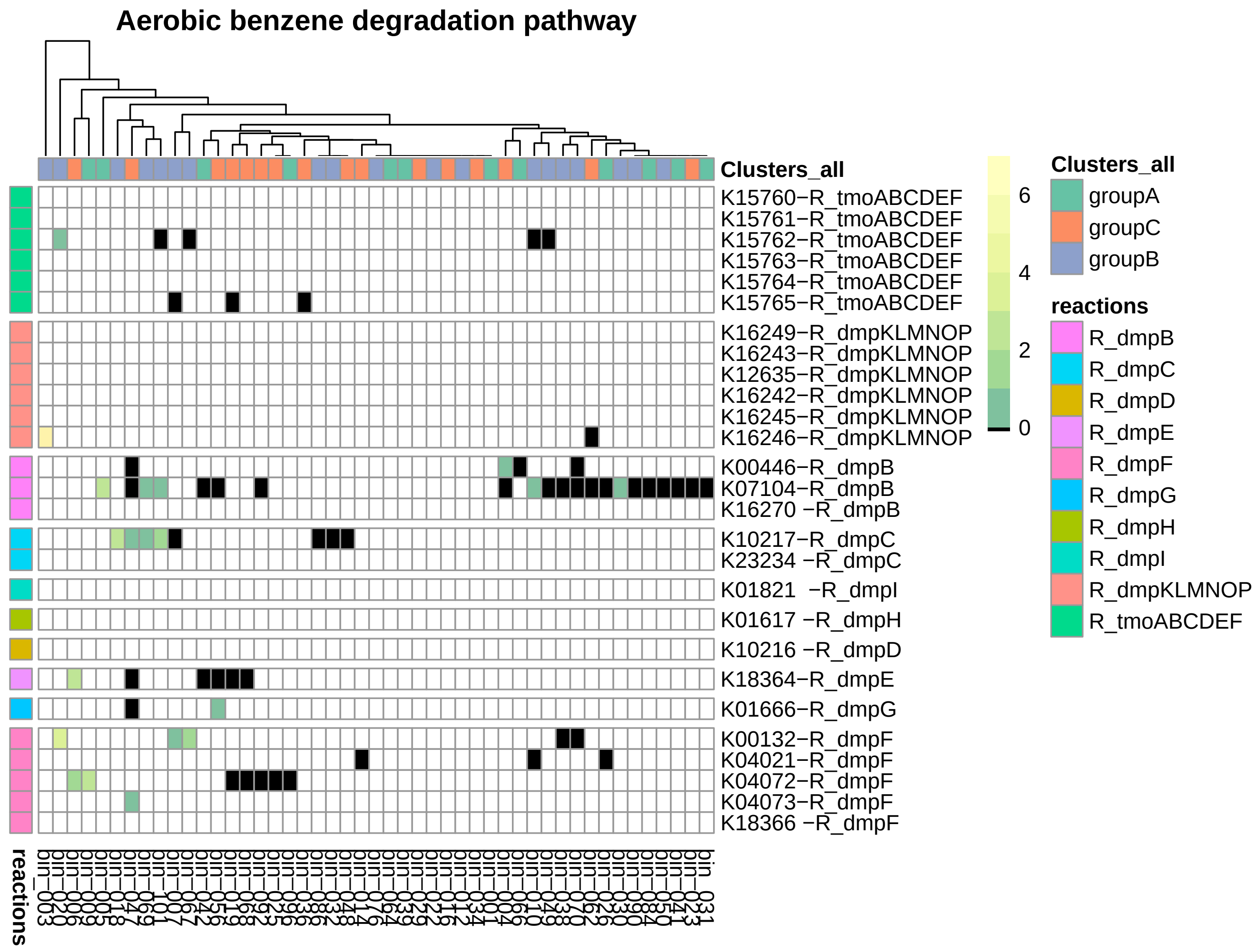


**Figure 27.** Heatmap of KOs to identify aerobic benzene degradation, with the presence and absence (black/white) of the KOs and color scale indicating the strength of transcription.

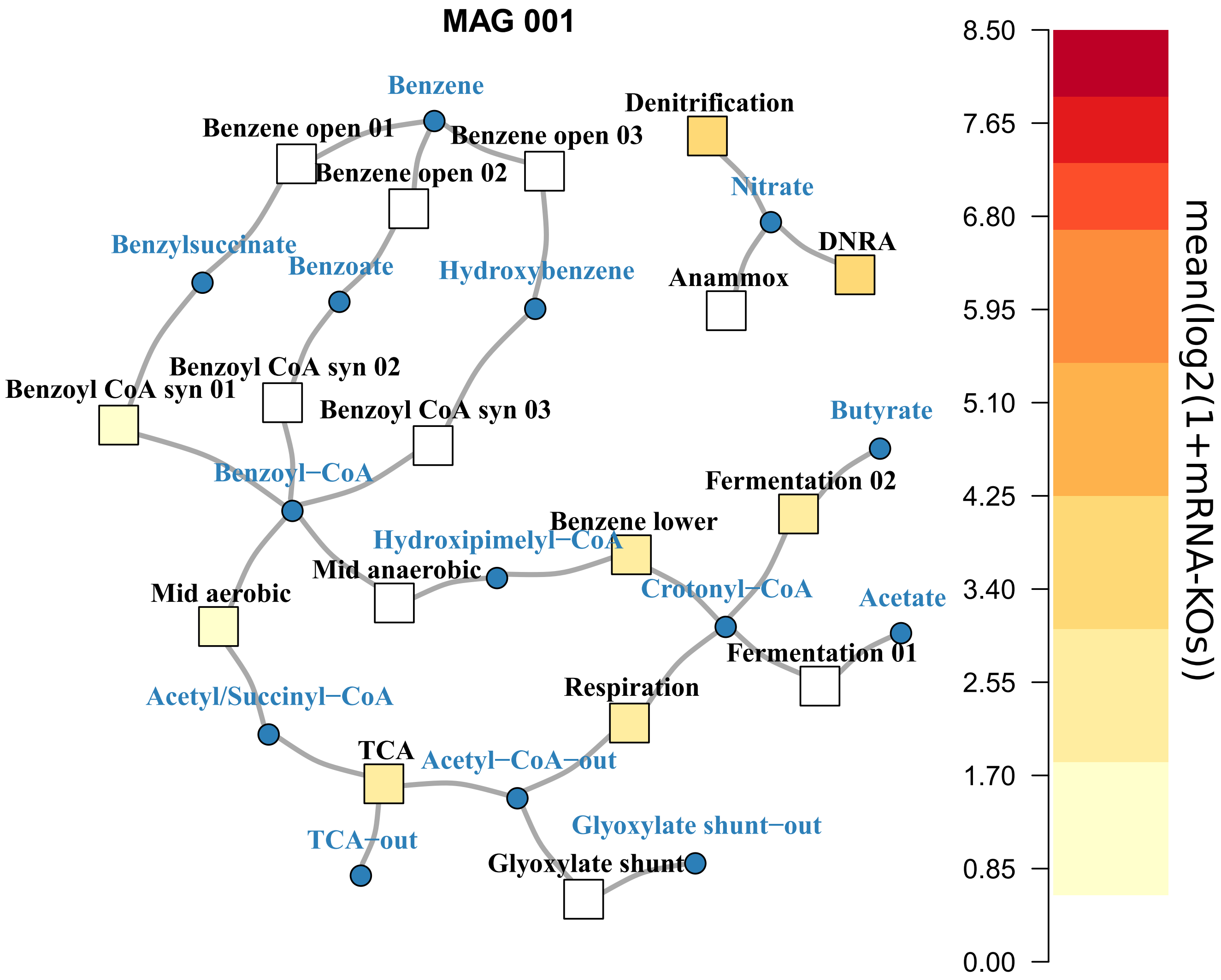


**Figure 28.** The qualitative transcription core metabolic graph of anaerobic benzene degradation of dominant MAG 1 (P: Chloroflexi, F: *Anaerolineaceae*)

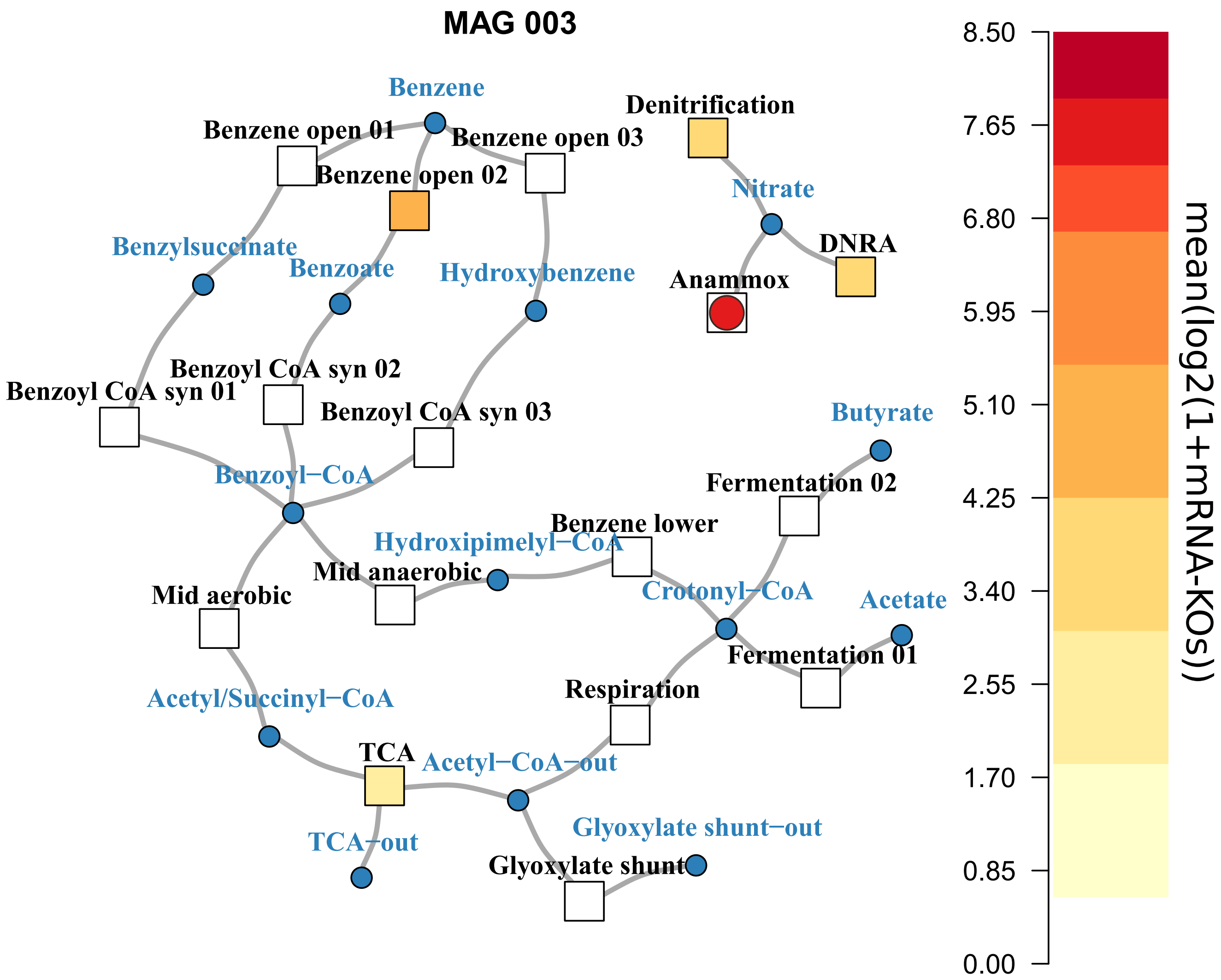


**Figure 29.** The qualitative transcription core metabolic graph of anaerobic benzene degradation of dominant MAG 3 (P: Planctomycetes, G: *Candidatus* Kuenenia). The MAG has the key genes for anaerobic ammonium oxidation (anammox), hydrazine synthase and dehydrogenase. The one encoding the catalytic subunit of the synthase is highly expressed but had to be assigned to MAG 0 as the contig where it is on did not meet the criteria for placement in MAG 3

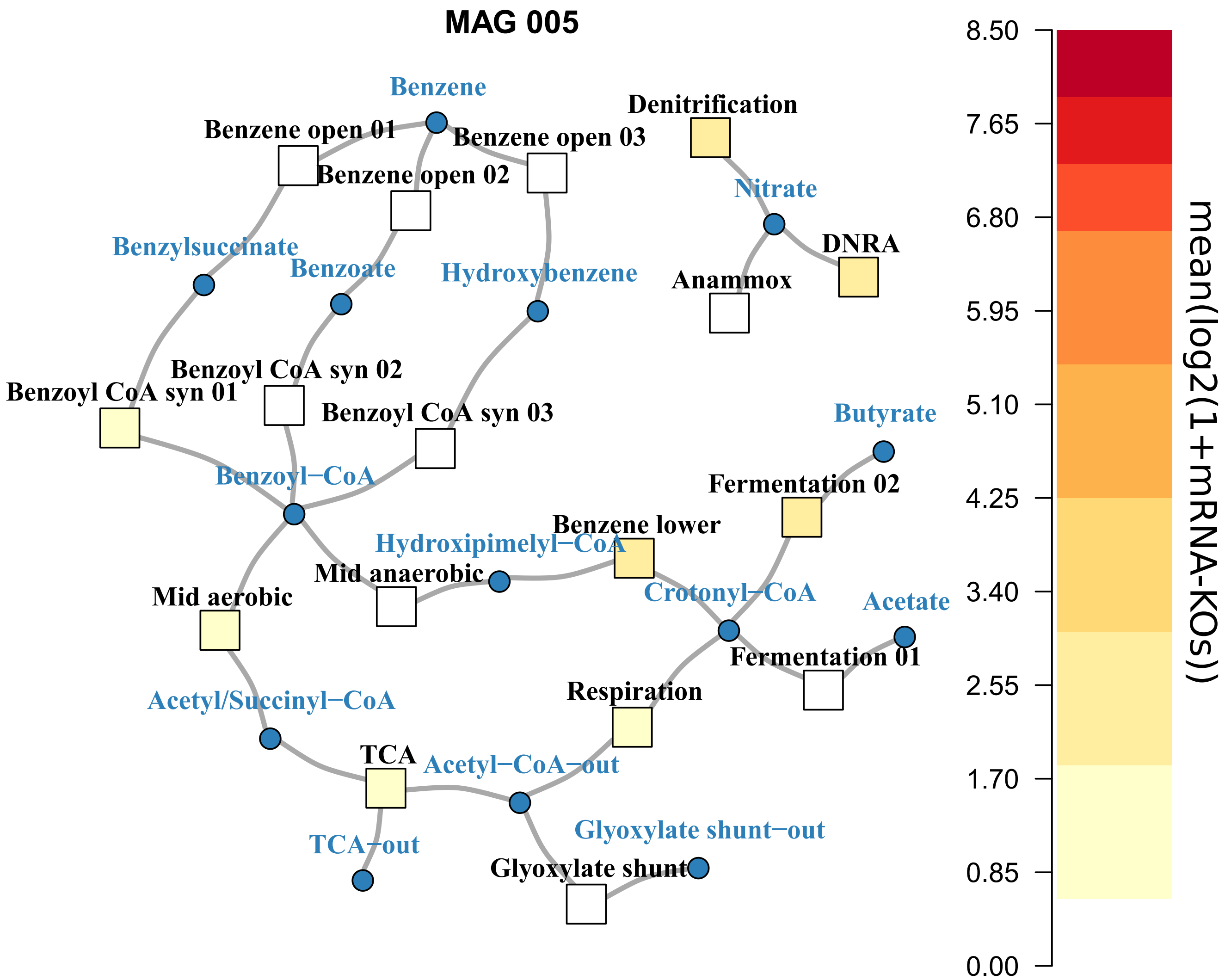


**Figure 30.** The qualitative transcription core metabolic graph of anaerobic benzene degradation of dominant MAG 5 (P: Chloroflexi, F: *Anaerolineaceae*)

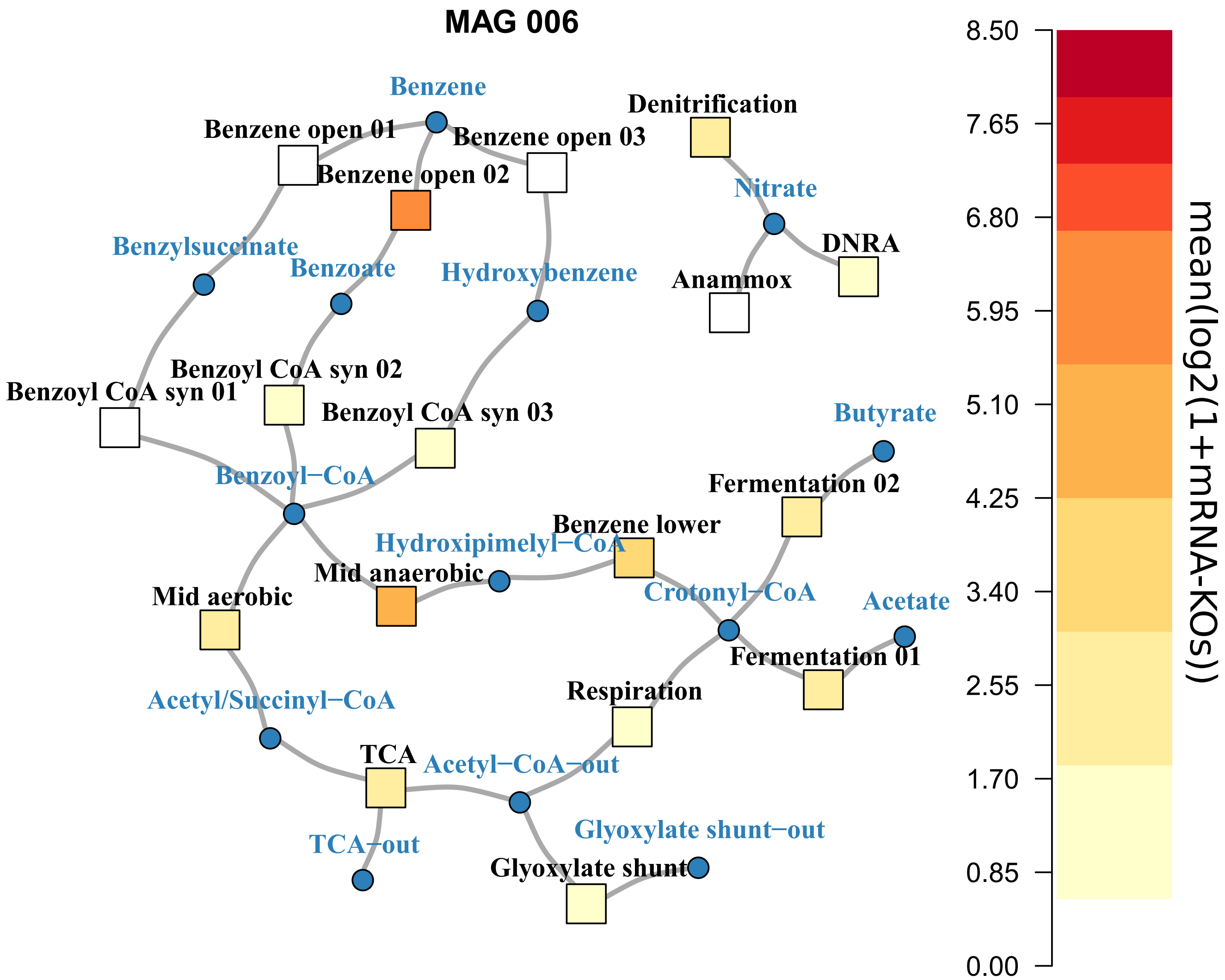


**Figure 31.** The qualitative transcription core metabolic graph of anaerobic benzene degradation of dominant MAG 6 (P: Proteobacteria, F: *Rhodocyclaceae*)

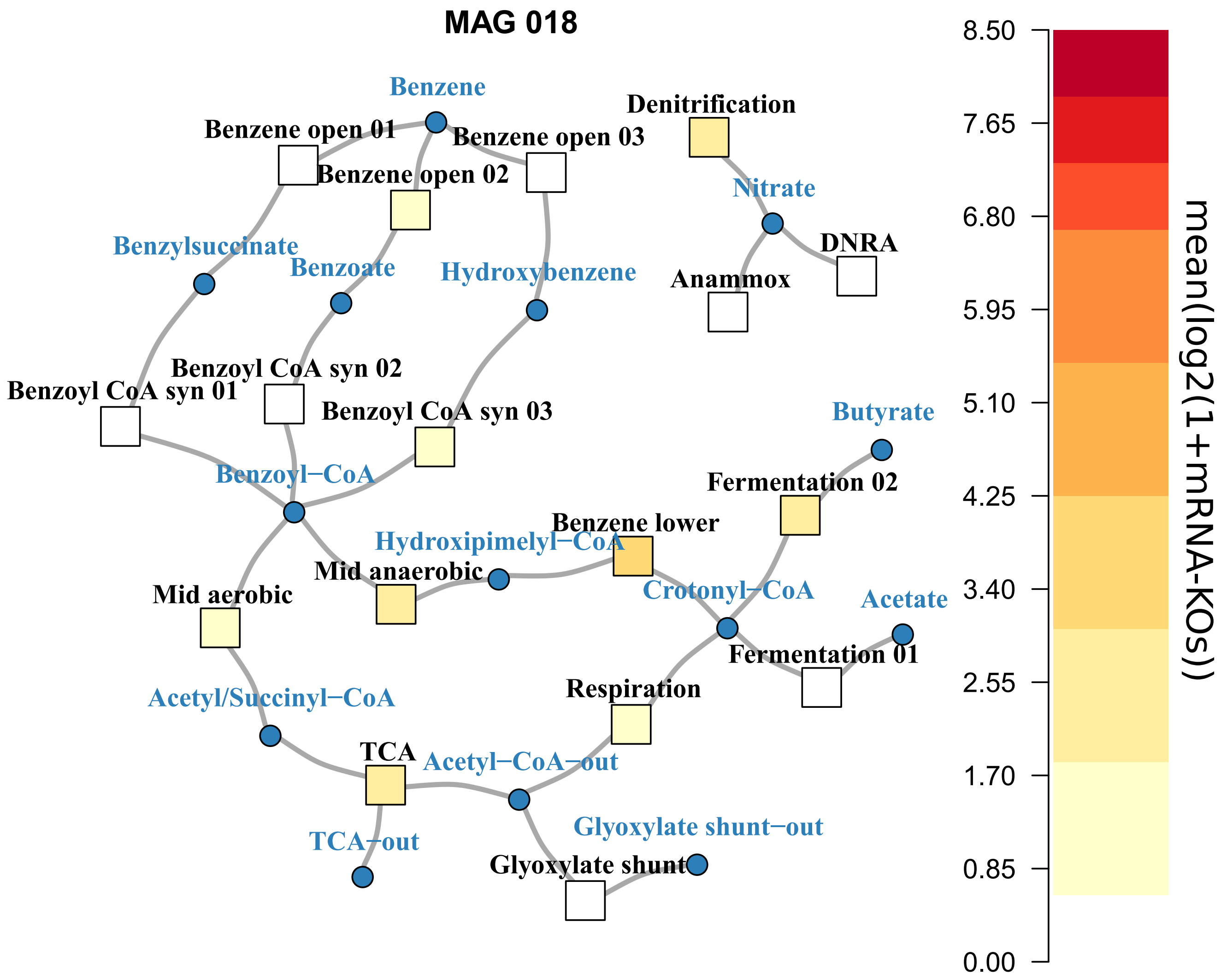


**Figure 32.** The qualitative transcription core metabolic graph of anaerobic benzene degradation of dominant MAG 18 (P: Bacteroidetes, F: *Cryomorphaceae*)

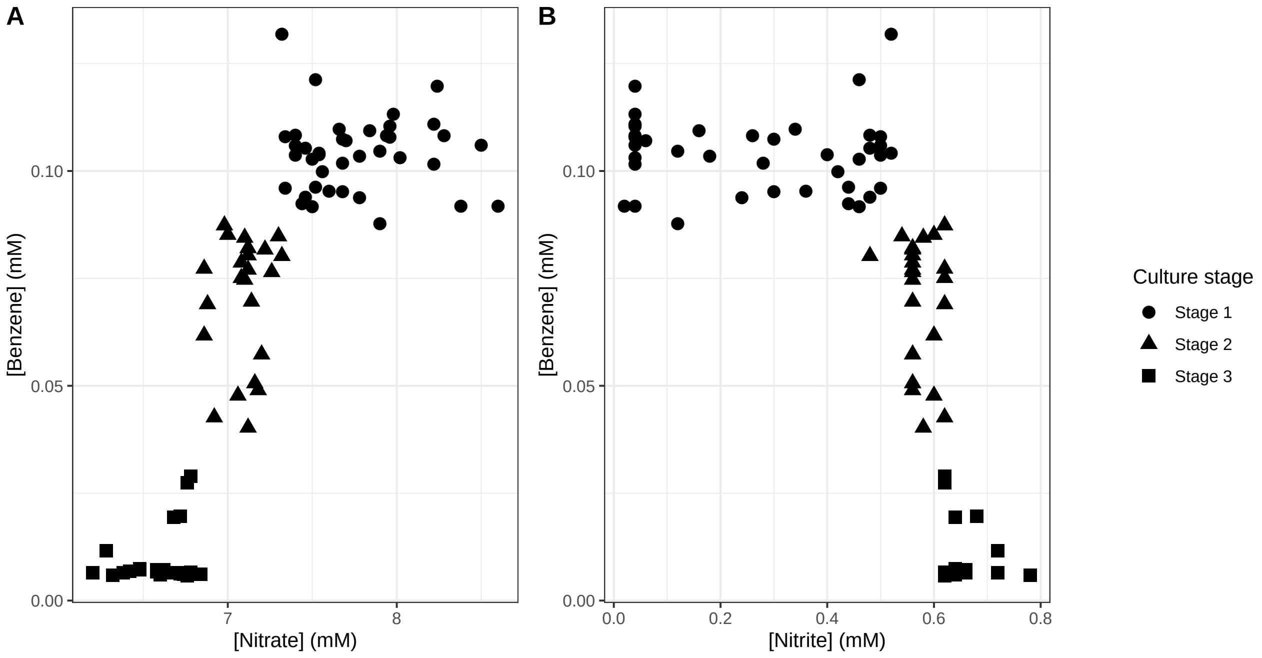


**Figure 33.** Dependency between benzene and nitrate (A) or nitrite (B) concentrations in all cultures. Note that in Stage 1 cultures no benzene has been consumed.

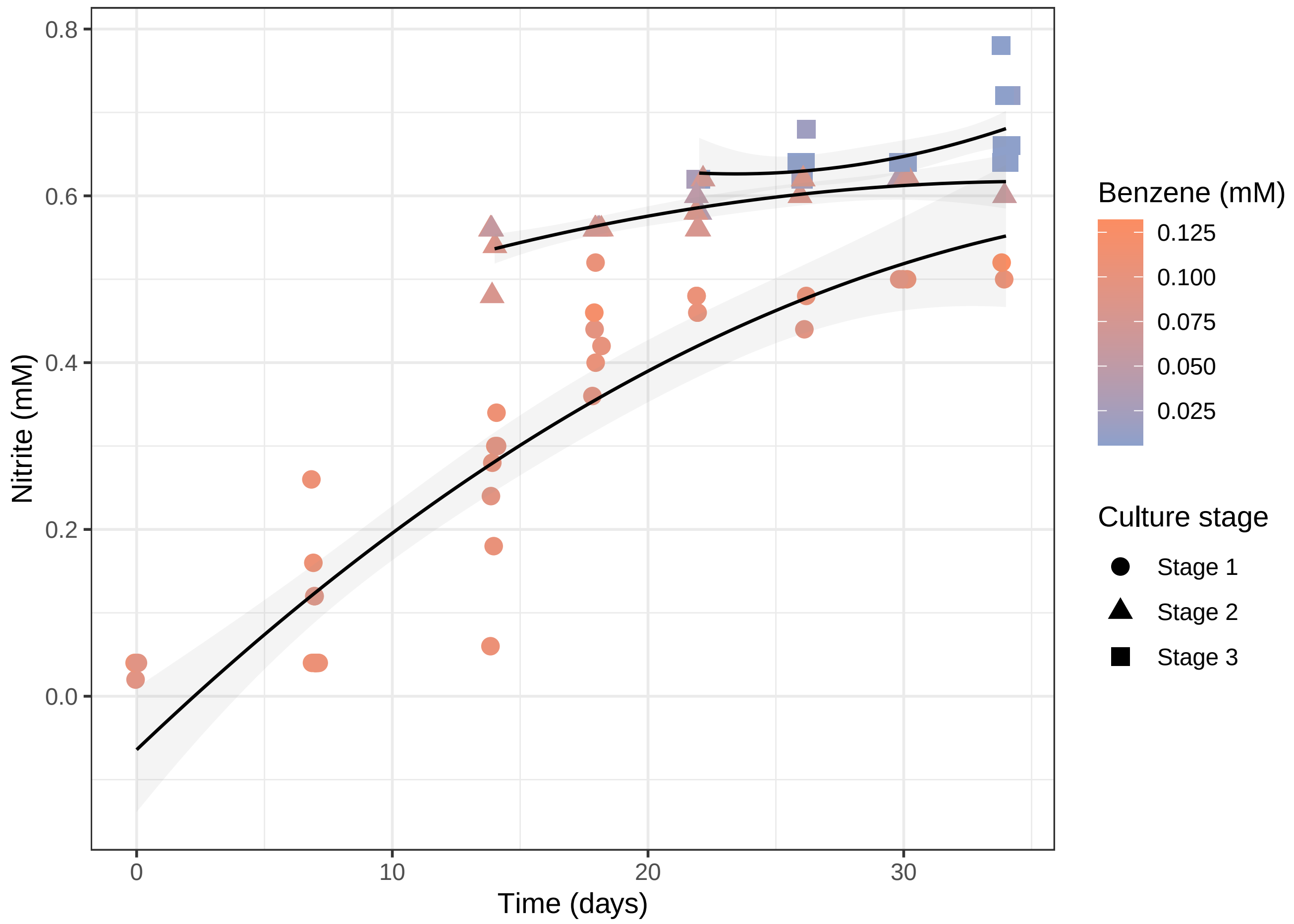


**Figure 34.** Relationship between nitrite and benzene concentrations in all cultures over time. Fits are second order (quadratic) polynomial. Note that the rate of nitrite production is higher in stage 1, decreases in stage 2 while benzene consumption starts and slowly increasing again when benzene is fully consumed at stage 3.

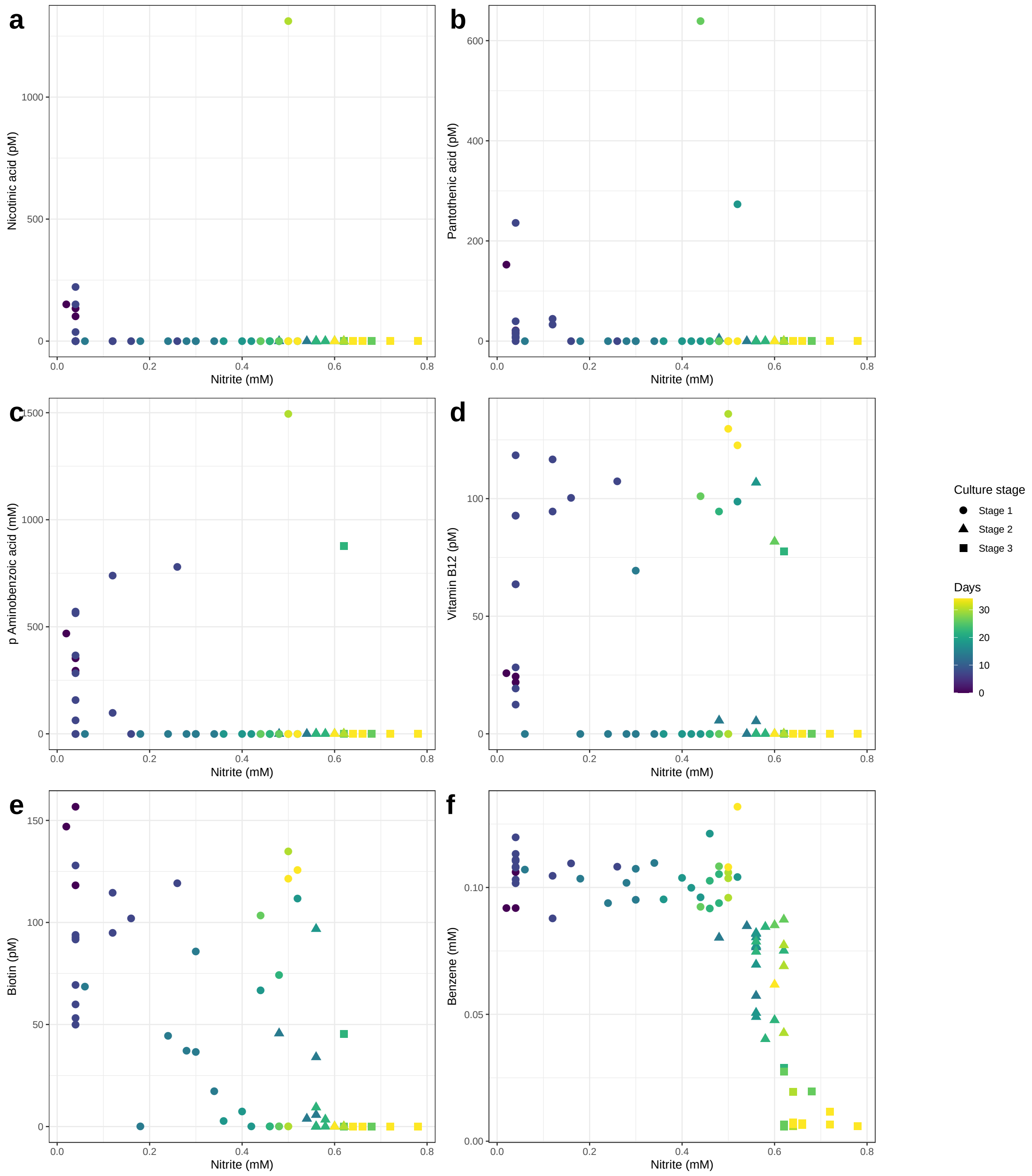


**Figure 35.** Relation between (a) nicotinic acid, (b) pantothenic acid, (c) para-aminobenzoic acid, (d) vitamin B12, (e) biotin and (f) benzene with nitrite levels in all cultures overtime.

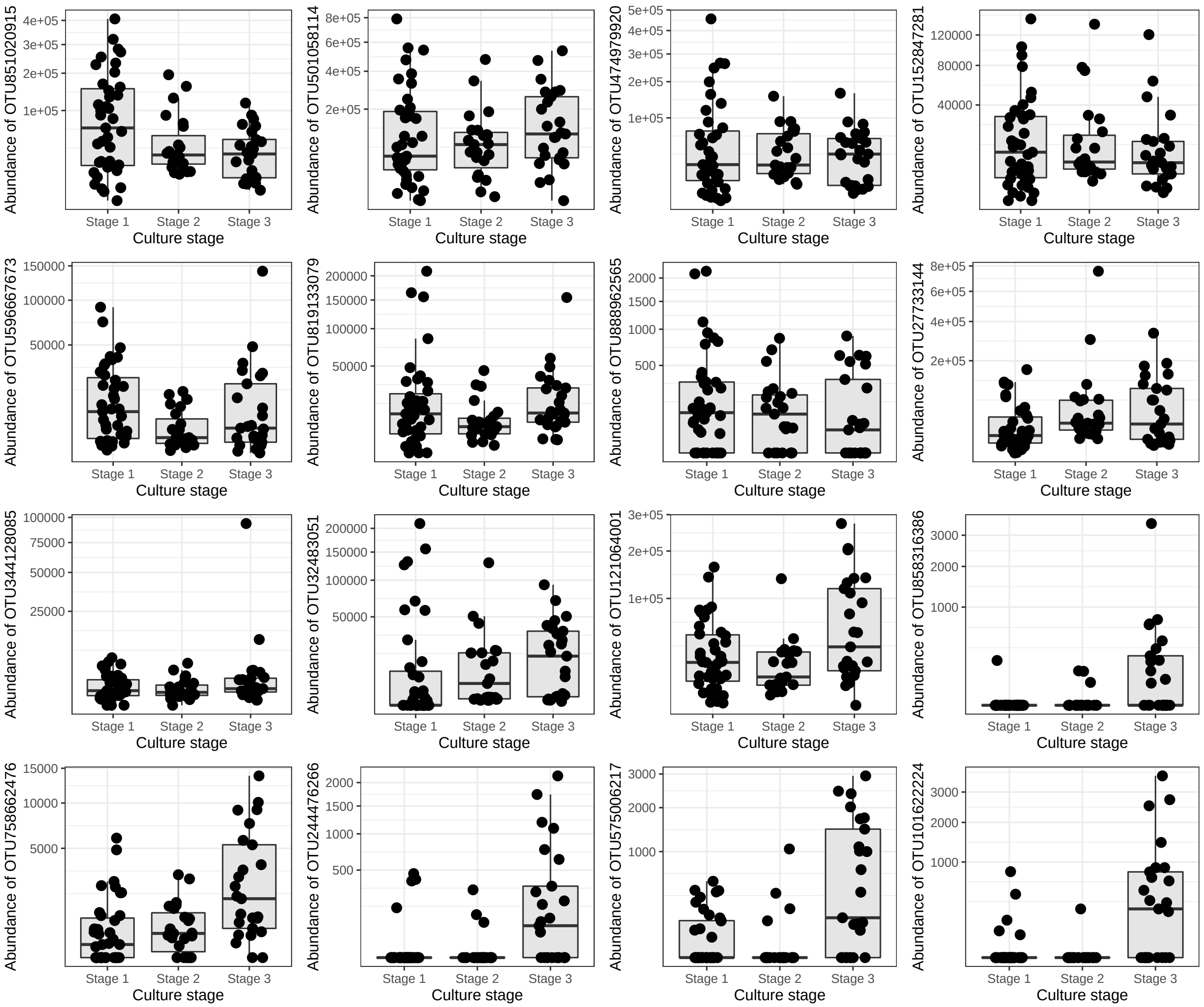


**Figure 36.** Relationship between culture stage and cell abundance of top significant OTUs based on random forest variable importance and PERMANOVA analysis. For taxonomy see Table 9

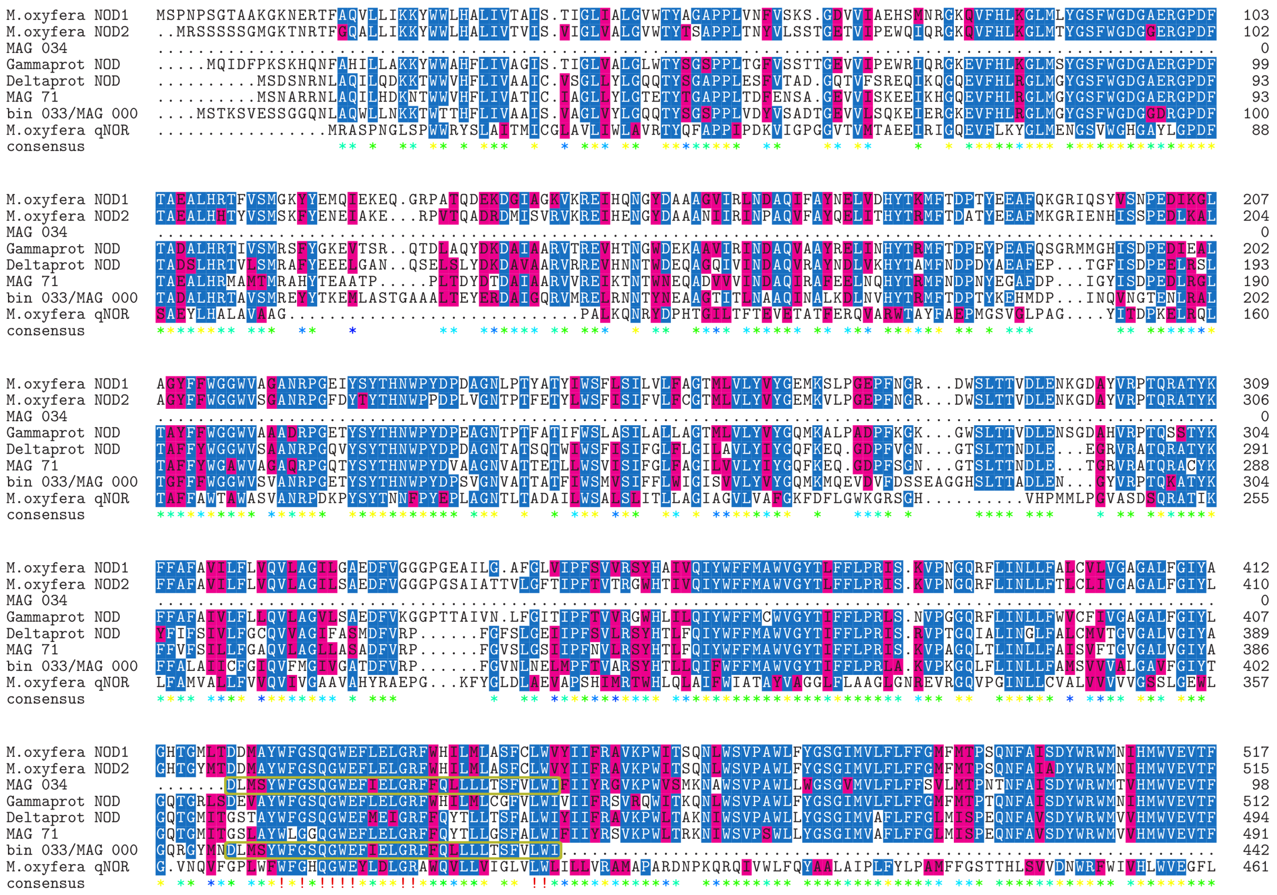

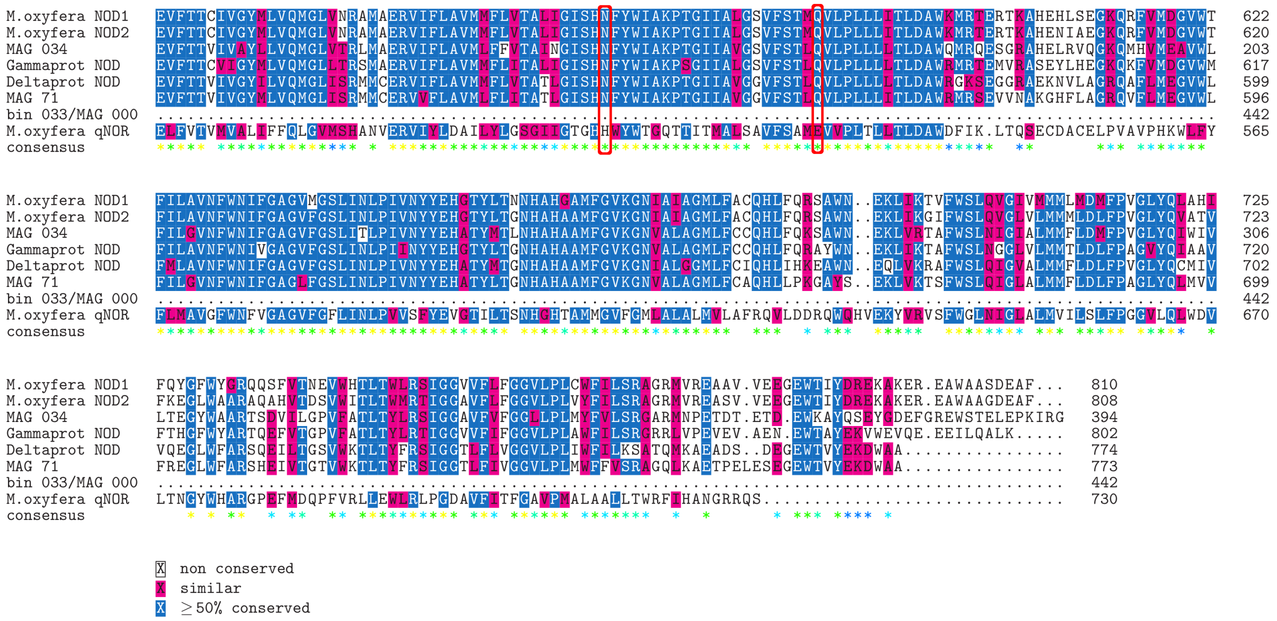


**Figure 37.** Multiple sequence alignment result between predicted NODs from MAG 34,MAG 33/MAG 0 and 71 with *M. oxyfera* NOD1, NOD2, $\gamma$-protebacteria NOD, $\delta$-protebacteria NOD and qNOR (GenBank ids: CBE69502.1, CBE69496.1, TAJ95298.1, MAG34007.1 and CBE68939.1 respectively). The analysis were performed with CLUSTAL 2.1 algorithm which used as default setting in msa R-package [8]. Note that the NOD genes from MAG34 and MAG 33/MAG0 is likely to be fragments of a complete version of the gene as the following amino acid sequence is overlap between the two "DLMSYWFGSQGWEFIELGRFFQLLLLTSFVLWI" - indicated with horizontal yellow rectangle. The conserved H and E in qNor enzymes and displacement by N and Q respectively in NODs are marked with a red rectangle. [21].
